# Potential future climate change effects on global reptile distributions and diversity

**DOI:** 10.1101/2022.05.07.490295

**Authors:** Matthias F. Biber, Alke Voskamp, Christian Hof

## Abstract

**Aim:** Until recently, complete information on global reptile distributions has not been widely available. Here, we provide the first comprehensive climate impact assessment for reptiles on a global scale.

**Location:** Global, excluding Antarctica

**Time period:** 1995, 2050, 2080

**Major taxa studied:** Reptiles

**Methods:** We modelled the distribution of 6,296 reptile species and assessed potential global as well as realm-specific changes in species richness, the change in global species richness across climate space, and species-specific changes in range extent, overlap and position under future climate change. To assess the future climatic impact on 3,768 range-restricted species, which could not be modelled, we compared the future change in climatic conditions between both modelled and non-modelled species.

**Results:** Reptile richness was projected to decline significantly over time, globally but also for most zoogeographic realms, with the greatest decrease in Brazil, Australia and South Africa. Species richness was highest in warm and moist regions, with these regions being projected to shift further towards climate extremes in the future. Range extents were projected to decline considerably in the future, with a low overlap between current and future ranges. Shifts in range centroids differed among realms and taxa, with a dominating global poleward shift. Non-modelled species were significantly stronger affected by projected climatic changes than modelled species.

**Main conclusions:** With ongoing future climate change, reptile richness is likely to decrease significantly across most parts of the world. This effect as well as considerable impacts on species’ range extent, overlap, and position were visible across lizards, snakes and turtles alike. Together with other anthropogenic impacts, such as habitat loss and harvesting of species, this is a cause for concern. Given the historical lack of global reptile distributions, this calls for a reassessment of global reptile conservation efforts, with a specific focus on anticipated future climate change.

## Introduction

Emissions from anthropogenic activities have led to an increase in global surface temperature of around 1°C in the last 100 years. This has already led to changes in the weather and the occurrence of climate extremes in every region across the globe (IPCC, 2021). Unless emissions are vastly reduced in the coming decades, global warming will continue and exceed 1.5°-2°C compared to pre-industrial levels by the end of the 21st century (IPCC, 2021).

Climate change has already had adverse effects on biodiversity and ecosystem functioning and these effects are likely to worsen as warming continues in the future (IPBES, 2019; IPCC, 2022). Climate change impacts on ecological processes at scales ranging from genes to entire ecosystems can affect organisms, populations, or entire communities, and vary between physiological, morphological, phenological and distributional shifts (Bellard *et al*., 2012; Scheffers *et al*., 2016). In particular, changes in species abundance and distribution due to climate change have already been frequently observed (Bowler *et al*., 2017; Lenoir *et al*., 2020), with many species shifting their range towards higher latitudes and elevations (Chen *et al*., 2011). However, some species also respond to climate change through idiosyncratic range shifts (Gibson-Reinemer & Rahel, 2015).

In the past most climate change impact assessments on vertebrate biodiversity have focused on endotherms (birds & mammals). Reptiles, although they account for a third of global terrestrial vertebrate diversity, have been largely ignored (Pacifici *et al*., 2015), and previous assessments of climate change impacts on reptile species have either used only a subset of species (*Warren et al., 2018*; Newbold, 2018) or have not been of global extent (Araújo *et al*., 2006). Moreover, global biodiversity assessments often either consider overall effects on a single taxon (*Baisero et al., 2020*; Voskamp *et al*., 2021) or compare multiple taxa (Hof *et al*., 2018; Newbold, 2018; Warren *et al*., 2018; Thuiller *et al*., 2019), but only very rarely compare different taxonomic groups within one taxon (but see, e.g. Hof *et al*. 2011).

Reptiles have in the past also often been neglected when assessing global conservation priorities (Brooks *et al*., 2006). They are most diverse in arid and semi-arid regions, which suggests that their distributions are driven by ecological and evolutionary processes that differ from other vertebrate taxa, and these regions have been previously unrecognised as conservation priorities because other vertebrate taxa could be more efficiently protected elsewhere (Roll *et al*., 2017).

While reptiles as a whole have been found to use similar habitats to mammals and birds (Cox et al. 2022), treating all reptile species as uniform may be problematic because reptile taxonomic groups (lizard, snakes and turtles) exhibit large differences in species richness hotspots (Roll *et al*., 2017) and habitat use, making them vulnerable to different anthropogenic impacts (Cox *et al*., 2022).

As reptiles are ectothermic, they are likely to be strongly influenced by climate warming, with some species already experiencing body temperatures above their physiological optima. This could indicate a higher vulnerability of these species to climate warming compared to species from cold environments (Diele-Viegas *et al*., 2018). Nonetheless, temperate species are also likely to be vulnerable, assuming that their physiological adaptations for living in cold environments may hinder their ability to cope with hotter climates (Monasterio *et al*., 2013). However, previous studies assessing climate change impacts on reptile species are biased towards certain species and taxonomic groups as well as zoogeographic realms (Diele-Viegas *et al*., 2020).

The aim of our research was to provide a detailed account of projected climate change impacts on global reptile distributions and diversity, looking at species-specific changes as well as broad-scale geographic trends across and within different taxonomic groups. We assessed changes in reptile species richness globally, within each zoogeographic realm and across their respective climate space. For each species, we further quantified the change in range extent, range overlap and range distribution and again assessed differences across zoogeographic realms and taxonomic groups. Given that we cannot model range-restricted species, we also performed a more general assessment of species-specific changes in climate space across both modelled and non-modelled species.

## Methods

### Species data

Until recently, global reptile distribution data was unavailable, but this has changed with the release of the global distribution database by the Global Assessment of Reptile Distributions (GARD) initiative (Roll *et al*., 2017) and more recently the release of the full set of IUCN reptile range maps (IUCN, 2022). We obtained global range maps of 10,064 reptile species from the GARD initiative (Roll *et al*., 2017). The range maps cover lizards, snakes, turtles, worm lizards, crocodiles and the tuatara, but in this paper, in an approach similar to Roll *et al*. (2017), we only contrast snakes, turtles and paraphyletic lizards (for simplicity, we subsequently refer to the latter as lizards).

Range maps were gridded to a 0.5° x 0.5° grid in WGS84, to align with the available climate data (see next section), considering any grid cell that intersected with a species’ range polygon as a presence. As range maps only provide information on a species’ presence but not on a species’ absence, pseudo-absence data for each species were generated by randomly selecting grid cells with no presence, using a distance-weighted approach, where grid cells closer to the range edge were favoured over grid cells further away (see *Hof et al*., 2018). The number of absences was either equal to the number of presences for species with 1000 or more presences or 1000 absences for species with less than 1000 presences because a minimum of 1000 absences considerably increases model performance (Barbet-Massin *et al*., 2012). For each species we derived 10 replicate sets of pseudo-absences, to account for the variability in model accuracy because of the random sampling of pseudo-absence data. Barbet-Massin *et al*. (2012) found that depending on the number of pseudo-absences and model algorithm chosen, 5 to 12 replicates provide the best model quality. Given the relatively high number of pseudo-absences, 10 replicates should thus result in high quality for our models. We created a separate model for each of these 10 sets, but the results were then averaged across the 10 sets.

### Climate data

Global bias-corrected daily climate (minimum temperature, maximum temperature and precipitation) data at a spatial resolution of 0.5° (WGS84) were obtained from the meteorological forcing dataset ‘EartH2Observe, WFDEI and ERA-Interim data Merged and Bias-corrected for ISIMIP’ (EWEMBI; Lange, 2016) for current conditions (1980-2009) and from the Inter-Sectoral Impact Model Intercomparison Project phase 2b (ISIMP2b; Frieler *et al*., 2017) for future simulations (2036 - 2065 & 2066 - 2095). Future climate simulations were available from four global circulation models (GCMs; GFDL-ESM2M, HadGEM2-ES, IPSL-CM5A-LR and MIROC5) and for three representative concentration pathways (RCPs; RCP2.6, RCP6.0 and RCP8.5) under the Coupled Model Intercomparison Project Phase 5 (CMIP5). Future climate simulations strongly depend on the GCM used (Watterson, 2019). The four GCMs chosen by ISIMIP2b capture a large range of plausible future climate projections from all the different GCMs available (Frieler *et al*., 2017), while the RCPs represent different emission scenarios depending on the ongoing and future trajectories of global CO_2_ emissions (Van Vuuren *et al*., 2011).

Monthly means of each climate variable over the respective 30-year time periods, centred around 1995, 2050 and 2080, and for each future scenario (GCM & RCP) were used to calculate 19 bioclimatic variables (see Table S1.1 in Appendix S1 in Supporting Information) using the biovars() function of the ‘dismo’ package (Hijmans *et al*., 2021) in R (R Core Team, 2021). Bioclimatic variables represent annual trends, seasonality and extreme or limiting environmental factors and are thus more biologically meaningful variables than temperature or precipitation alone.

### Species Distribution Models (SDMs)

Species distribution models (SDMs) are a common way of assessing species-specific responses to climate change (Guisan & Thuiller, 2005), but are also used to assess climate change impacts on biodiversity (Thuiller *et al*., 2005). SDMs statistically infer a relationship between the observed distribution of a species and the underlying environmental conditions (Elith & Leathwick, 2009), and can then be used to project current distributions into the future (*Elith et al., 2010*), assuming that the species maintains its climatic niche (Wiens & Graham, 2005). By doing this for multiple species, these projections can be combined to assess future changes in species richness (e.g. Hof *et al*., 2018).

We fitted species distribution models using the presence/pseudo-absence data of a species as response variable and the derived bioclimatic variables for current (1995) conditions as explanatory variables. The number and choice of explanatory variables used strongly influences the outcome of the SDMs (Petitpierre *et al*., 2017). We thus performed a rigorous variable selection approach. After pre-selecting the 10 most commonly used variables from the literature (see Porfirio *et al*., 2014), all potential combinations of three and four bioclimatic variables with a low Pearson correlation (*r ≤* 0.7) were used to model 10 % of all species (N= 987), which were randomly selected, using a Generalised Additive Model (GAM) approach (see Hof *et al*., 2018). The model performance of the different variable combinations was tested and models for all species were fitted using the best-performing variable combination, which was temperature seasonality, maximum temperature of the warmest month, annual precipitation and precipitation seasonality (see Fig. S1.1).

Projections based on SDMs vary considerably among model algorithms (Thuiller *et al*., 2019), therefore we fitted two different modelling algorithms with good performance and discrimination capacity (Meynard & Quinn, 2007; Elith *et al*., 2010), an additive model (GAM) and a regression tree based model (Generalised Boosted Regression Models (GBM)).

GAMs were fitted with a Bernoulli response, a logit link and thin-plate regression splines using the ‘mgcv’ package (Wood, 2003, 2011) in R (R Core Team, 2021). GBMs were fitted with the ‘gbm’ package (Greenwell *et al*., 2020) in R (R Core Team, 2021) and the optimal parameter settings for learning rate (0.01 and 0.001), tree complexity (1, 2 and 3) and number of trees (1000-10000) for each species were identified by cross-validation (Bagchi *et al*., 2013).

Spatial autocorrelation in species distributions can bias parameter estimates and error probabilities (Kühn, 2007). Two different methods were used to account for spatial autocorrelation in the SDMs. Species with equal or more than 50 presences were modelled using an ecoregion-blocking approach. Here the world was divided into 10 blocks based on a representative subset of the climate space across each of the world’s ecoregions (Olson *et al*., 2001, Bagchi *et al*., 2013). Subsequently, 10 models per species were built leaving out one block at a time, using the left-out block for model evaluation (Bagchi *et al*., 2013). For species with 10 to 49 presences, we split the data into 10 datasets by repeatedly randomly selecting 70% of the data, then using the left-out 30% for model evaluation. Species occurring in less than 10 grid cells (N = 3602, Table 1) were not modelled because the sampling size would be too low to produce meaningful results (Hernandez et al. 2006). Together with the 10 different pseudoabsence sets this resulted in 100 models for each model algorithm and species.

**Table 1.** Number of species that were excluded from the species distribution models due to their restricted range or low model performance.

| Taxonomic group | Lizard | Snake | Turtle | Total |
| --- | --- | --- | --- | --- |
| No. of species with available data | 6328 | 3414 | 322 | 10064 |
| No. of range-restricted species (N < 10, removed) | 2536 | 1047 | 19 | 3602 |
| No. of species with low model performance (AUC < 0.7, removed) | 97 | 62 | 7 | 166 |
| **Total number of species modelled** | **3695** | **2305** | **296** | **6296** |
| Percentage of available species modelled | 58.4 % | 67.5 % | 91.9 % | 62.6 % |

The performance of the fitted SDMs was evaluated by calculating the overall AUC for each species (the average AUC across the 10 blocks and the 10 sets of pseudo-absences). Models with an overall AUC smaller than 0.7 were dropped (N = 166), which left us with SDMs for 6296 reptile species (see Fig. S1.2), which represents 62.6 % of the total number of available species from GARD (Table 1). In addition to assessing the model fit of the individual models, we also compared the observed species richness with the projected current richness per grid cell (Fig S1.3) in order to assess the performance of all our models when looking at changes in species richness.

The same modelling approach has been used previously to assess climate change impacts on amphibians, birds and mammals, see Hof *et al*. (2018) and Biber *et al*. (2020). The former provides a more detailed explanation of the modelling methodology, while the latter gives a thorough account of the caveats and uncertainties associated with species distribution models.

### Future projections

Future species distributions were projected using the future bioclimatic variables for the two future time periods (2050, 2080), each GCM (GFDL-ESM2M, HadGEM2-ES, IPSL-CM5A-LR and MIROC5), each RCP (RCP2.6, RCP6.0 and RCP8.5) and both model algorithms (GAM & GBM). Model results are presented as the ensemble mean across the four GCMs and two model algorithms (GAM & GBM) considered.

Future projections of each species were limited to the extent of their original and the neighbouring ecoregions to prevent predictions of areas with analogue climatic conditions. Future projections were further limited by applying a species-specific dispersal buffer. For most species considered in this paper, species-specific dispersal distances are still unknown (*Nathan et al., 2012*), hence we used species-specific dispersal buffers which were based on the diameter (d) of the largest range polygon of a species. We used three species-specific dispersal scenarios (d/4, d/8, d/16, see Fig. S1.4) and provide a detailed comparison of these in the Supporting Information (see Appendix S5). Here we provide results under a medium dispersal scenario (d/8), which corresponds to a mean dispersal distance of 2.4 km per year.

### Impact analysis

The current and future probabilities of occurrence of the individual SDMs were thresholded into binary presence-absence data using species-specific thresholds according to the true skill statistic (MaxTSS; Allouche *et al*., 2006; Fig. S1.5 d). Thresholded species occurrences were then used to calculate current and future species richness, as well as richness increase, decrease, change and relative change (%). Richness increase and decrease were identified by using the presence information of each individual species and then summing up the number of species that newly occur in a given grid cell (species increase) or species that disappear from the respective grid cell (species decrease).

Summing up the thresholded species occurrences frequently overestimates species richness (Calabrese et al., 2014), thus we also present the results using the sum of the raw non-thresholded probabilities of occurrence of each species in the Supporting Information (see Appendix S4).

We calculated the projected species richness for each cell of a global grid of 0.5° x 0.5° resolution globally as well as for each zoogeographic realm, as defined by Holt *et al*. (2013), for each time period. We then tested for significant changes in species richness over time using a paired t-test with Holm correction. To assess how species richness and richness changes are related to the overall change in climatic conditions, we assessed both against univariate temperature and precipitation as well as the interaction of temperature and precipitation conditions. To assess potential future climate effects on individual species, we quantified the percentage of change in range extent, the percentage of range overlap and the direction and distance in range shift. The percentage of change in range extent was calculated based on the projected current and future range extent of a species (Fig. S1.5 f). The percentage of range overlap was calculated by extracting the total area of spatial range overlap between the projected current and future range extent and dividing it by the projected current range extent (Fig. S1.5 g). To assess the magnitude of the range shift for each species, we derived the range centroid for both time periods and calculated the distance and direction of the projected range shift (Fig. S1.5 g).

Given that 37.4% of all reptile species for which data were available could not be modelled (largely due to their restricted range extent, Table 1), we performed an additional analysis considering all 10,064 species for which data were available. We used the same 4 bioclimatic variables used for the SDMs to transform the multidimensional climate data to a two-dimensional climate space using the first two axes of a principal component analysis (PCA). PCAs were performed for both current and future conditions, taking into consideration the same GCMs, RCPs, and time periods as previously. The explained variance of the first two PCA axes was above 75 % under all scenarios (Fig. S1.6). For each scenario combination, we then calculated the Euclidean distance between the two PCA axes of current and future conditions, to obtain a measure of climatic change (Fig. S1.7). We then extracted the climatic distance for the gridded locations of each species and compared the climatic distance of modelled and non-modelled (range-restricted) species using a non-paired t-test with Holm correction.

Where no specific groups (lizards, snakes, turtles) are mentioned, we present the results for all reptile species together. Nevertheless, Appendix S2 presents group-specific results. Results are presented for the year 2080 under a medium representative concentration pathway (RCP6.0). A sensitivity analysis with regards to the variation across years and RCPs is shown in the Supporting Information (see Appendix S6).

## Results

### Spatial changes

Projected reptile richness (sum of thresholded SDM projections) based on a resolution of 0.5° x 0.5° for current conditions varied between 0 at high latitudes and 251 in the tropics, with particular hotspots in Brazil, Cameroon and Indonesia (Fig. 1a). Overall, reptile richness was dominated by lizard species (N = 3695), followed by snakes (N = 2305), while turtle species only contributed marginally to the total number of modelled species (N = 296, Table 1, Fig. 1b).

**Figure 1.**
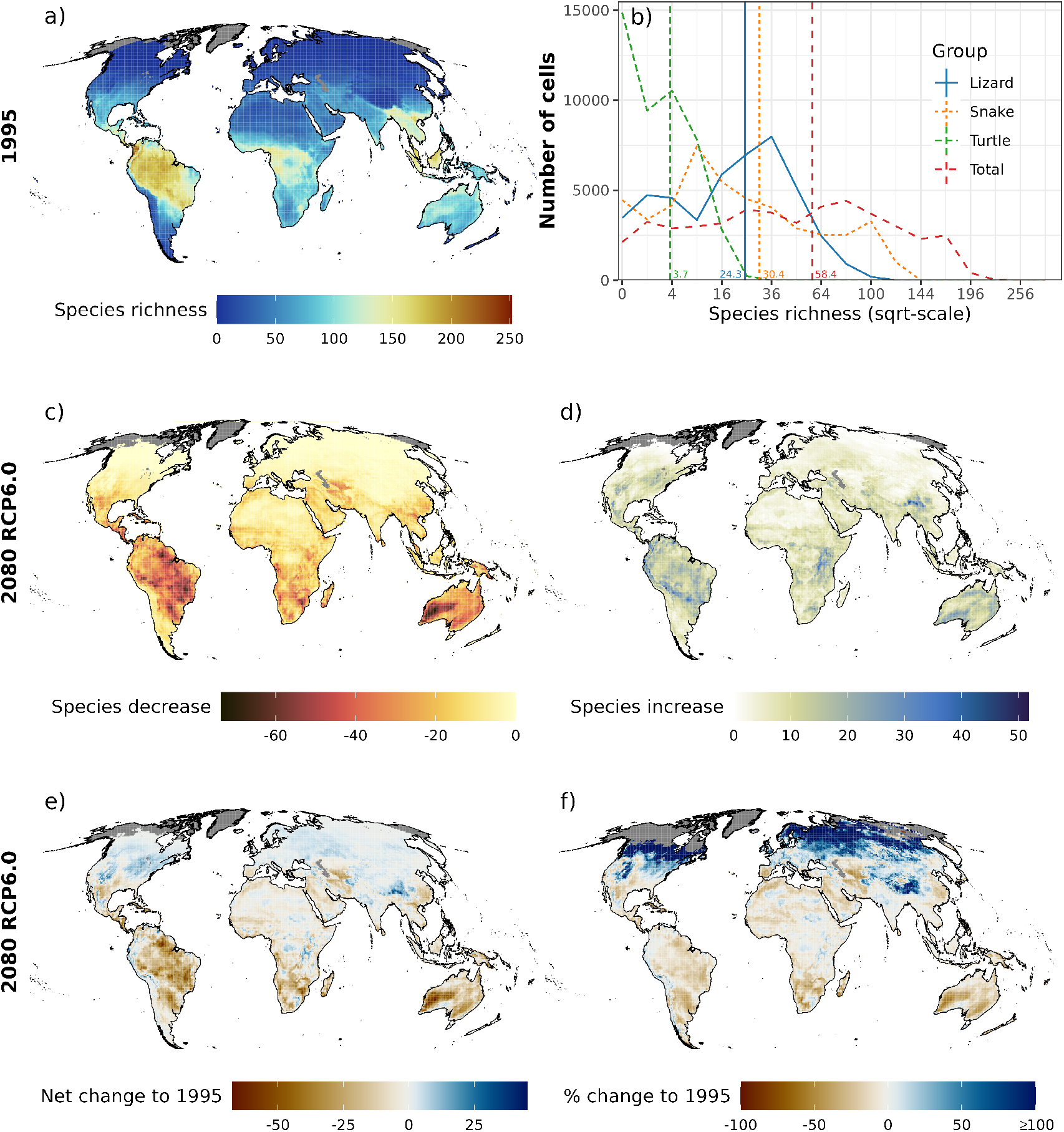
a) Map of projected global terrestrial reptile species richness (1995), b) frequency of species richness by taxonomic group (lizard, snake, turtle and total) with mean values highlighted by vertical line and c) increase, d) decrease, e) net change and f) relative change (%) in reptile species richness for all modelled reptile species (N = 6296) for the year 2080 under a medium representative concentration pathway (RCP6.0) and a medium dispersal scenario (d/8). Results are presented as the ensemble mean, across the four global circulation models (GCMs) and two model algorithms (GAM & GBM) considered. All maps are based on 0.5° x 0.5° grid cells, which have been projected to Mollweide equal-area projection (EPSG:54009). Grey areas are regions for which no projections are available. Note that the colour scales differ among the individual panels.

For future conditions, a large number of reptile species were projected to disappear and at the same time a large number of new species were projected to appear in Brazil and Australia, while other regions showed either a strong species decrease or increase (Fig. 1 c, d). The greatest future decreases of species richness were projected east of the Caspian Sea and in South Africa (Fig. 1 c), while strong future increases were projected in the south-west of China and in the eastern United States (Fig. 1 d). Overall, the projected richness decrease was greater than the richness increase, which resulted in a greater net loss in species richness from 1995 to 2080 (Fig. 1 c, d, e). The lowest net loss in species richness was projected for Brazil, Australia and South Africa, while the highest net gain was projected for south-west China and the western United States (Fig. 1 e). Relative change (%) was projected to be negative - in particular for most of the southern hemisphere, while the high northern latitudes showed a strong positive relative change (Fig. 1 f).

Spatial patterns in species richness changes varied greatly across the three taxa, with lizards seeing both strong increases and decreases in Australia, snakes showing a strong decrease in South America and turtles seeing a strong increase in the eastern part of North America (Fig. S2.9 & Fig. S2.11). All three taxa showed a net gain in species richness in northern latitudes, while lizards showed the greatest net loss in parts of Australia, snakes in large parts of South America and turtles in parts of South America and southern Africa (Fig. S2.10 & Fig. S2.12).

### Global and zoogeographic realm changes

Summed up across all 0.5° x 0.5° grid cells globally, species richness was projected to decline significantly (*p* < 0.01) from 1995 to 2080 globally, with a decline in mean reptile richness per grid cell from 58.4 ± 0.24 (SE) in 1995 to 53.39 ± 0.19 in 2080 (Fig. 1b, Fig. 2 a). 8 out of 11 zoogeographic realms showed a significant decline in reptile richness by 2080 (Fig. 2 b, c, d, f, g, h, j, k), while the Nearctic and Palearctic realms showed a significant increase (Fig. 2 e, i) and the Sino-Japanese realm showed no significant change (Fig. 2l).

**Figure 2.**
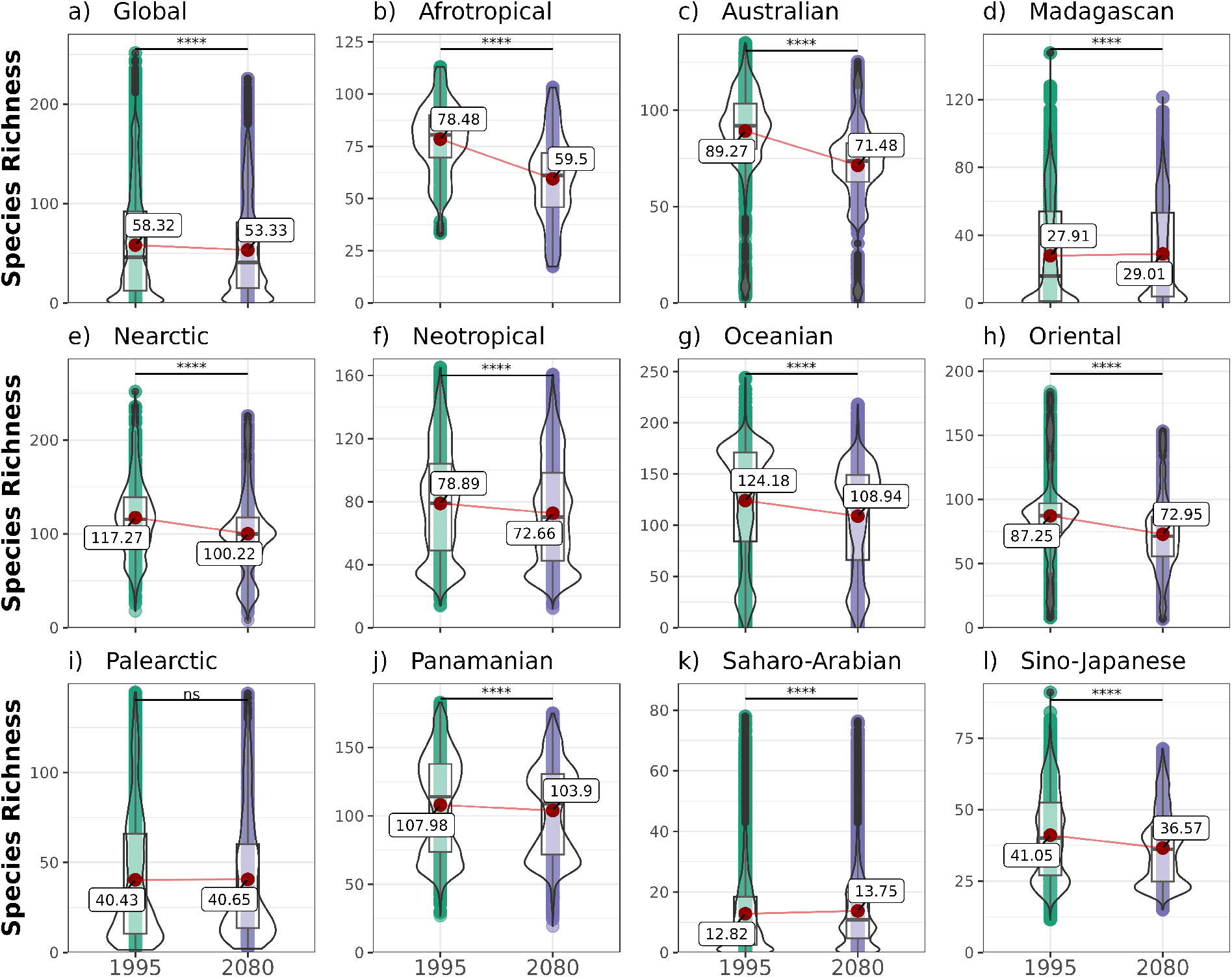
Terrestrial reptile species richness across the globe and for each zoogeographic realm (Afrotropical, Australian, Madagascan, Nearctic, Neotropical, Oceanian, Oriental, Palearctic, Panamanian, Saharo-Arabian & Sino-Japanese) over time (1995, 2080) based on all modelled reptile species (N = 6296). Results are presented as the ensemble mean across the four global circulation models (GCMs) and two model algorithms (GAM & GBM) considered, under a medium representative concentration pathway (RCP6.0) and a medium dispersal scenario (d/8). The statistical difference between years was tested using a paired Student’s t-test with Holm correction (*p* < 0.05 = *, *p* < 0.01 = **, *p* < 0.001 = ***, *p* < 0.0001 = ****). Plots show mean (red point & label), median (black horizontal line), 25th to 75th percentiles (box), entire range of data (violin & data points) and density of values (width of violin). Figure 5 provides a map outlining the different zoogeographic realms.

Looking at the global averages of species richness for each 0.5° grid cell for the three reptile groups separately, snakes had the highest mean species richness (μ_mean_ = 30.4 ± 0.15), followed by lizard (μ_mean_ = 24.3 ± 0.09 SE) and turtle richness (μ_mean_ = 3.71 ± 0.02 SE; Fig. 1b, Fig. S2.8). Globally, similar to the total reptile richness, the individual taxonomic groups (lizards, snakes and turtles) all showed a significant decline in species richness, while there were slight differences across the individual realms (Fig. S2.13 – S2.15). Lizards only showed a significant increase in richness in the Palearctic realm and no significant change in richness in the Sino-Japanese realm, while in all other realms they showed a significant decrease (Fig. S2.13). Snake and turtle richness increased significantly in the Nearctic and Palearctic realms. Snake richness decreased significantly in all other realms apart from the Sino-Japanese one (Fig. S2.14), while turtle richness significantly decreased in all other realms apart from the Saharo-Arabian and the Sino-Japanese one (Fig. S2.15).

### Biophysical changes

Reptile richness varied greatly across conditions with varying combinations of temperature and precipitation (Fig. 3 a, b, c). For 1995, reptile richness was projected to be highest in areas with a temperature of about 28.5°C, a precipitation of about 5500 mm and when considering temperature and precipitation together in warm and moist regions (21°C & 3000 mm, Fig. 3 c). The climatic conditions with the highest richness shifted to even more extreme (warmer & wetter) novel climate conditions by 2080 (Fig. 3 a, b, d).

**Figure 3.**
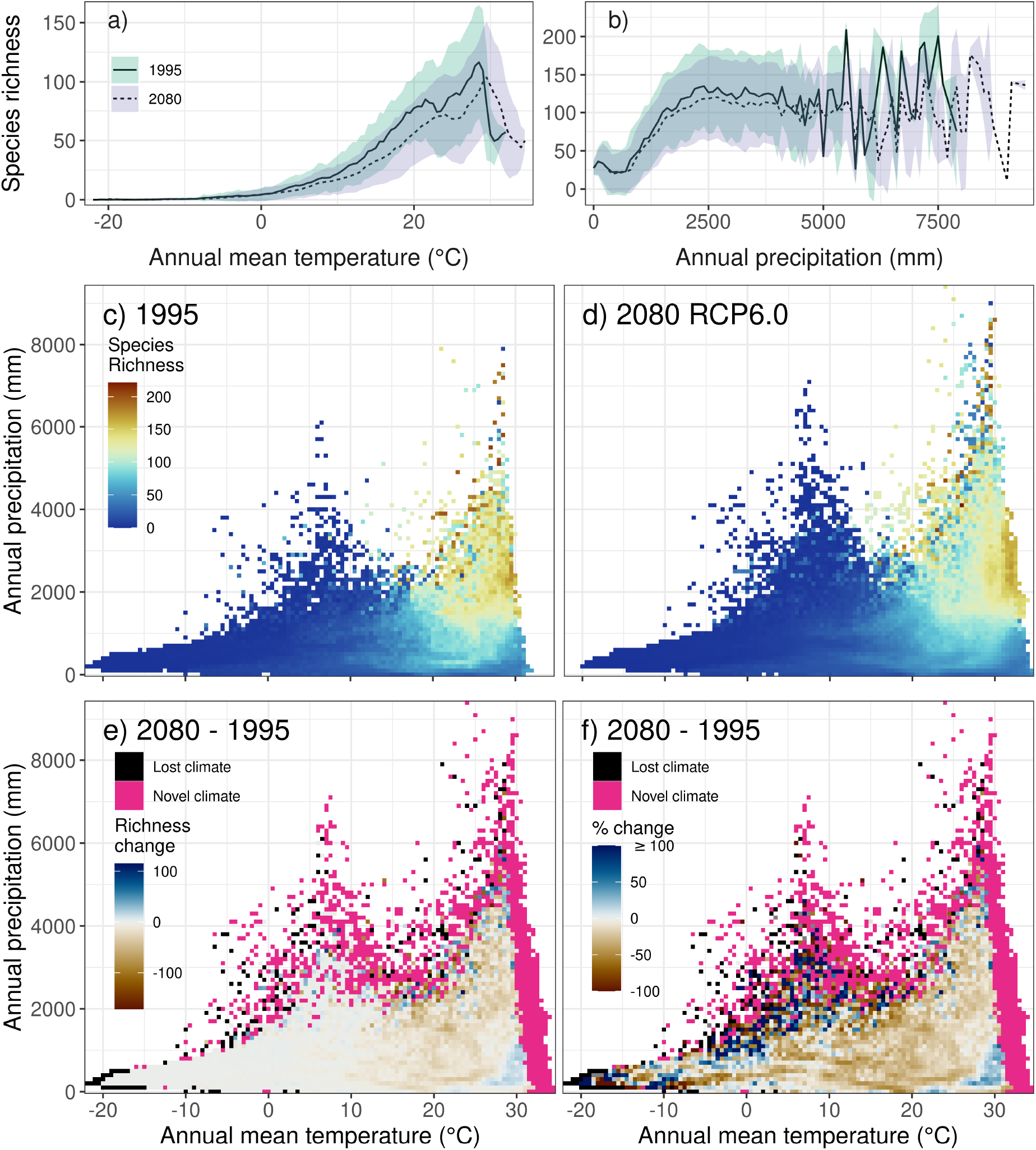
Univariate relationship of current (1995) and future (2080 RCP6.0) reptile species richness with a) temperature, b) precipitation; and the bivariate relationship of temperature and precipitation with reptile species richness for c) 1995 and d) 2080 RCP6.0; and the respective e) net richness change and f) relative richness change (%) under a medium dispersal scenario (d/8). Heat maps and lines show the mean and ribbons the standard deviation in variance across space, global circulation models (GCMs) and the two model algorithms (GAM & GBM).

Looking at the species richness change across the 2-dimensional climate space, net change was positive at the upper precipitation limits across all temperatures and the very hot and very dry conditions and negative throughout the entire precipitation range especially for the higher temperatures. Overall, the negative change was much greater and more pronounced than the positive net change (Fig. 3 e). The highest positive and negative relative change values were clustered, both occurred at the upper precipitation limits at low and medium temperatures (Fig. 3 f). A considerable percentage of the climate space (29.5 %) was shifting towards novel climatic conditions, for which no change in species richness could be estimated, while only few discrete climatic conditions as well as very cold & very dry conditions (4.75 %) got lost (Fig. 3 e, f).

### Species-specific range changes

The range extent of most species (N = 6021) showed a considerable decrease (μ_mean_ = −27.7 ± 0.16 SE, Fig. 4 a, c, e). Lizard species showed the greatest decline (μ_mean_ = −31.8 ± 0.22 SE) in range extent (Fig. 4 a), followed by snakes (μ_mean_ = −22.6 ± 0.25 SE), while almost equal numbers of turtle species showed a decline (N = 274) and an increase (n = 205) with decreases being much more pronounced than increases (μ_mean_ = −17.5 ± 0.72 SE, Fig. 4 e). Almost half of the modelled reptile species (N = 3029) showed a strong change in range position, demonstrated by a relatively low range overlap (<= 60 %), which was consistent across all three groups (Fig. 4 b, d, f).

**Figure 4.**
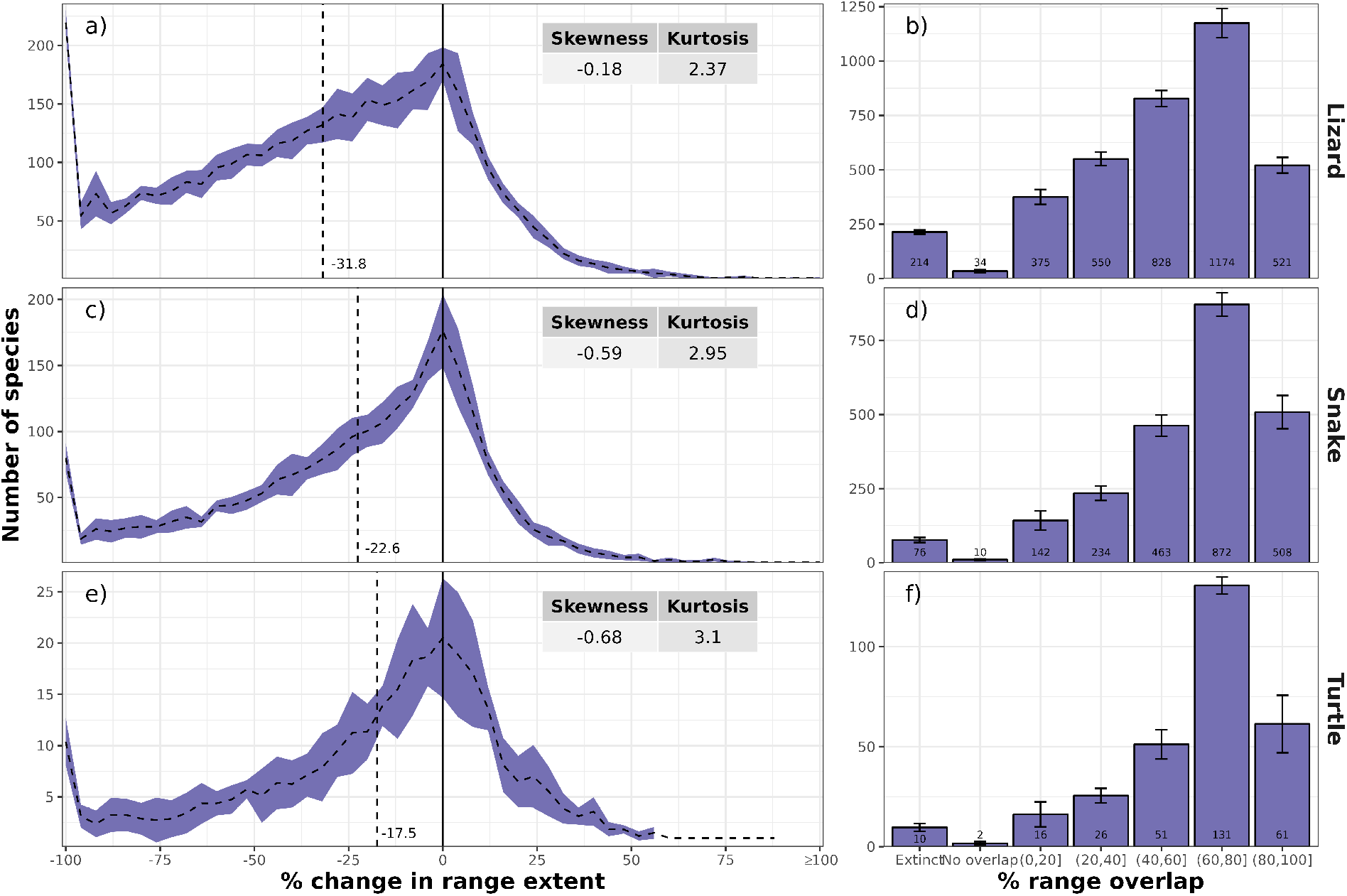
Frequency plots of the mean number of reptile species (a) lizard, c) snake, e) turtle) and their potential future change (%) in range extent (total area of occupied grid cells) and the mean number of reptile species (b) lizard, d) snake, f) turtle) per potential range overlap class (0-20, 20-40, 40-60, 60-80, 80-100). Error margins/bars indicate the standard deviation across the four global circulation models (GCMs) and the two model algorithms (GAM & GBM) used. Both shown for 2080 under a medium representative concentration pathway (RCP6.0) and a medium dispersal scenario (d/8).

Most of the range centroids (58 %) of all reptile species fell within the Neotropical (N = 1133), Afrotropical (N = 1039), Oriental (N = 785) and Australian realm (N = 698). Turtle species had 50 % of their range centroids in the Nearctic (N = 58), Oriental (N = 53) and Afrotropical realm (N = 38), while lizards and snakes reflected the overall, total reptile patterns (Fig. 5 d). Range centroids were highly clustered within the different realms, which reflects the overall richness hotspots, and hardly any centroids were found in the high northern latitudes (Fig. 5 d). By 2080 species centroids were projected to shift by a mean distance of 111 km ± 0.9 SE primarily towards the South. Lizards showed a shift in all directions, with a slightly greater number of species exhibiting a shift towards the South (Fig. 5 a), while snakes and turtles showed a more pronounced shift of species towards the North (Fig. 5 b, c). Turtle ranges shifted by the largest distances, followed by snakes (Fig. 5 a, b, c). The northern realms (Nearctic, Saharo-Arabian, Palearctic und Sino-Japanese) showed a dominant shift towards the North, while the southern realms (Neotropical, Afrotropical and Australian) showed a dominant shift towards the South. This was also reflected in the taxon-specific range shifts for the Northern and Southern hemisphere (Fig. S3.16 in Appendix S3). The Panamanian, Madagascan and Oriental realms also showed a northerly shift, while the Oceanian realm showed a bi-directional shift to the Northwest and Southeast (Fig. 5 d, Fig. S3.17). Large realms had a greater percentage of species that shifted their range over a greater distance (Fig. 5, Fig. S3.17).

**Figure 5.**
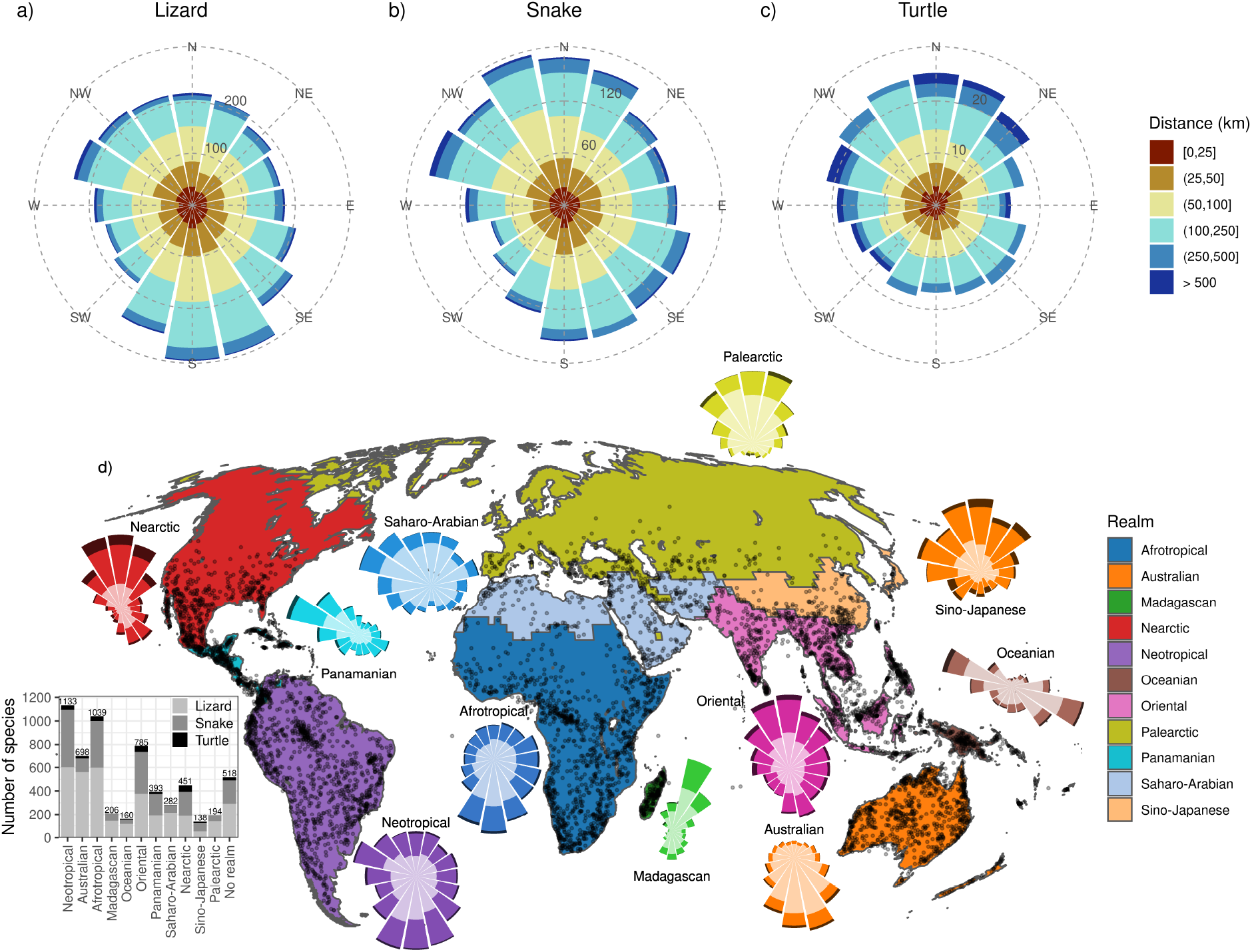
Cumulative direction and distance of potential range centroid changes per taxonomic group (a) lizard, b) snake, c) turtle) and d) range centroids (points on map) and the number of species and their directional shift in range centroid position per zoogeographic realm (inset polar plots). Results are presented as the ensemble mean across the four global circulation models (GCMs) and two model algorithms (GAM & GBM) considered, for the year 2080 under a medium representative concentration pathway (RCP6.0) and a medium dispersal scenario (d/8). The inset bar chart shows the number of species that have their range centroid located in the respective realm. Numbers show the total number of species per realm, while the bars are colour coded by the three different taxonomic groups. Please note that a considerable number of species (n = 518) have their range centroid located outside of the zoogeographic realm boundaries and thus were not associated with any realm and so are not considered for the polar plots in Fig. 5 d).

### Non-modelled species

37.4 % of reptile species for which data would have been available could not be modelled using SDMs, either due to a small sample size or a low model performance (Table 1). We found that the species that could not be modelled showed a significantly greater (*p* < 0.05) mean climatic distance between current and future conditions than did the modelled species and thus occurred in areas that are projected to experience a greater change in climatic conditions. This pattern was consistent across all three taxa

### Sensitivity analysis

Looking at the sum of occurrence probabilities, we found similar spatial patterns and a similar magnitude of change compared to the sum of thresholded occurrences (species richness) (Fig. S4.18 - S4.20). Projected richness values and their future changes were slightly higher assuming a larger dispersal ability (d/4), but on the whole all results were consistent across the three dispersal scenarios considered (Fig. S5.21 - S5.26). Climate change impacts on future species richness increased over time, with greater effects seen for 2080 than 2050, and the greatest impacts being observed under a high emission scenario (RCP8.5) compared to the two lower scenarios (Fig. S6.27 - S6.38).

## Discussion

Reptile richness was projected to decrease significantly across most parts of the world in the future (Fig. 1 & 2). This effect was apparent for lizards, snakes and turtles alike, although regional and species-specific responses differed across the three groups (Fig. S2.8 – S2.12).

### Spatial changes

Reptile richness is projected to decrease in Brazil, Australia and South Africa (Fig. 1 c, e). These areas overlap significantly with the biotic convergence zones, areas with a high spatial concentration of Lepidosaurians (i.e. snakes and lizards) identified by Diele-Viegas *et al*. (2020), which were also found to cover a large number of the Lepidosaurian species vulnerable to climate change (Diele-Viegas *et al*., 2020). Huey *et al*. (2012) further found that ectotherms sharing climate vulnerability traits seem to be concentrated in lowland tropical forests. Combining climate-based SDMs with land-use change information, Newbold (2018) created future projections for 20938 vertebrate species and found that Brazil is strongly affected by vertebrate diversity loss due to climate change, and, together with Australia, is also likely to be strongly affected by future land-use changes, especially under a high-emission scenario (RCP8.5). Given that Brazil in particular does not only host a high reptile richness (*Roll et al*., 2017; Fig. 1 a), but has recently been found to host a large number of threatened reptile species (Cox *et al*., 2022), this highlights the responsibility of this mega-diverse country for protecting reptile diversity. South-western China and the western United States were projected to show a net gain in reptile richness under climate warming (Fig. 1 e), but they also belong to those areas where most reptile species are threatened by habitat loss from agriculture and logging or the harvesting of species (Böhm *et al*., 2013). These projected losses due to habitat change could potentially counteract any positive effects climate warming might have.

The high variation in species richness changes across regions and taxa that we found obviously reflects their original richness patterns. Species richness of amphibians, birds and mammals together is a good spatial surrogate for species richness of all reptiles combined and of snakes, but is not a good surrogate for lizard or turtle richness (Roll *et al*., 2017). Thus, it is not surprising that the areas with the highest decline in overall reptile richness (see Fig. 1) overlap significantly with the areas of highest projected changes in vertebrate species richness (amphibians, birds and mammals) found by Hof *et al*. (2018), although global reptile richness is largely constrained by temperature, while global richness of all other vertebrate groups is primarily constrained by the availability of energy and water (Qian, 2010). Further, historical shifts in geographic ranges and climatic niches have further demonstrated that niche shifts in endotherms are significantly faster than in ectotherms (Rolland *et al*., 2018).

### Global and zoogeographic realm changes

Globally reptile richness was projected to decline significantly, from an average of about 58 to about 53 (9.4 %) species per grid cell from 1995 to 2080 (Fig. 2 a). This projected percentage change in average future reptile richness (9.4 %) is considerably lower than the reptile richness decline due to climate warming predicted by Newbold (2018). However, while Newbold (2018) also used SDMs to infer future reptile richness changes, he used only a subset of reptile species, for which IUCN range maps were available at that time. In addition, he also applied a much smaller dispersal buffer (0.5 km per year), which might indicate that our projections provide a rather optimistic scenario. Newbold (2018) also found that reptiles, together with amphibians, are disproportionately sensitive to future human land-use. Given the synergistic effect of future climate and land-use changes on biodiversity (Brook *et al*., 2008) as well as species populations (Williams *et al*., 2022), land-use change is likely to further exacerbate climate change impacts on global reptile distribution and diversity.

Changes in reptile richness differed among zoogeographic realms, but species richness declined significantly across most realms over both time periods (Fig. 2 b, c, d, f, g, h, j, k). Lizards, snakes and turtles all showed similar declines in species richness globally and across most realms, but differed slightly across individual realms. This is in line with a previous study covering various realms from tropical to temperate regions which found that 60 % of assessed Lepidosaurian species (N = 1114) were vulnerable to changes in climate (*Diele-Viegas et al*., 2020). Diele-Viegas *et al*. (2020) further found that the Afrotropical, Nearctic and Sino-Japanese realms were the three realms where Lepidosaurians were most vulnerable to climatic change, while Lepidosaurians in the Madagascan and Oceanian realm were least vulnerable. By contrast, we found no significant decline in projected total reptile richness for the Sino-Japanese realm (Fig. 2 l), although individual subgroups were projected to show a significant decline in species richness from 1995 to 2050 (Fig. S2.13 – S2.15). Both the Madagascan and Oceanian realm were further projected to significantly decrease in species richness including for all reptile species (Fig. 2 d, g) and all three subgroups (Fig. S2.13 – S2.15). However, given that the Madagascan realm, specifically, boasts over 90% of endemic reptile species and genera (Glaw & Vences, 2007) and that both realms are composed of island territories which are usually considered highly vulnerable to climate change and might also be affected by future sea level rise and erosion (Diele-Viegas *et al*., 2020), our estimates might still underestimate potential climate change impacts in these realms. Interestingly, turtle richness seemed to be least affected by climate change, when looking at the different realms (Fig. S2.15), although a recent review has deemed turtles as the vertebrate group with the highest extinction risk, as it is strongly affected by habitat loss, human consumption and pet trading (Stanford *et al*., 2020).

### Biophysical changes

Reptile richness differed significantly with temperature and precipitation, with the highest richness being observed in warm and moist conditions. Under future climate change scenarios, the climatic conditions with high species richness were projected to shift to even more extreme (warmer & wetter) conditions (Fig. 3). This gives cause for concern given that tropical forest and desert lizards already live in environmental conditions that are close to their thermal limits (Sinervo *et al*., 2010), while desert and temperate lizard species have been found to be less able to regulate their temperature in order to deal with heat stress than tropical species (Anderson *et al*., 2022). Reptiles cannot regulate their body temperature internally, so are strongly dependent on using solar energy captured by the environment to regulate their body temperature (Huey, 1982). This might lead to overheating when temperatures go beyond a species’ critical limit, which makes them particularly susceptible to climatic changes (Sinervo *et al*., 2018). However, this might be compensated by other biological processes that help species to buffer climate change effects, i.e. genomic and phenotypic plasticity (Rodríguez *et al*., 2017) as well as behavioural and physiological adaptation (Sunday *et al*., 2014). Overall, the persistence of reptile species would be much more affected by climate cooling than warming, but it has been suggested that increasing droughts, which will be a consequence of continued warming, pose a significant future threat to European reptiles (Araújo *et al*., 2006). It is likely that climate warming will have an additional impact on reptiles that have temperature-dependent sex determination. Altered sex ratios will not only result in a higher extinction risk for local populations, but, together with a reduction in nesting sites due to habitat destruction and fragmentation, will also affect the dispersal and potential range expansion of a species. Therefore it could also have an impact on population demography and size unless temperature shifts in sex determination or female nest-site choice evolves in pace with rising temperatures (Boyle *et al*., 2016; Gibbons *et al*., 2000).

### Species-specific range changes

The range extent of most species were projected to considerably decrease, with lizard species showing the greatest decline (Fig. 4 a). Further, most reptiles also showed a strong decline in range overlap, which was consistent across all three groups (Fig. 4). This is in line with results by Warren *et al*. (2018), who found that projected future range losses of more than 50% occur in 8 - 52% of considered reptile species by 2100 depending on the climate scenario considered, although this study included only a fraction of all reptile species (N=1850) and no species dispersal was considered.

Compared to other terrestrial vertebrate groups (especially birds and mammals), reptiles have small geographic ranges, which also indicates narrower niche requirements. This is not only likely to make them more susceptible to future climate change (Newbold, 2018), but also to other threats such as habitat loss or invasive species (Böhm *et al*., 2013, Cox *et al*., 2022). Reptiles are a paraphyletic class with a diverse range of body forms, habitat affinities and functional roles (Pincheira-Donoso *et al*., 2013), which will probably result in their response to climatic and habitat changes being equally varied. In addition, cascading effects generated by disease, invasive species, habitat loss and climate change might lead to declines of sympatric species and a faster deterioration of ecosystem structure than anticipated for climate change alone (Zipkin *et al*., 2020).

The majority of reptile species showed projected shifts towards the South, which was largely driven by range shifts in lizards (Fig. 5). Comparing the Northern and Southern hemispheres (Fig. S3.16), we found a clear overall poleward shift in species ranges across all three groups, which has also been found previously in various other taxonomic groups (Chen *et al*., 2011). Turtle ranges were projected to shift farthest, followed by snakes (Fig. 5). The relatively short range shift distances in lizard species are probably due to the fact that lizards have the smallest range extents across the three groups (Roll *et al*., 2017), which, given that our dispersal buffers are based on range extent, also resulted in smaller dispersal distances for lizards compared to the two other groups (Fig. S1.4). This is, however, obviously somewhat counter-intuitive and in stark contrast to generally perceived differences in mobility characteristics (e.g. movement speed) among lizards, snakes and turtles. While it is hard to tackle this problem in a globally and taxonomically comprehensive study like ours, it underlines the need for more efforts in collecting data on actual, empirically quantified dispersal distances in order to consider them in SDM exercises and large-scale biogeographical analyses.

### Non-modelled species

Range-restricted (non-modelled) species were projected to experience significantly higher shift in climatic distance than modelled species (Fig. 6), indicating that range-restricted species are also likely to be strongly affected by climate change. This highlights once more that sample size restrictions of SDMs are likely to downplay climate change effects on narrow-ranging and threatened species (Platts *et al*., 2014). Hof *et al*. (2018) also found significant impacts of climate and land-use changes on range-restricted vertebrate species, excluding reptiles. However, similar to the latter study, we only looked at climate anomalies (Euclidean distance between current and future climatic conditions) as a metric of climate change, while different metrics have been found to indicate contrasting climate change patterns on a global scale (Garcia *et al*., 2014). In addition to climate change effects, habitat modification has been found to have a greater impact on range-restricted reptile species, as well as species with a small clutch size (Doherty *et al*., 2020). Range-restricted reptile species are further often evolutionarily unique (Murali *et al*., 2021) and have been found to overlap least with current conservation priority areas (Roll *et al*., 2017, Cox *et al*., 2022).

**Figure 6.**
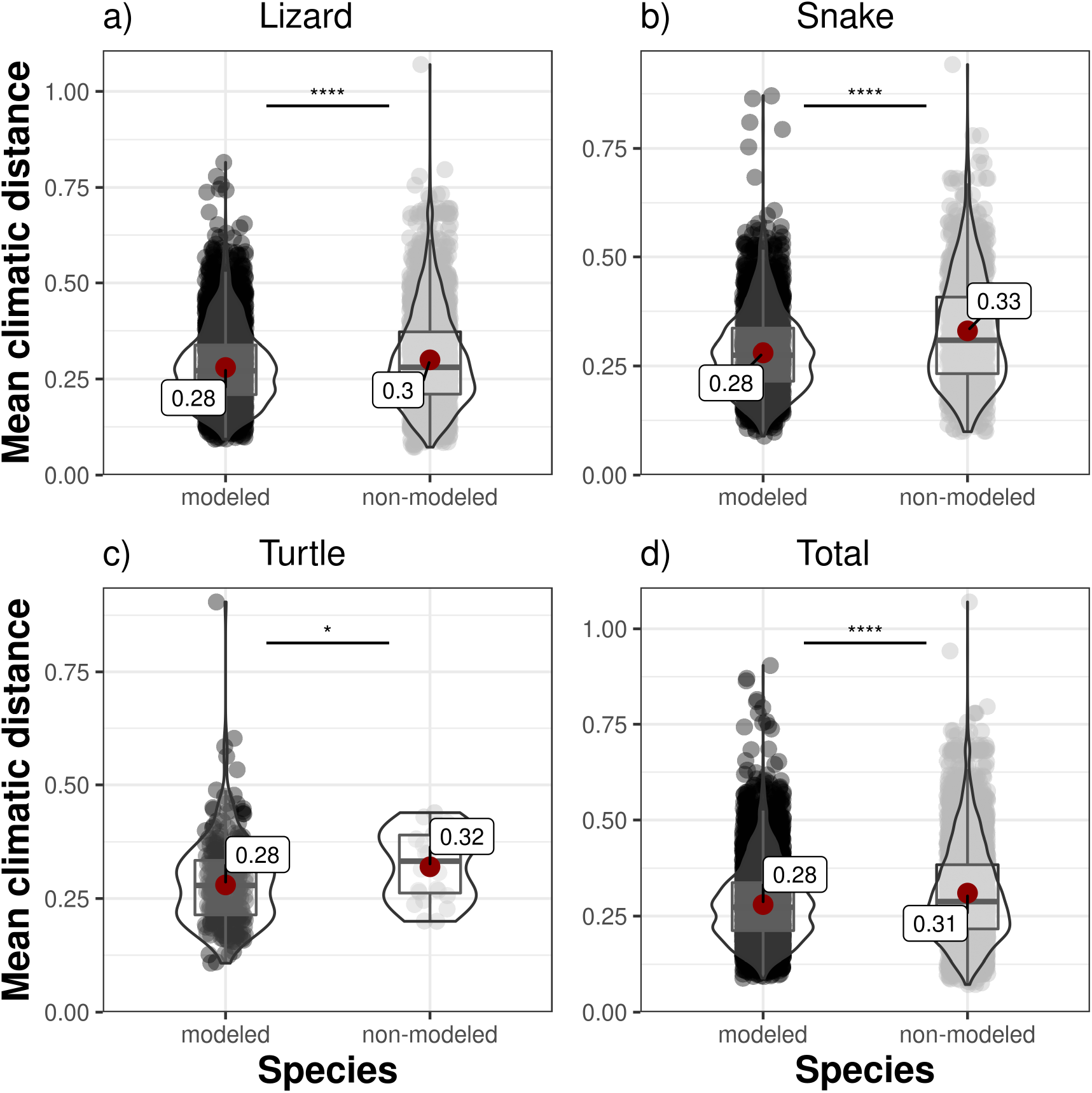
Mean climatic distance for modelled and non-modelled species, split by taxonomic group (a) lizard, b) snake, c) turtle and d) total). The statistical difference between modelled and non-modelled species was tested using a Student’s t-test with Holm correction (*p* < 0.05 = *, *p* < 0.01 = **, *p* < 0.001 = ***, *p* < 0.0001 = ****). Plots show mean (red point & label), median (black horizontal line), 25th to 75th percentiles (box), entire range of data (violin & data points) and density of values (width of violin). The results are shown as the ensemble mean across the four global circulation models (GCMs) for the year 2080 under a medium representative concentration pathway (RCP6.0).

### Sensitivity analysis

Our results were strongly dependent on the dispersal assumption, time period and emission scenario (RCP) considered (see Appendices S5 & S6). As expected, the overall patterns and richness changes were more pronounced in a later time period and scenarios representing higher levels of greenhouse gas emissions, as these reflect potential futures with a higher level of climate warming. Thuiller *et al*. (2019) have previously assessed the uncertainty originating from dispersal, model algorithm, GCM and RCP on the future biodiversity scenario of amphibians, birds and mammals and found that model algorithm and RCP have the greatest influence.

On the contrary, greater dispersal distances imply that reptile species are able to move greater distances in order to track their optimal climatic niche and thus provide more optimistic potential changes in species occurrence and richness patterns. Reptile-specific studies have either considered no dispersal at all (Araújo *et al*., 2006; Warren *et al*., 2018) or a dispersal rate of 0.5 km per year (Newbold, 2018). We use species-specific dispersal buffers with an average of 2.4 km per year (Fig. S1.4). While these buffers might be optimistic, they are based on the transparent rationale of a range-size dependent dispersal buffer, i.e. species with a small range size have a smaller dispersal buffer than species with a large range size, which avoids the unlikely assumption of uniform dispersal distances across species. Given that our model results, as well as the underlying climate scenarios, are based on a 0.5° grid size (about 50 x 50 km), small differences in dispersal distance do not have a strong impact on our results (see Appendix S5). Nevertheless, these considerations again highlight the challenges in sensibly accounting for the influence of dispersal in global climate impact assessments on species distributions and diversity.

The projected changes in species distributions help to investigate potential changes in global reptile richness patterns and in highlighting potential hotspots of climate change impacts. They also allow comparison of climate change vulnerability across taxonomic groups and help to identify areas where conservation efforts might be most urgently needed (Voskamp et al. 2022). However, SDMs are always simplifications and the resulting projections need to be interpreted with caution.

To improve SDMs, future studies should try to consider additional factors, such as biotic interactions (Schleuning *et al*., 2020) and the reshuffling of species communities (Voskamp *et al*., in prep), which might lead to a change in competitive balance (Ockendon *et al*., 2014), altered predator-prey relationships (Harley, 2011) or changes in functional diversity (*Stewart et al., 2022*) and thus the provision of ecosystem functions and services (Pecl *et al*., 2017). While the above factors might improve climate change impact projections in the future, such modelling exercises will never reflect the truth, as a species’ response to climate change will be strongly influenced by behaviour, diel rhythm, thermoregulatory potential and microclimatic conditions (Anderson *et al*., 2022). This complexity highlights the need for integrative approaches when investigating species’ responses to climate change (Hof, 2021).

## Conclusion

Our study shows that reptiles are not only likely to be impacted by future climate change, globally but also within most zoogeographic realms. These impacts are projected to have a considerable effect on the extent and location of species’ geographic ranges. Thus, to prevent large scale declines in reptile species, not only is it of key importance to lower CO_2_ emissions in order to stop on-going climate change, but also to maintain adequate habitats of sufficient size and quality, especially grassland and savanna habitats (Roll *et al*., 2017). Furthermore, it is necessary to establish new protected areas that will help to prevent the extinction of particularly vulnerable species, i.e. by establishing high-elevation climate refugia within current species ranges (Sinervo *et al*., 2018).

## Data Accessibility Statement

GARD range maps are available from https://doi.org/10.5061/dryad.83s7k. EWEMBI and ISIMIP2b climate data are available from https://data.isimip.org/10.5880/pik.2019.004 and https://data.isimip.org/search/query/ISIMIP2b%20Input/tree/ISIMIP2b/InputData/climate/atmosphere/. The code for creating the species distribution models can be found at https://github.com/christianhof/BioScen1.5_SDM, whereas the code and data for the performed data analysis and the presented figures, can be found on Dryad (doi: 10.5061/dryad.rn8pk0pgb; Biber et al. 2023).

## Supplementary Material

Appendix S1 Methodological details

Appendix S2 Taxon-specific results

Appendix S3 Additional range change results

Appendix S4 Summed probability results

Appendix S5 Variation across dispersal distances

Appendix S6 Variation across RCPs and years

