## Supporting Information for "Potential future climate change effects on global reptile distributions and diversity"

### Supporting Information for: 'Potential future climate change effects on global reptile distribution and diversity'

#### Appendix S1 Methodological details

**Table S1.1.** Description of the 19 bioclimatic variables used for variable selection in our modelling approach, calculated with the biovars() function from the `dismo` package in R. Bioclimatic variables are derived from monthly minimum and maximum temperature as well as precipitation data in order to generate more biologically meaningful variables. The bioclimatic variables represent annual trends, seasonality and extreme or limiting environmental factors. A quarter is a period of three months (1/4 of the year).

| **Variable abbreviation** | **Variable description** |
|---|---|
| BIO1 | Annual Mean Temperature |
| BIO2 | Mean Diurnal Range (Mean of monthly (max temp - min temp)) |
| BIO3 | Isothermality (BIO2/BIO7) (×100) |
| BIO4 | Temperature Seasonality (standard deviation ×100) |
| BIO5 | Max Temperature of Warmest Month |
| BIO6 | Min Temperature of Coldest Month |
| BIO7 | Temperature Annual Range (BIO5-BIO6) |
| BIO8 | Mean Temperature of Wettest Quarter |
| BIO9 | Mean Temperature of Driest Quarter |
| BIO10 | Mean Temperature of Warmest Quarter |
| BIO11 | Mean Temperature of Coldest Quarter |
| BIO12 | Annual Precipitation |
| BIO13 | Precipitation of Wettest Month |
| BIO14 | Precipitation of Driest Month |
| BIO15 | Precipitation Seasonality (Coefficient of Variation) |
| BIO16 | Precipitation of Wettest Quarter |
| BIO17 | Precipitation of Driest Quarter |
| BIO18 | Precipitation of Warmest Quarter |
| BIO19 | Precipitation of Coldest Quarter |

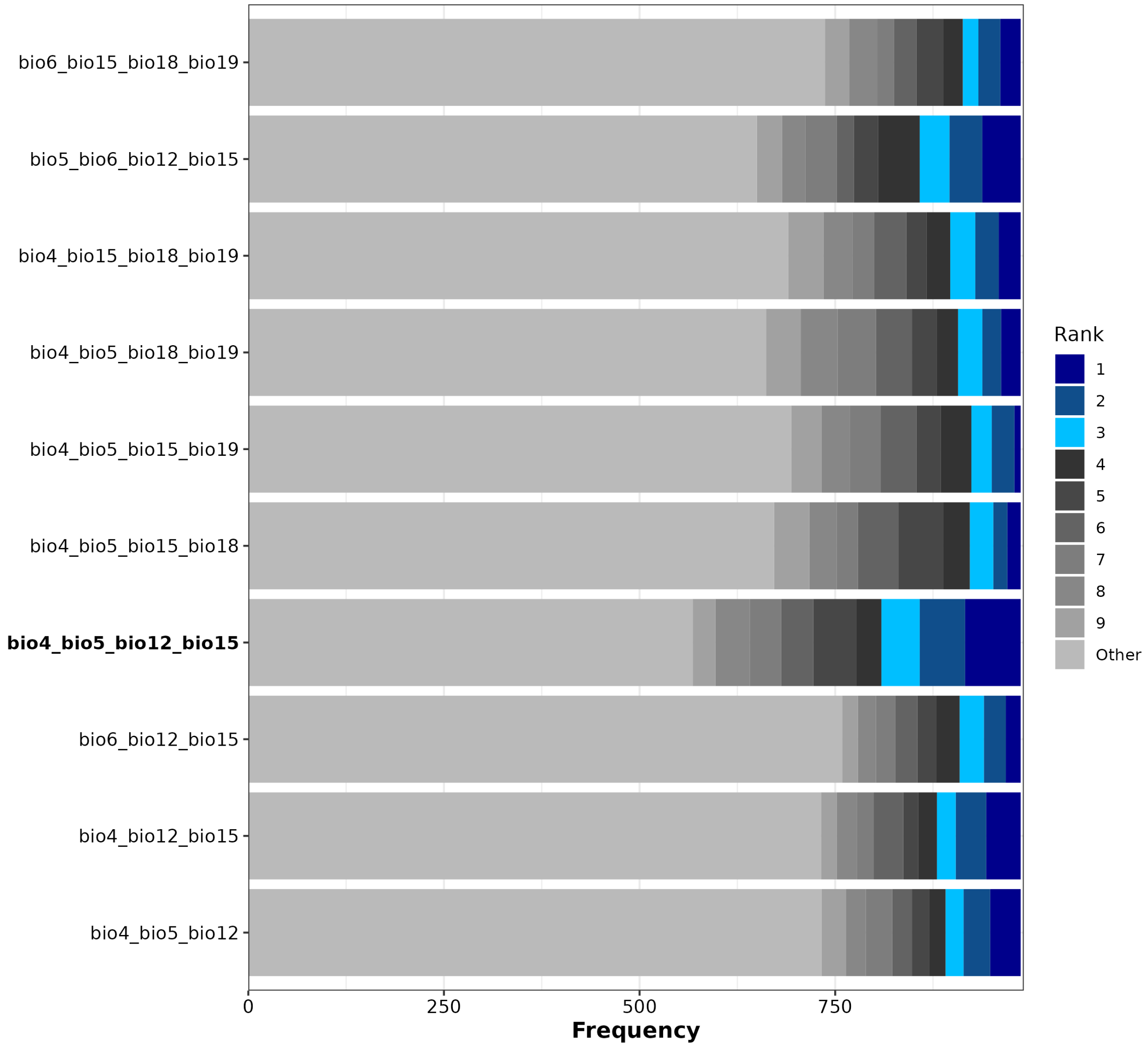


**Figure S1.1.** Various (Top10) non-correlated combinations of three and four bioclimatic variables and their rank of model performance across a representative subset of species (see Hof et al. (2018) for a detailed description of the variable selection method). The optimal variable combination, used for the models presented in this study, is highlighted in bold.

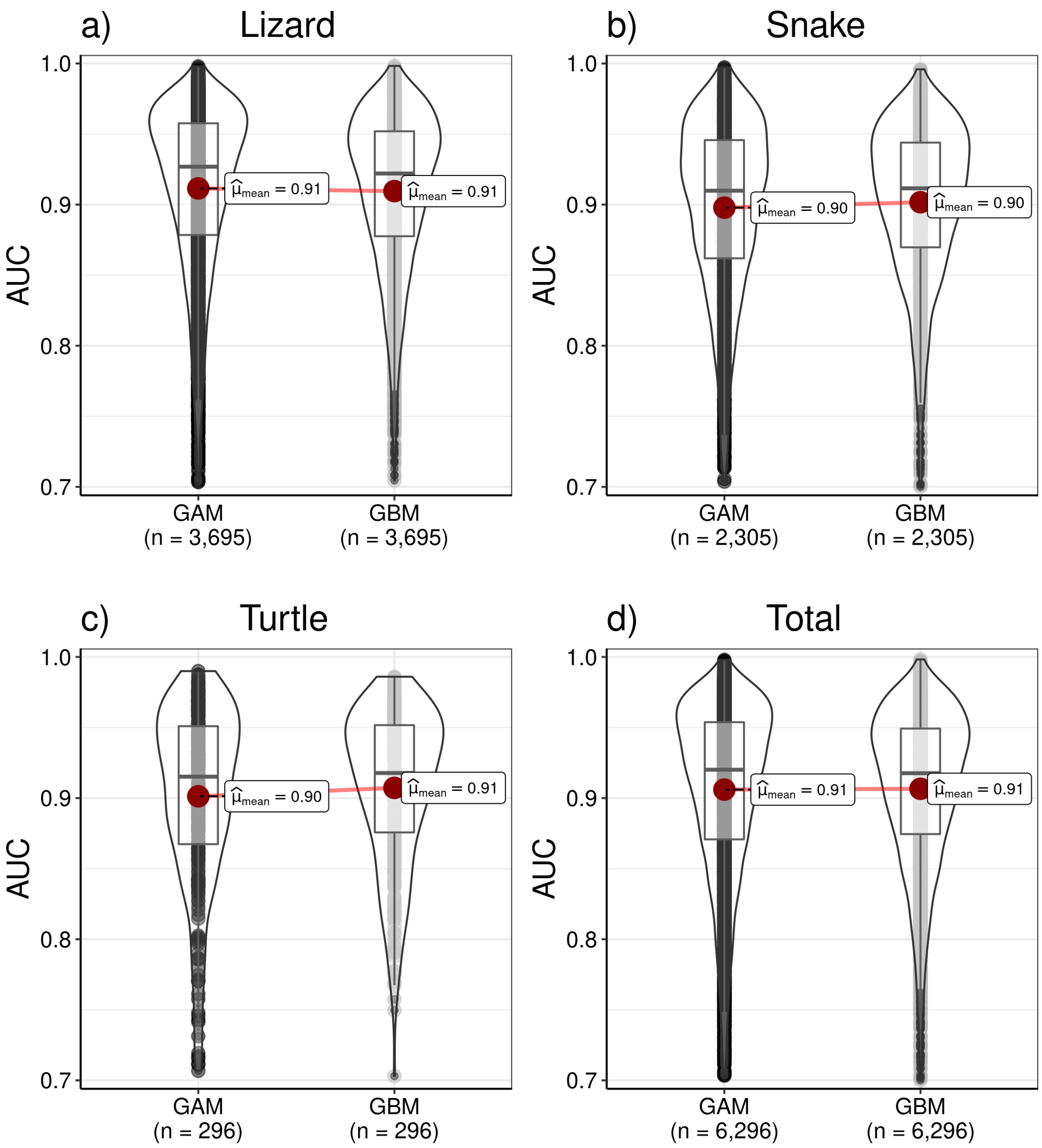


**Figure S1.2.** Model performance (AUC) for the different reptile groups (a) lizard, b) snake, c), turtle, d) total) for each of the two model algorithms (GAM, GBM) used. AUC values are only shown for models used in the analysis, hence only values >= 0.7 are shown, as species which had models with a lower performance were removed from the analysis. See Table 1 for an overview on the number of species removed due to a low AUC. The statistical difference between GAM and GBM models was tested using a paired Student's t-test with Holm correction ($p < 0.05$ = *, $p < 0.01$ = **, $p < 0.001$ = ***, $p < 0.0001$ = ****). Plots show mean (red point & label), median (black horizontal line), 25th to 75th percentiles (box), entire range of data (violin & data points) and density of values (width of violin).

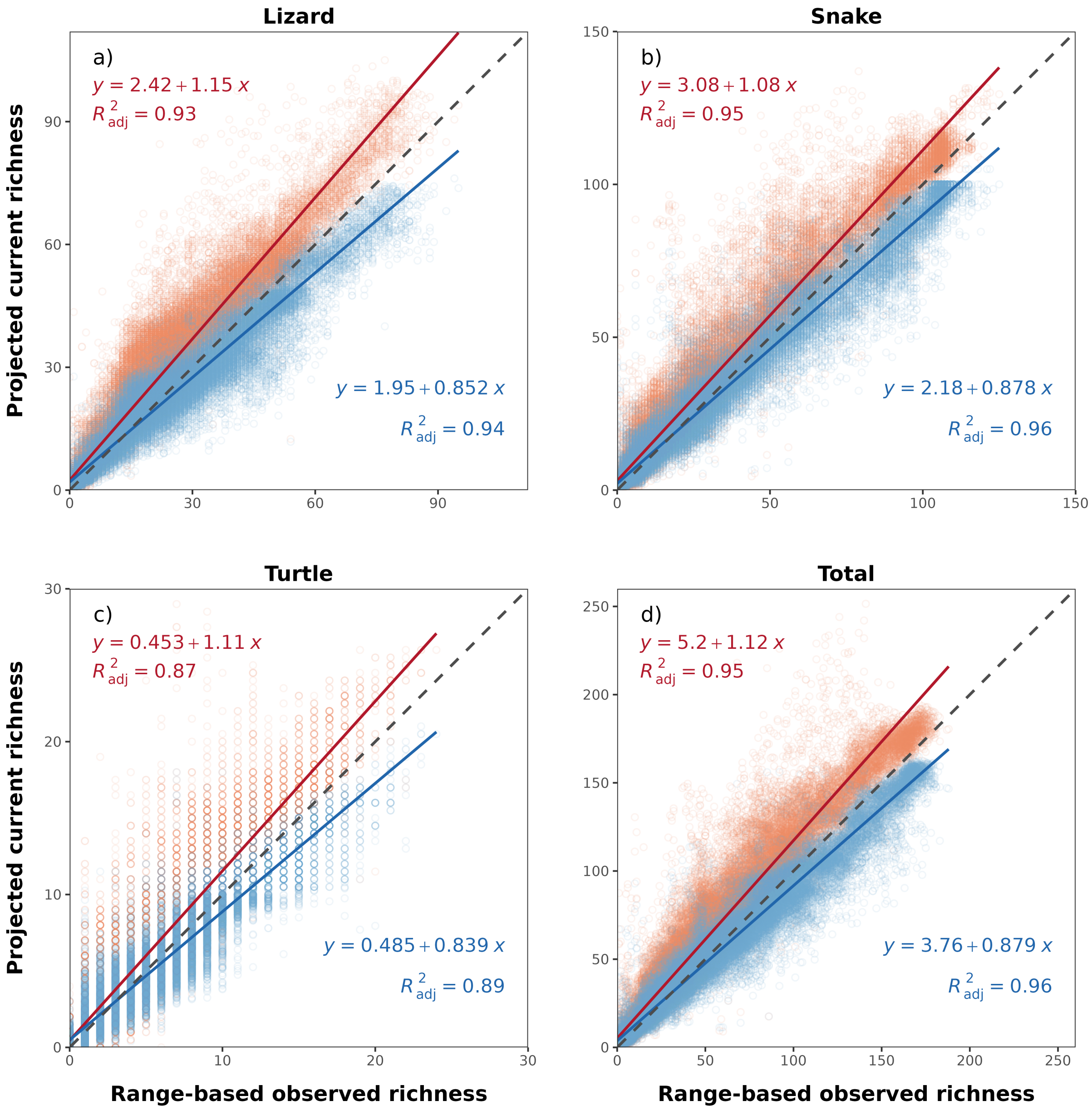


**Figure S1.3.** Range-based observed species richness (gridded to 0.5°) versus projected current species richness for each grid cell per taxon (a) lizard, b) snake, c) turtle and d) total) for ensemble models (average among model algorithms). Thresholded sum of species occurrence per grid cell is shown in red, while the sum of species probability per grid cell is shown in blue. Stacked species distribution models (S-SDMs) were performed using a medium dispersal scenario (d/8). Perfect fits would have intercepts of 0, slopes of 1 and a R2 value of 1 (dashed grey line).

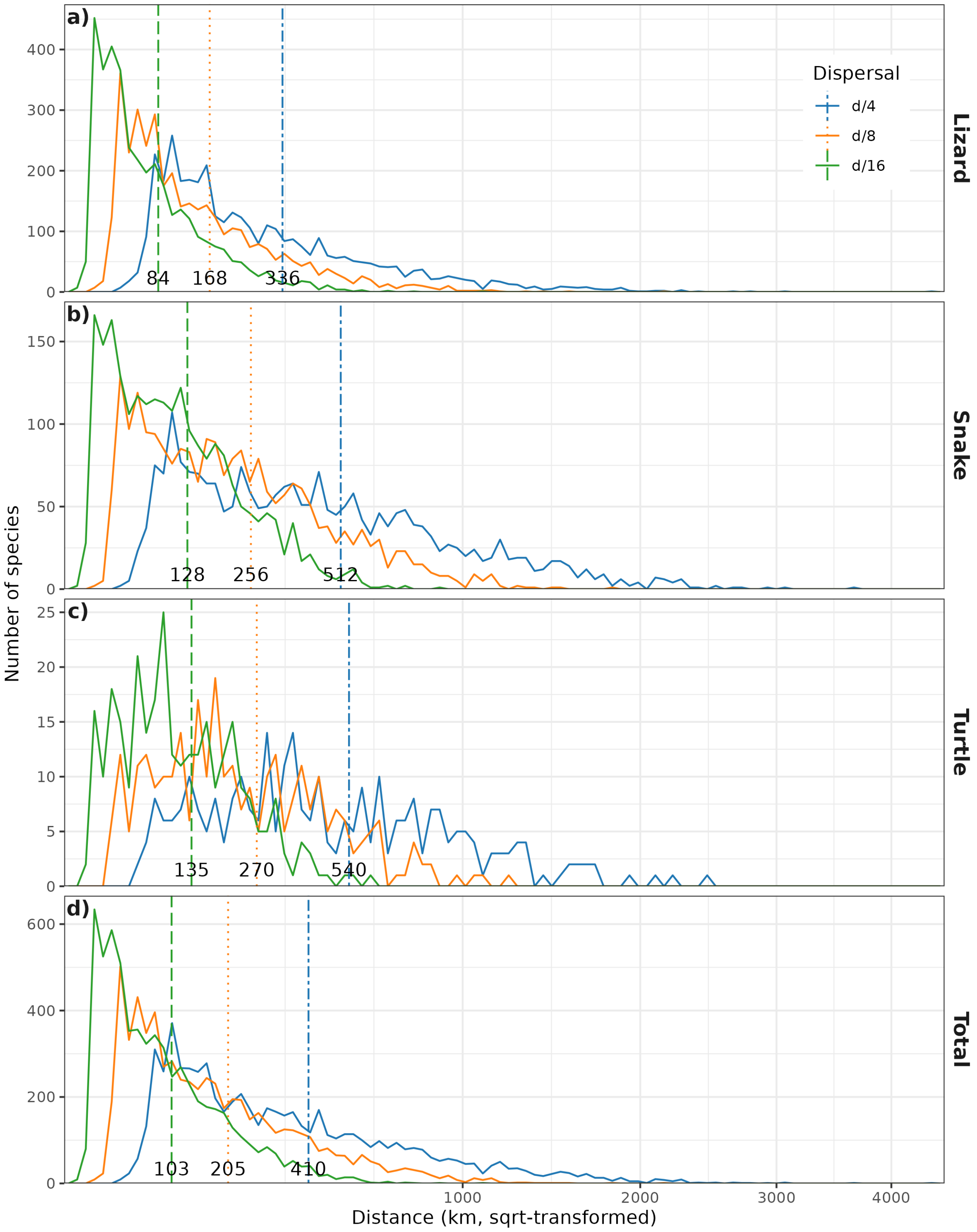


**Figure S1.4.** Frequency plot of dispersal distance for all a) lizard, b) snake, c) turtle and d) total reptile species under the three different dispersal scenarios considered (d/4, d/8 and d/16). Vertical lines and values represent mean dispersal distance for the respective dispersal scenarios.

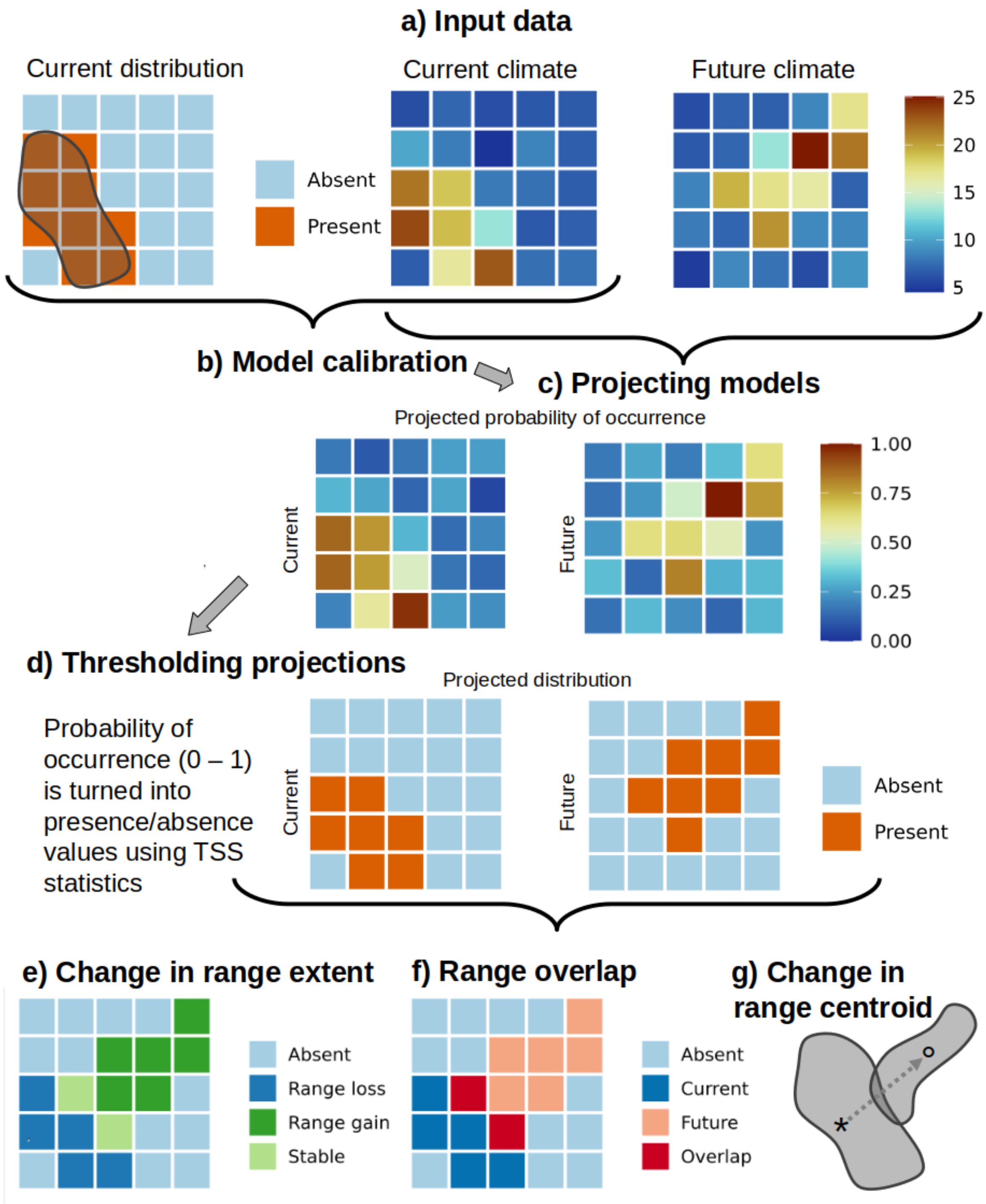


**Figure S1.5.** Schematic overview of the different methodological steps of our analyses. a) We prepared distribution data and current and future climate data in order to b) calibrate our species distribution models (SDMs). Based on these models, we c) created projections for current and future conditions (1995 and 2050/2080). In order to perform our analyses, we then d) thresholded the projected probability of occurrence into binary (presence/absence) projected distribution data. In the subsequent steps (e - g), we perform different species-specific analyses to infer information about e) the change in range extent, f) the range overlap between current and future projected distribution and g) the change in range position by assessing the direction and distance of change in range centroid.

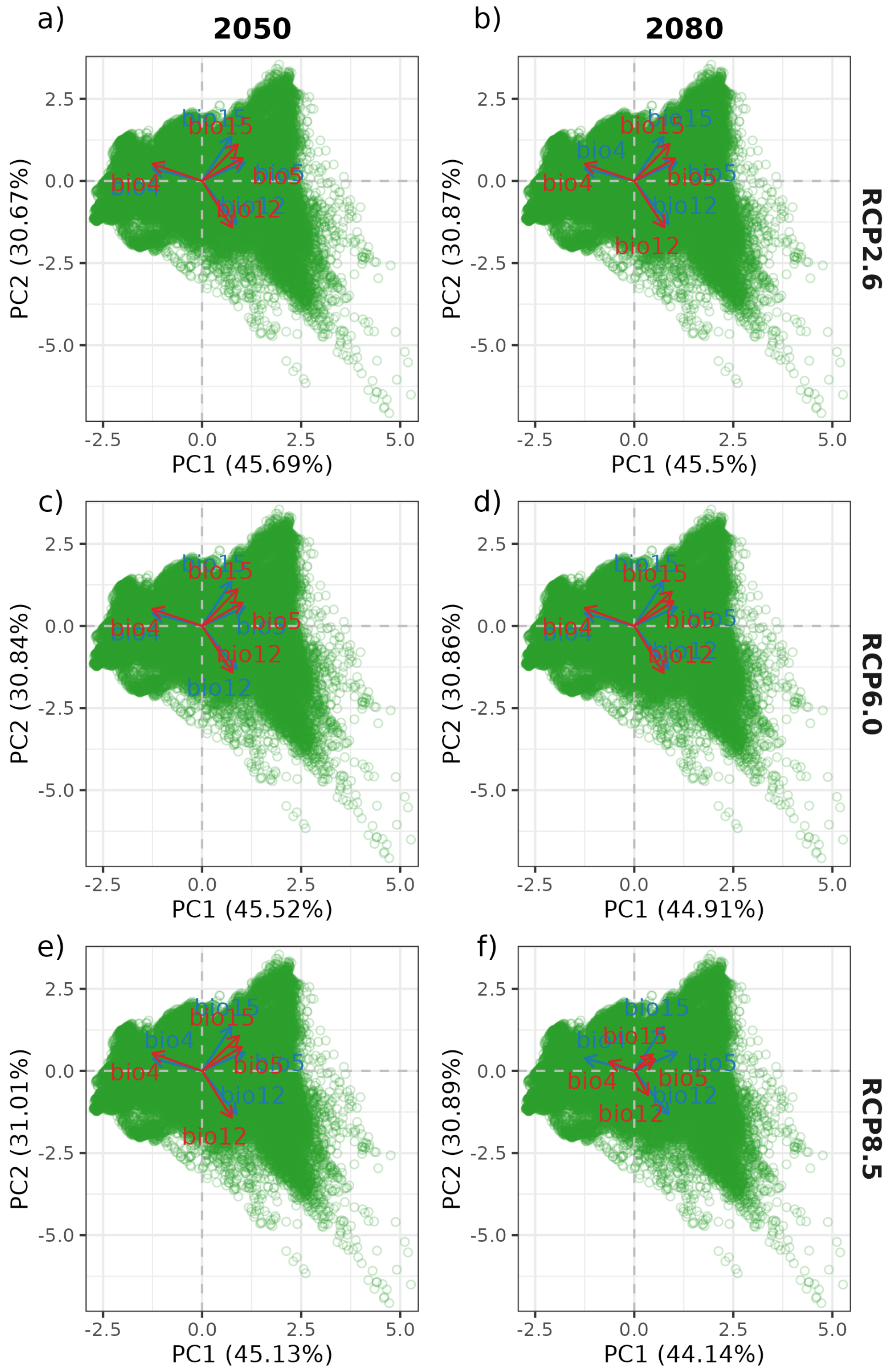


**Figure S1.6.** Principal component analysis of bioclimatic variables used for species distribution models (bio4, bio5, bio12, bio15) for current conditions (green dots and blue arrows) and future conditions (yellow dots and red arrows). Different panels show variation across years (a),c),e) 2050, b),d),f) 2080) and representative concentration pathways (a),b) RCP2.6, c),d) RCP6.0, e),f) RCP8.5). Explained variance (%) of PCA axes for future conditions are presented in the axis titles. Explained variance of PCA axes for current conditions was 46.9% for PC1 and 29.8% for PC2.

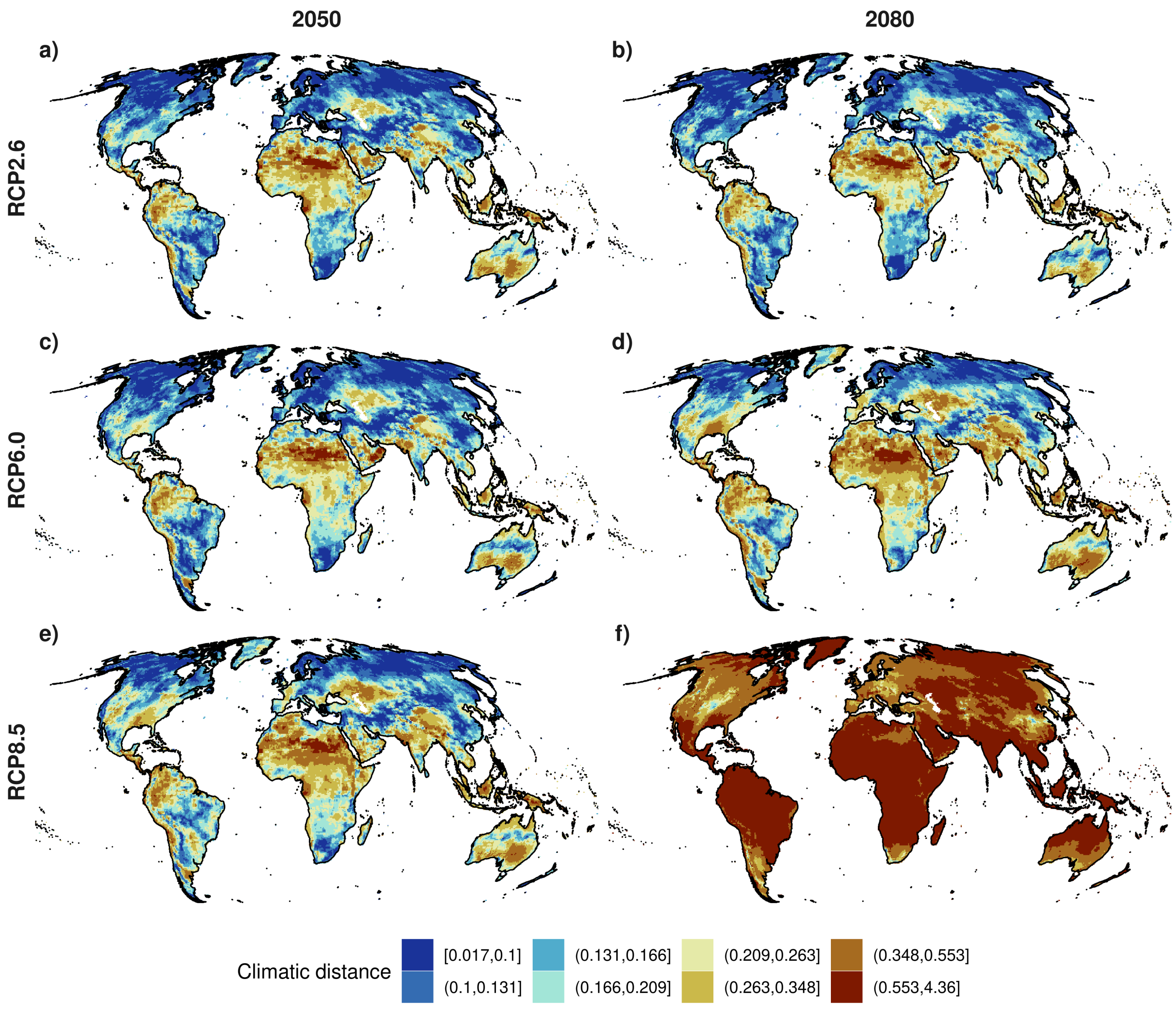


**Figure S1.7.** Climatic distance (Euclidean distance between current and future principal components (PC1 & PC2, see Fig. S32)) for the years 2050 and 2080 under the different representative concentration pathways (a),b) RCP2.6, c),d) RCP6.0, e),f) RCP8.5). The color key is divided into 10 quantile intervals. The results are presented as the ensemble mean, across the four global circulation models (GCMs). All maps are based on 0.5° x 0.5° grid cells, which have been projected to Mollweide equal-area projection (EPSG:54009).

#### Appendix S2 Taxon-specific results

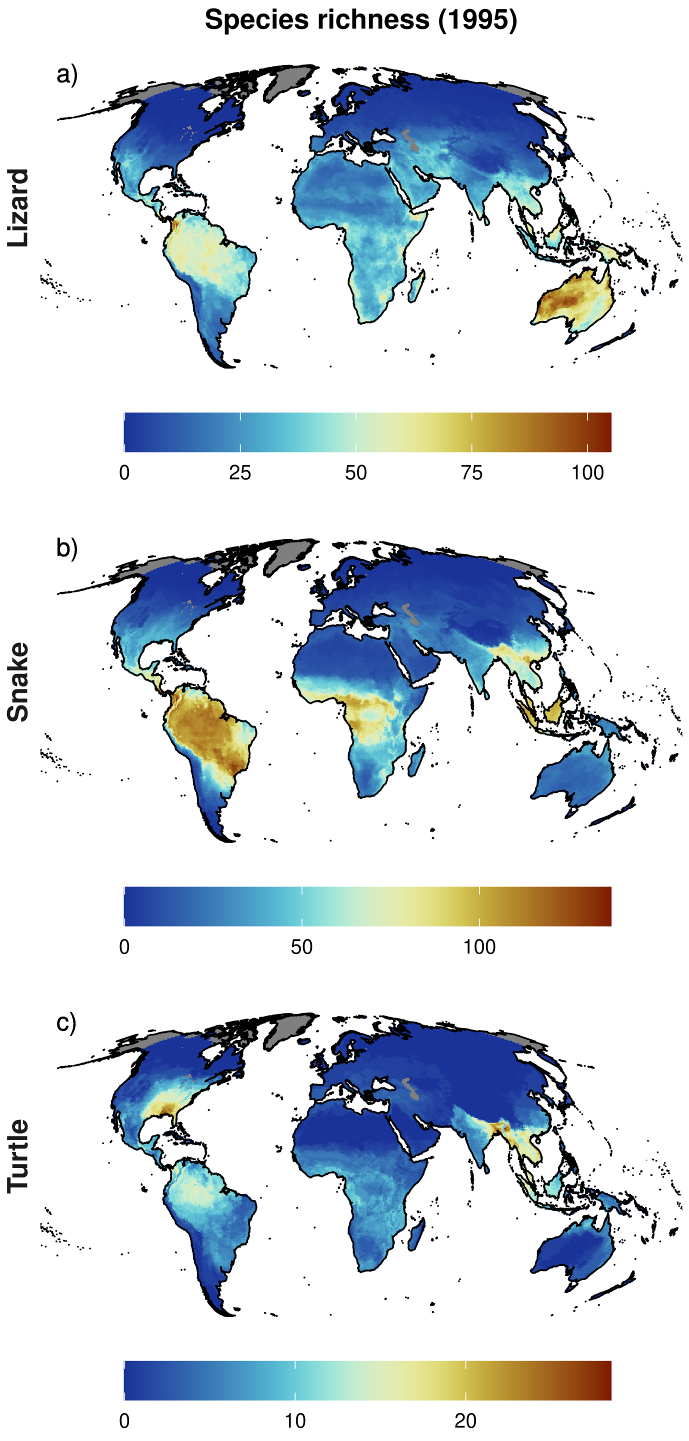


**Figure S2.8.** Projected species richness for the three taxa (a) lizards, b) snakes and c) turtles) for current conditions (1995) under the dispersal scenario d/8. The color key is divided into 10 quantile intervals. The results are presented as the ensemble mean, across the two model algorithms (GAM & GBM) considered. All maps are based on 0.5° x 0.5° grid cells, which have been projected to Mollweide equal-area projection (EPSG:54009). Grey areas are regions for which no projections are available.

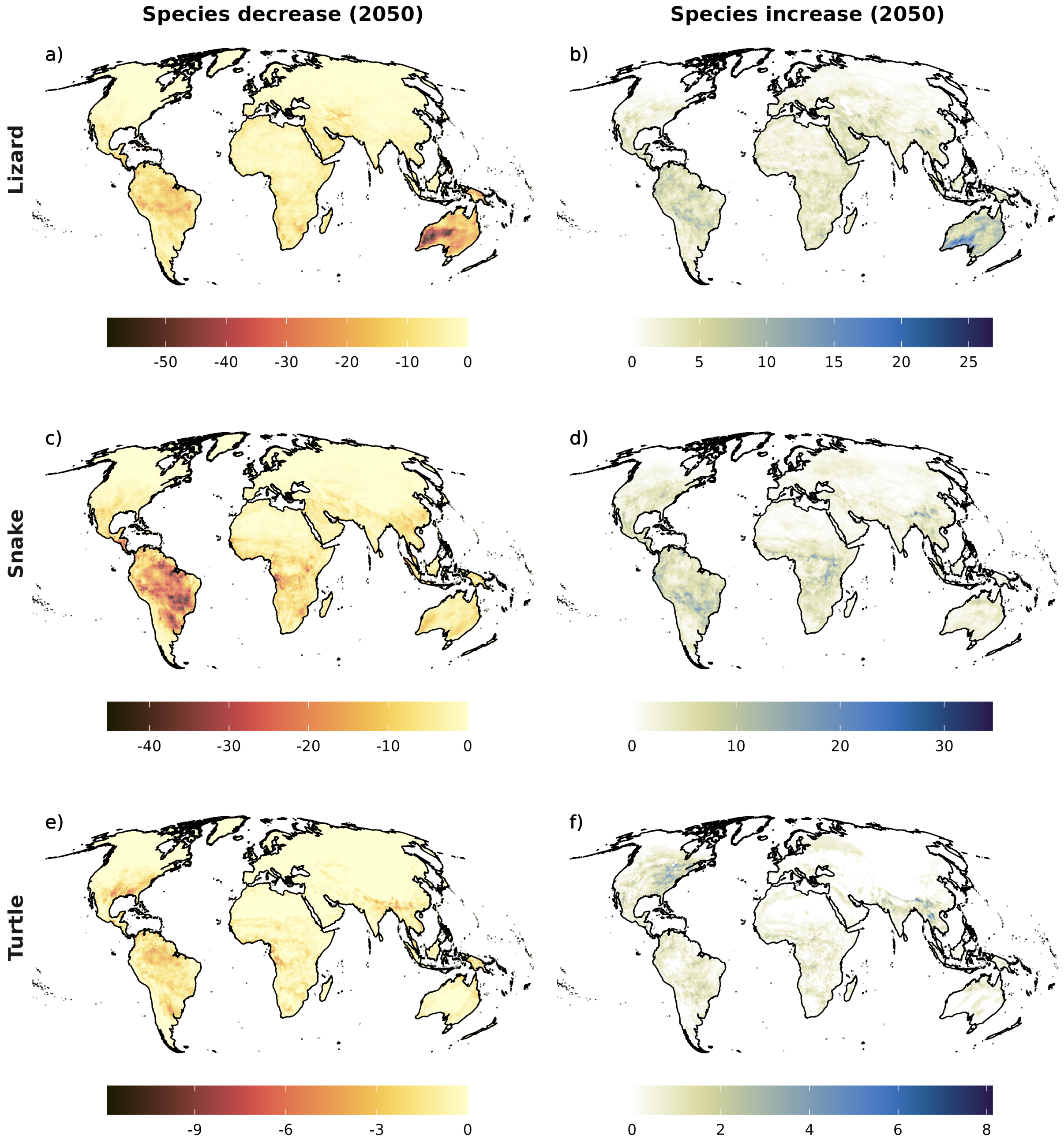


**Figure S2.9.** Potential species decrease (a),c),e)) and increase (b),d),f)) for the three taxa (lizards, snakes, turtles). The results are presented as the ensemble mean, across the four global circulation models (GCMs) and two model algorithms (GAM & GBM) considered, for the year 2050 under a medium representative concentration pathway (RCP6.0) and a medium dispersal scenario (d/8). All maps are based on 0.5° x 0.5° grid cells, which have been projected to Mollweide equal-area projection (EPSG:54009). Grey areas are regions for which no projections are available.

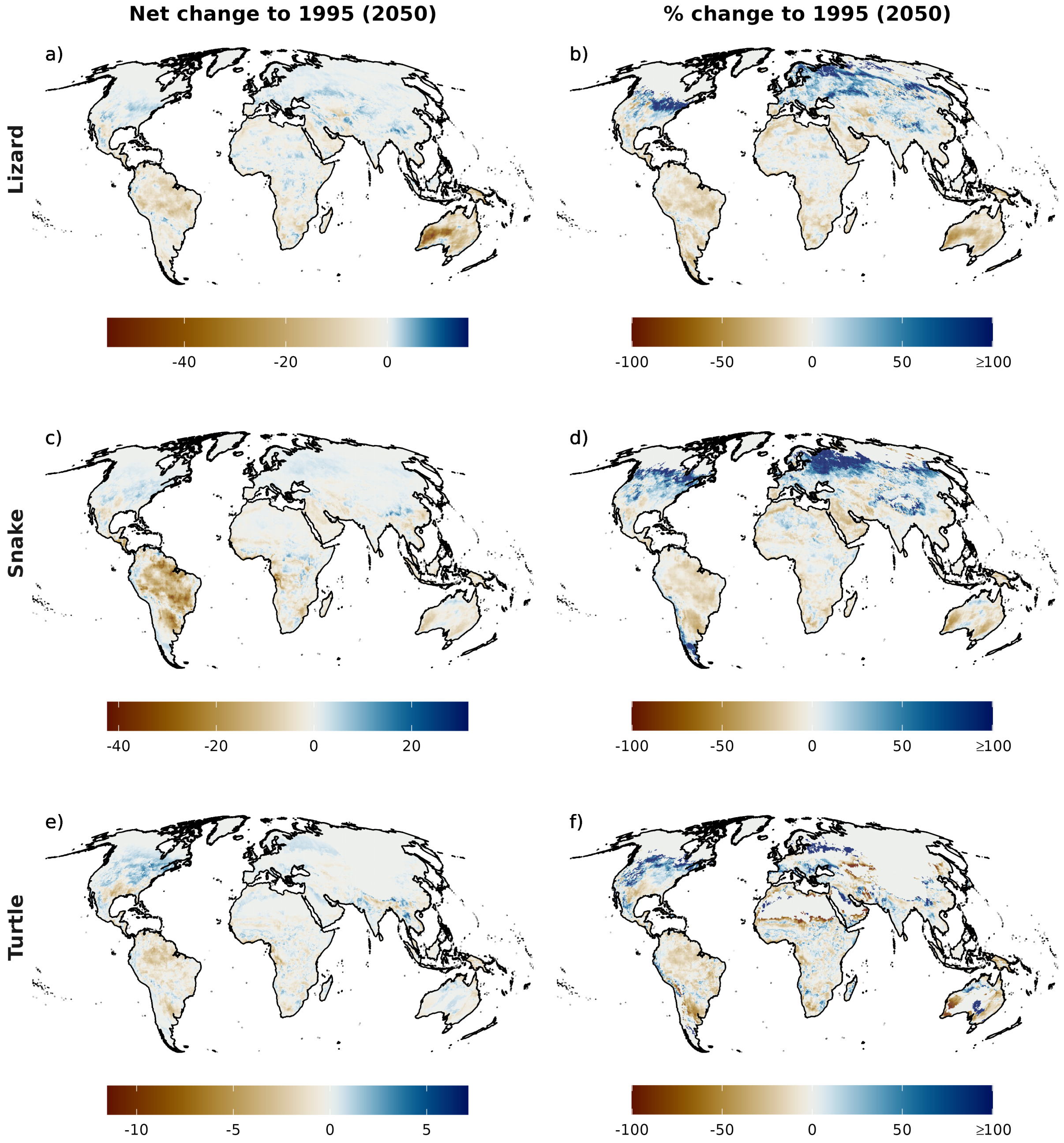


**Figure S2.10.** Potential net change (a),c),e)) and relative change (b),d),f)) in species richness for the three taxa (lizards, snakes, turtles). The results are presented as the ensemble mean, across the four global circulation models (GCMs) and two model algorithms (GAM & GBM) considered, for the year 2050 under a medium representative concentration pathway (RCP6.0) and a medium dispersal scenario (d/8). All maps are based on 0.5° x 0.5° grid cells, which have been projected to Mollweide equal-area projection (EPSG:54009). Grey areas are regions for which no projections are available.

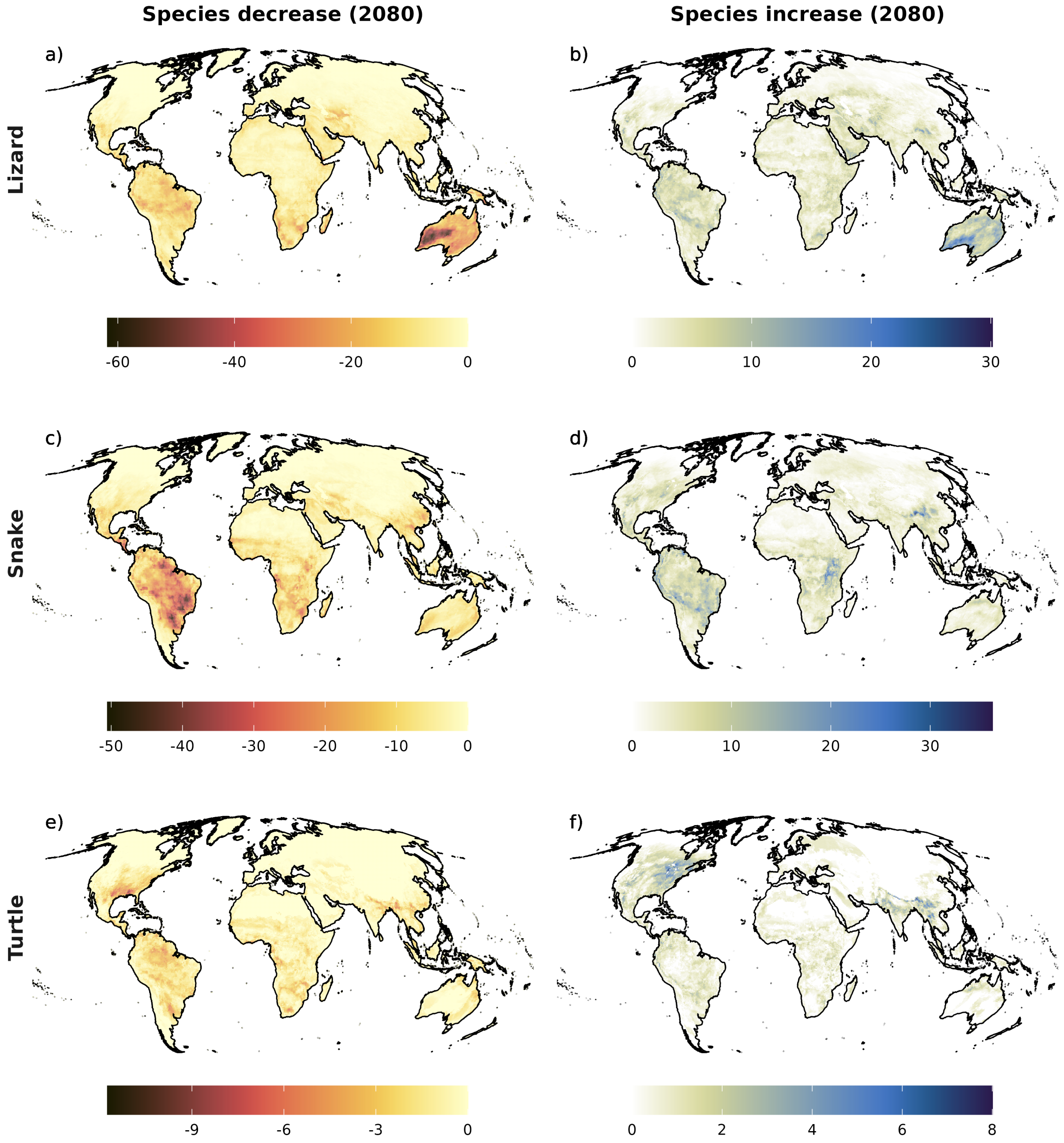


**Figure S2.11.** Potential species decrease (a),c),e)) and increase (b),d),f)) for the three taxa (lizards, snakes, turtles). The results are presented as the ensemble mean, across the four global circulation models (GCMs) and two model algorithms (GAM & GBM) considered, for the year 2080 under a medium representative concentration pathway (RCP6.0) and a medium dispersal scenario (d/8). All maps are based on 0.5° x 0.5° grid cells, which have been projected to Mollweide equal-area projection (EPSG:54009). Grey areas are regions for which no projections are available.

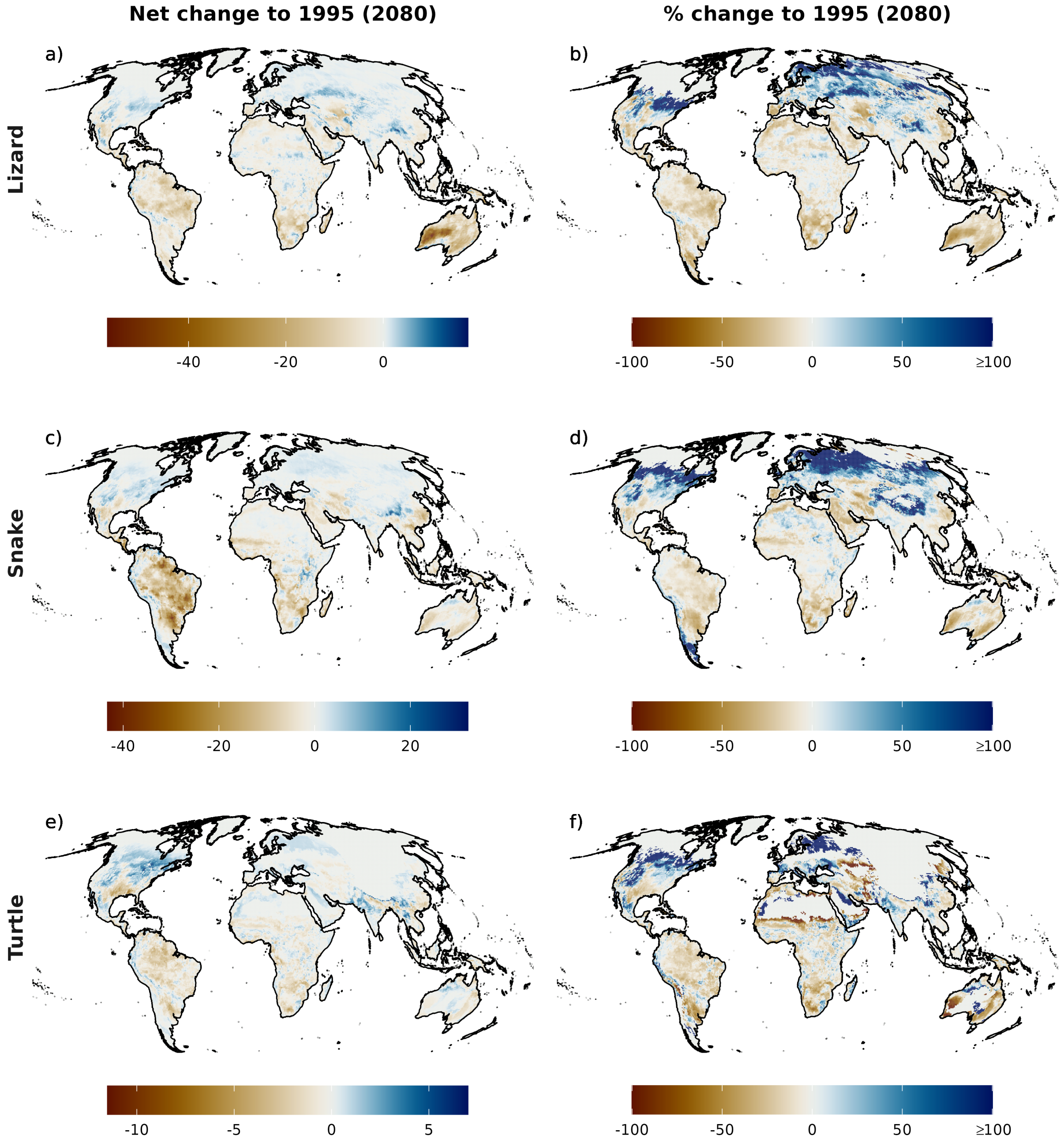


**Figure S2.12.** Potential net change (a),c),e)) and relative change (b),d),f)) in species richness for the three taxa (lizards, snakes, turtles). The results are presented as the ensemble mean, across the four global circulation models (GCMs) and two model algorithms (GAM & GBM) considered, for the year 2050 under a medium representative concentration pathway (RCP6.0) and a medium dispersal scenario (d/8). All maps are based on 0.5° x 0.5° grid cells, which have been projected to Mollweide equal-area projection (EPSG:54009). Grey areas are regions for which no projections are available.

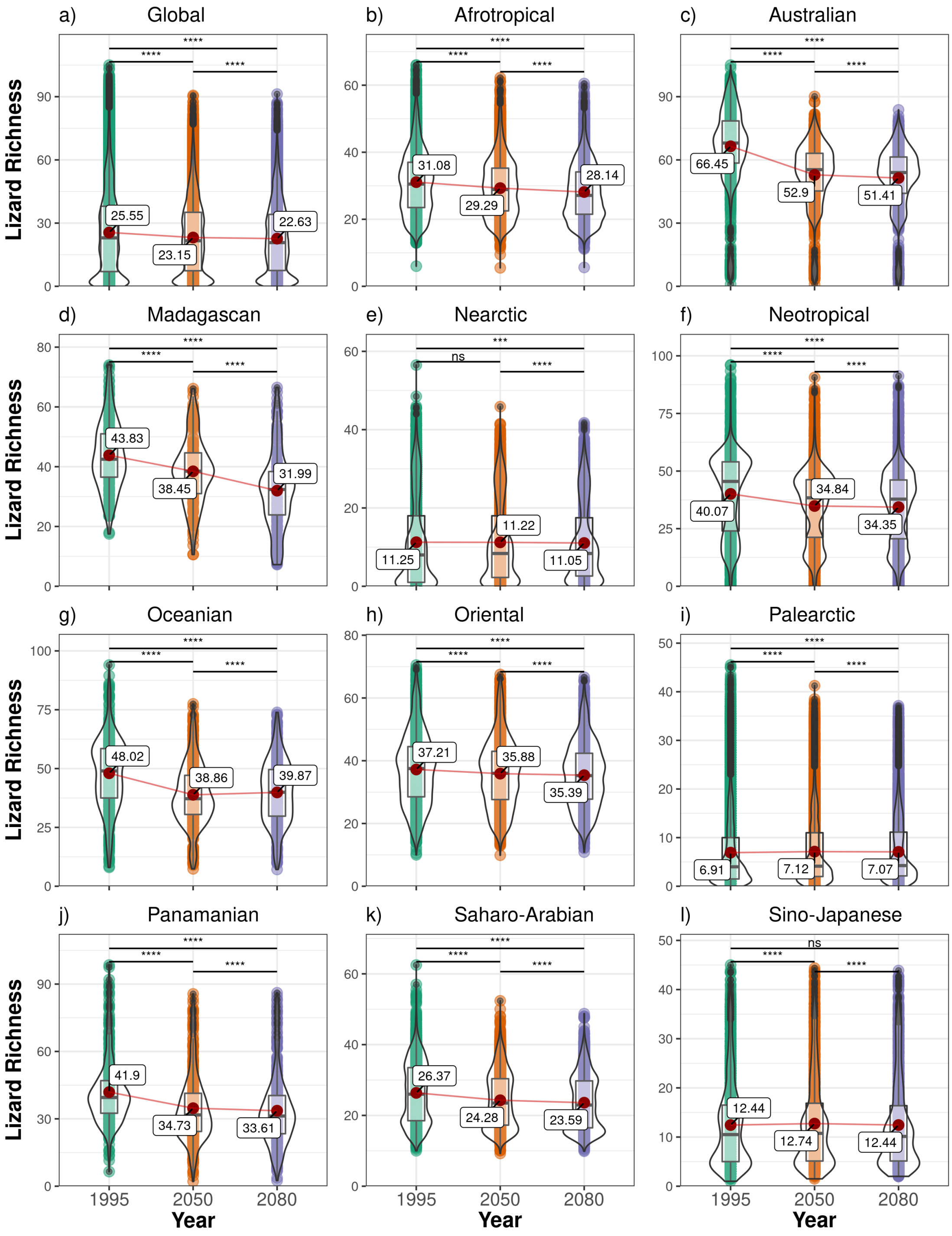


**Figure S2.13.** Terrestrial lizard richness across the globe and for each zoogeographic realm over time (1995, 2050, 2080) based on all modelled lizard species (N = 3695). The results are presented as the ensemble mean, across the four global circulation models (GCMs) and two model algorithms (GAM & GBM) considered, under a medium representative concentration pathway (RCP6.0) and a medium dispersal scenario (d/8). The statistical difference between years was tested using a paired Student's t-test with Holm correction ($p < 0.05$ = *, $p < 0.01$ = **, $p < 0.001$ = ***, $p < 0.0001$ = ****). Plots show mean (red point & label), median (black horizontal line), 25th to 75th percentiles (box), entire range of data (violin & data points) and density of values (width of violin).

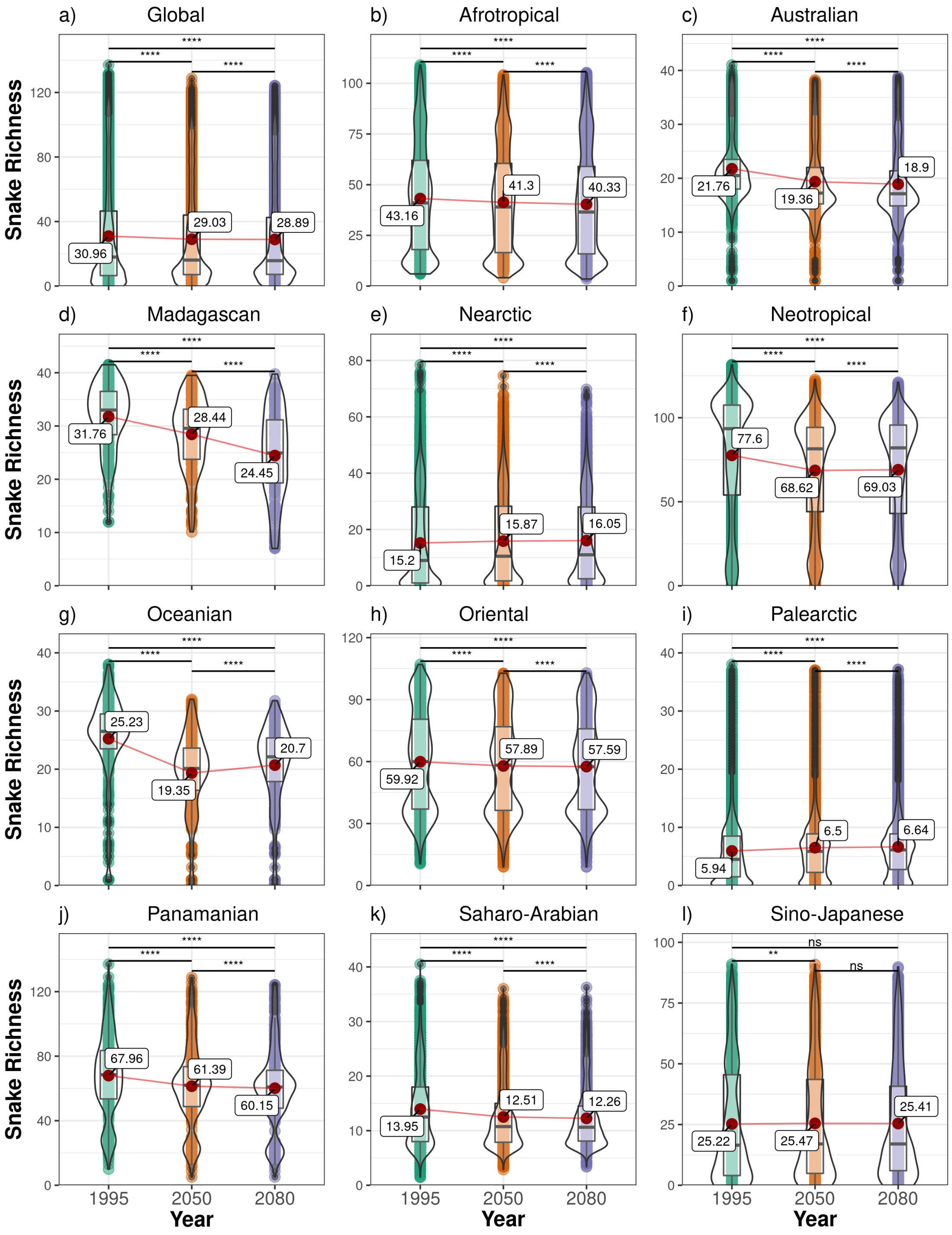


**Figure S2.14.** Terrestrial snake richness across the globe and for each zoogeographic realm over time (1995, 2050, 2080) based on all modelled snake species (N = 2305). The results are presented as the ensemble mean, across the four global circulation models (GCMs) and two model algorithms (GAM & GBM) considered, under a medium representative concentration pathway (RCP6.0) and a medium dispersal scenario (d/8). The statistical difference between years was tested using a paired Student's t-test with Holm correction ($p < 0.05$ = *, $p < 0.01$ = **, $p < 0.001$ = ***, $p < 0.0001$ = ****). Plots show mean (red point & label), median (black horizontal line), 25th to 75th percentiles (box), entire range of data (violin & data points) and density of values (width of violin).

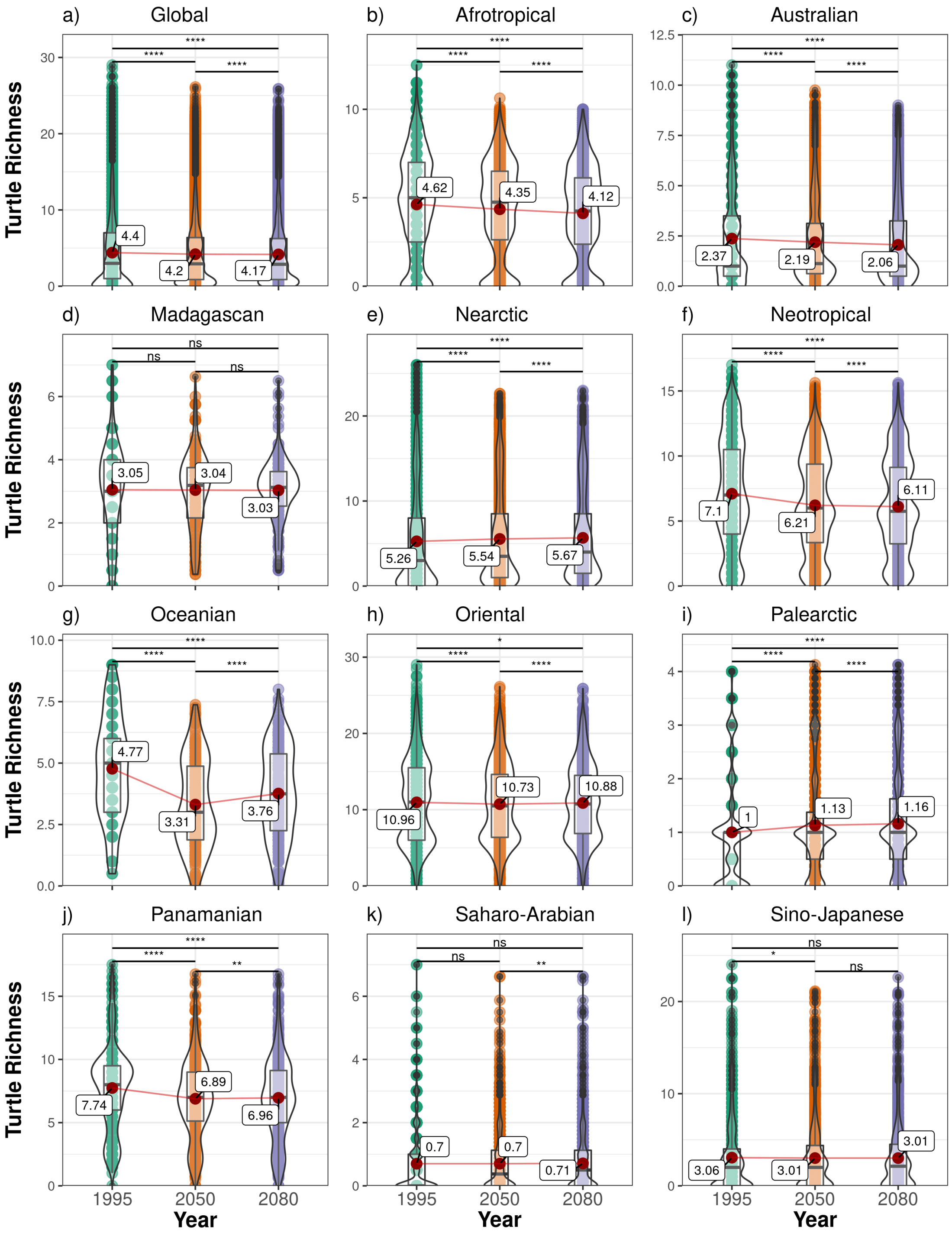


**Figure S2.15.** Terrestrial turtle richness across the globe and for each zoogeographic realm over time (1995, 2050, 2080) based on all modelled turtle species (N = 296). The results are presented as the ensemble mean, across the four global circulation models (GCMs) and two model algorithms (GAM & GBM) considered, under a medium representative concentration pathway (RCP6.0) and a medium dispersal scenario (d/8). The statistical difference between years was tested using a paired Student's t-test with Holm correction ($p < 0.05$ = *, $p < 0.01$ = **, $p < 0.001$ = ***, $p < 0.0001$ = ****). Plots show mean (red point & label), median (black horizontal line), 25th to 75th percentiles (box), entire range of data (violin & data points) and density of values (width of violin).

#### Appendix S3 Additional range change results

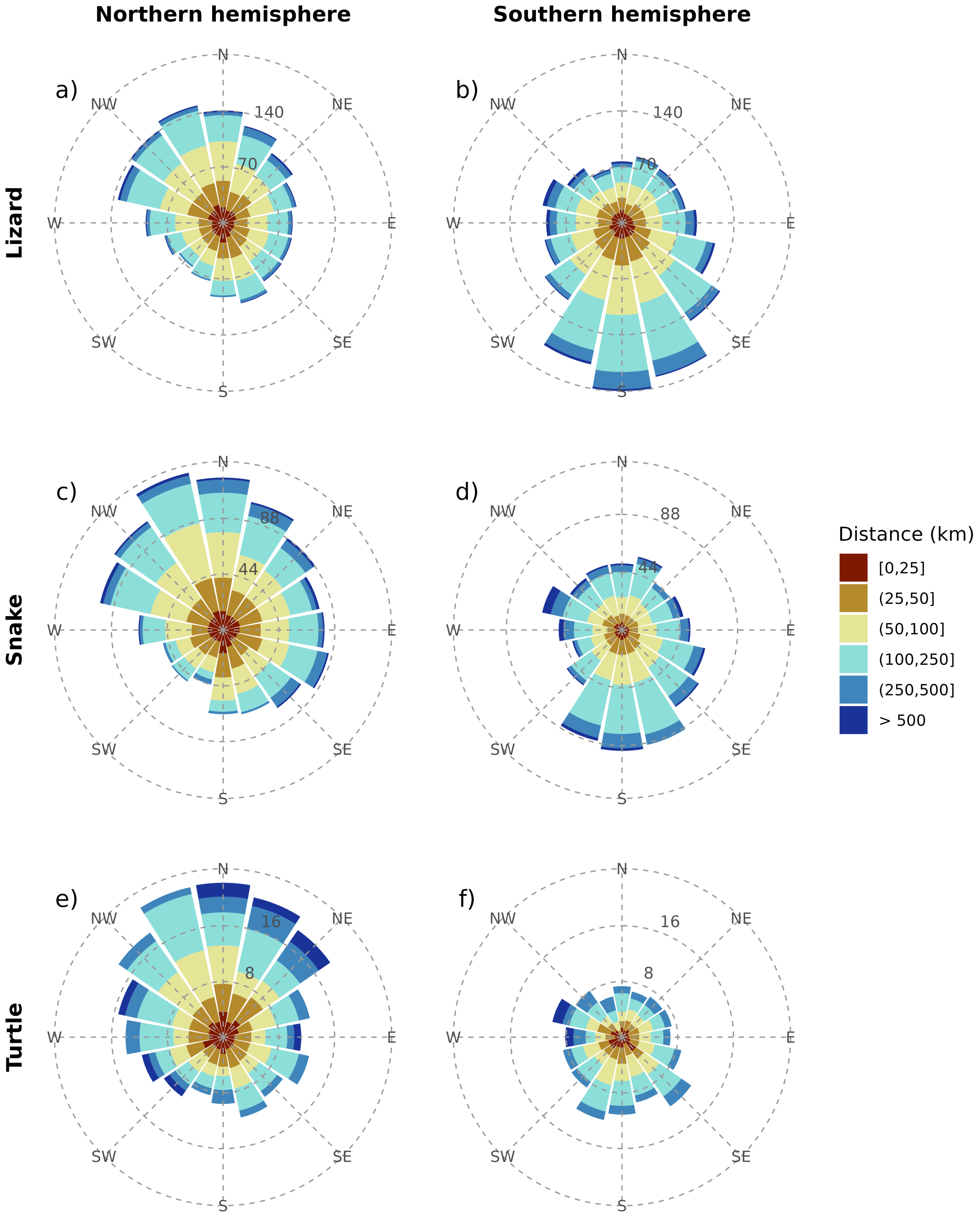


**Figure S3.16.** Cumulative direction and distance of potential range centroid changes per taxonomic group (lizard, snake, turtle) separated by northern and southern hemisphere. The results are presented as the ensemble mean, across the four global circulation models (GCMs) and two model algorithms (GAM & GBM) considered, for the year 2080 under a medium representative concentration pathway (RCP6.0) and a medium dispersal scenario (d/8).

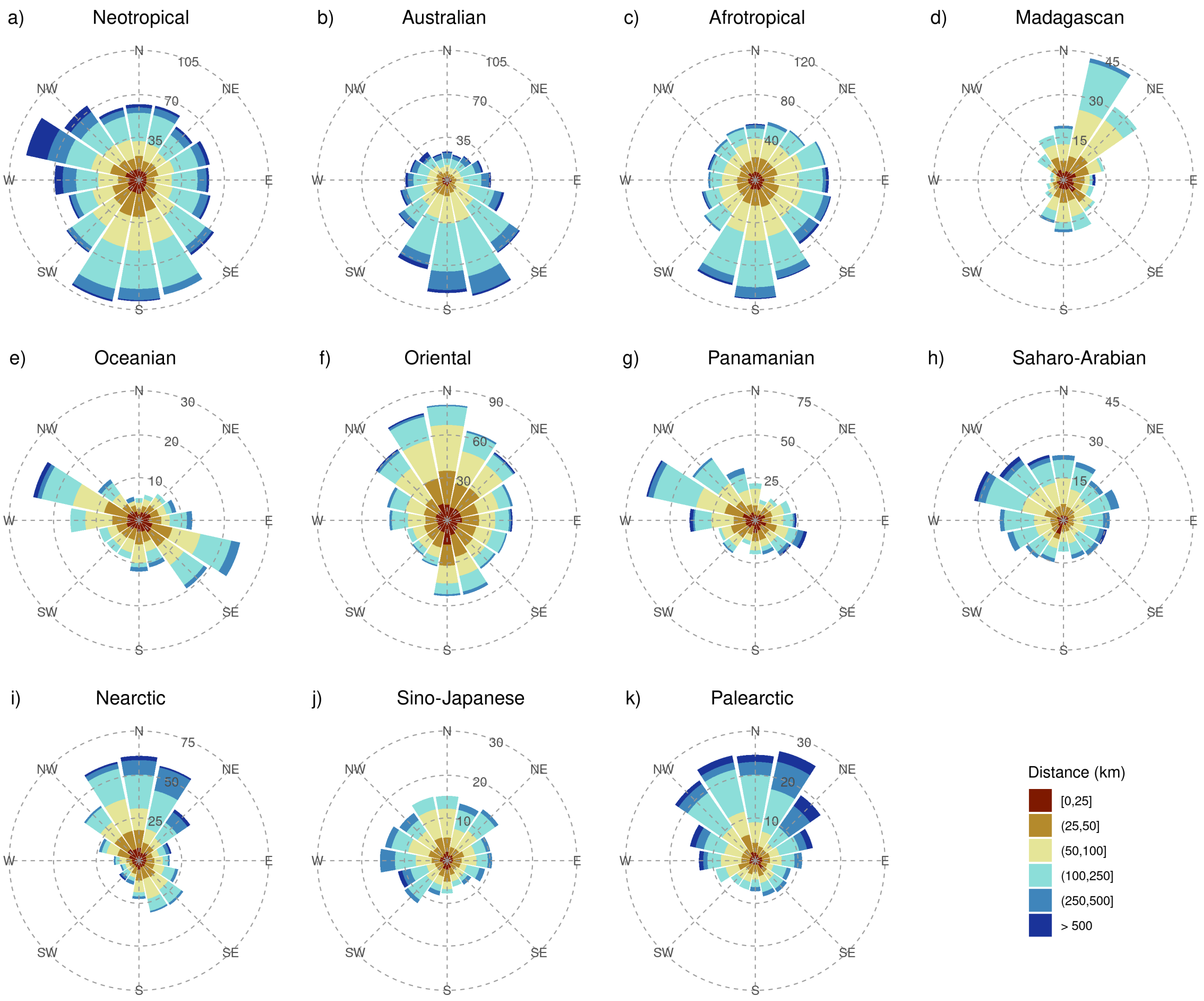


**Figure S3.17.** Cumulative direction and distance of potential range changes in total reptiles per zoogeographic realm. The results are presented as the ensemble mean, across the four global circulation models (GCMs) and two model algorithms (GAM & GBM) considered, for the year 2080 under a medium representative concentration pathway (RCP6.0) and a medium dispersal scenario (d/8).

#### Appendix S4 Summed probability results

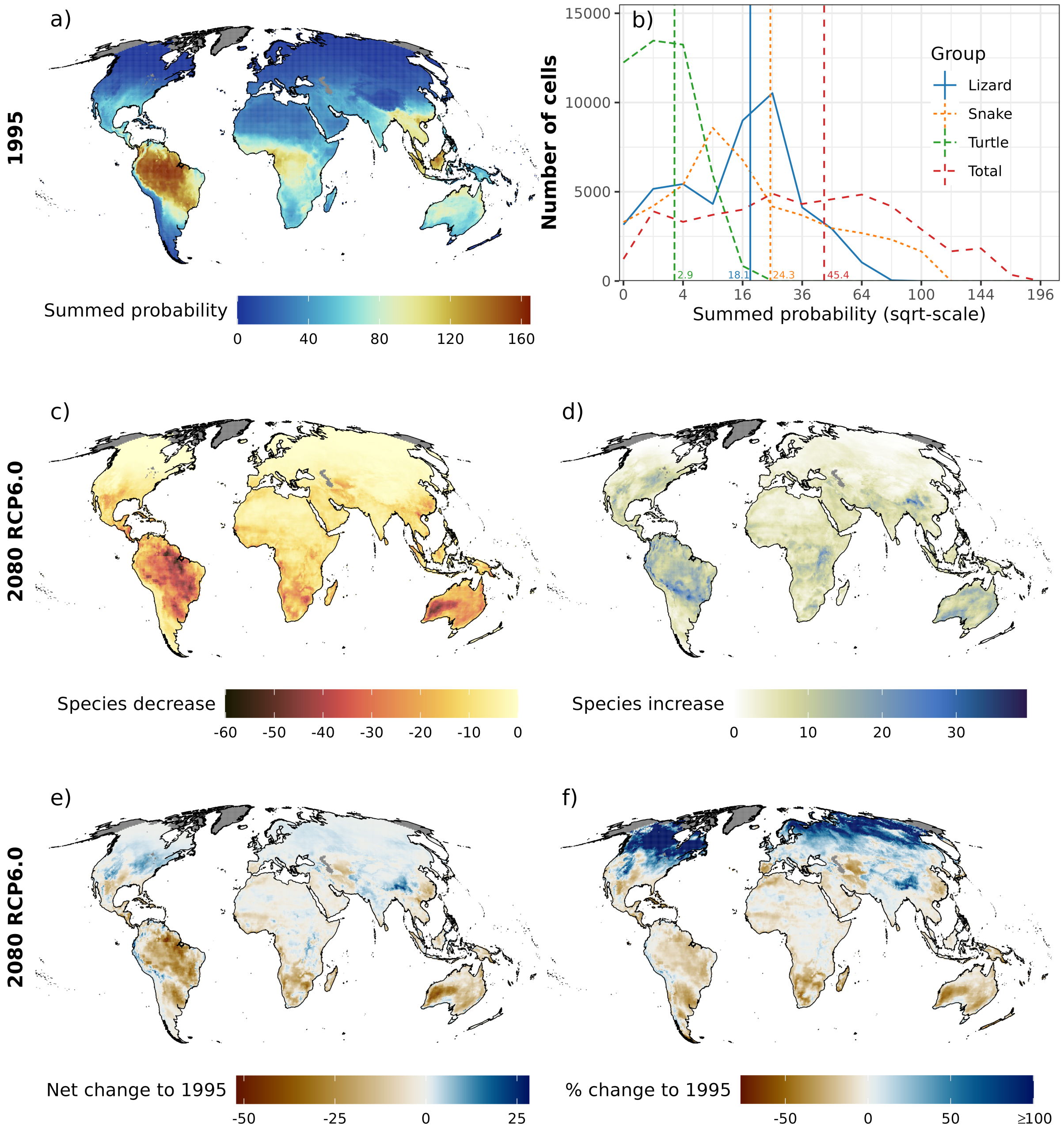


**Figure S4.18.** a) Map of projected global terrestrial reptile summed probability (1995), b) frequency of summed probability by taxonomic group (lizard, snake, turtle and total) with mean values highlighted by vertical line and c) increase, d) decrease, e) net change and f) relative change (%) in reptile summed probability for all modelled reptile species (N = 6296) for the year 2080 under a medium representative concentration pathway (RCP6.0) and a medium dispersal scenario (d/8). The results are presented as the ensemble mean, across the four global circulation models (GCMs) and two model algorithms (GAM & GBM) considered. All maps are based on 0.5° x 0.5° grid cells, which have been projected to Mollweide equal-area projection (EPSG:54009). Grey areas are regions for which no projections are available. Note that the colour scales differ among the individual panels.

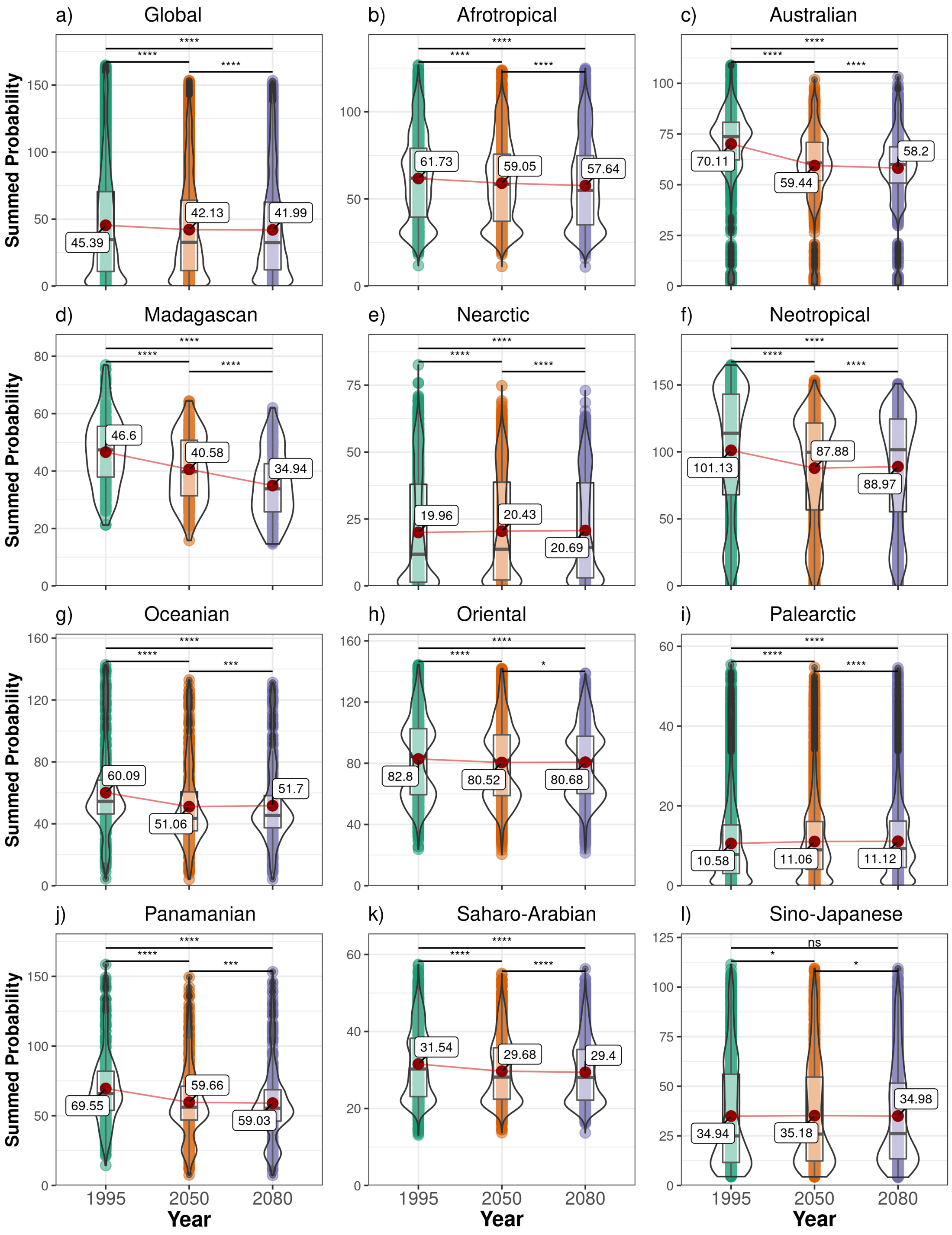


**Figure S4.19.** Terrestrial reptile summed probability across the globe and for each zoogeographic realm over time (1995, 2050, 2080) based on all modelled reptile species (N = 6296). The results are presented as the ensemble mean, across the four global circulation models (GCMs) and two model algorithms (GAM & GBM) considered, under a medium representative concentration pathway (RCP6.0) and a medium dispersal scenario (d/8). The statistical difference between years was tested using a paired Student's t-test with Holm correction ($p < 0.05$ = *, $p < 0.01$ = **, $p < 0.001$ = ***, $p < 0.0001$ = ****). Plots show mean (red point & label), median (black horizontal line), 25th to 75th percentiles (box), entire range of data (violin & data points) and density of values (width of violin).

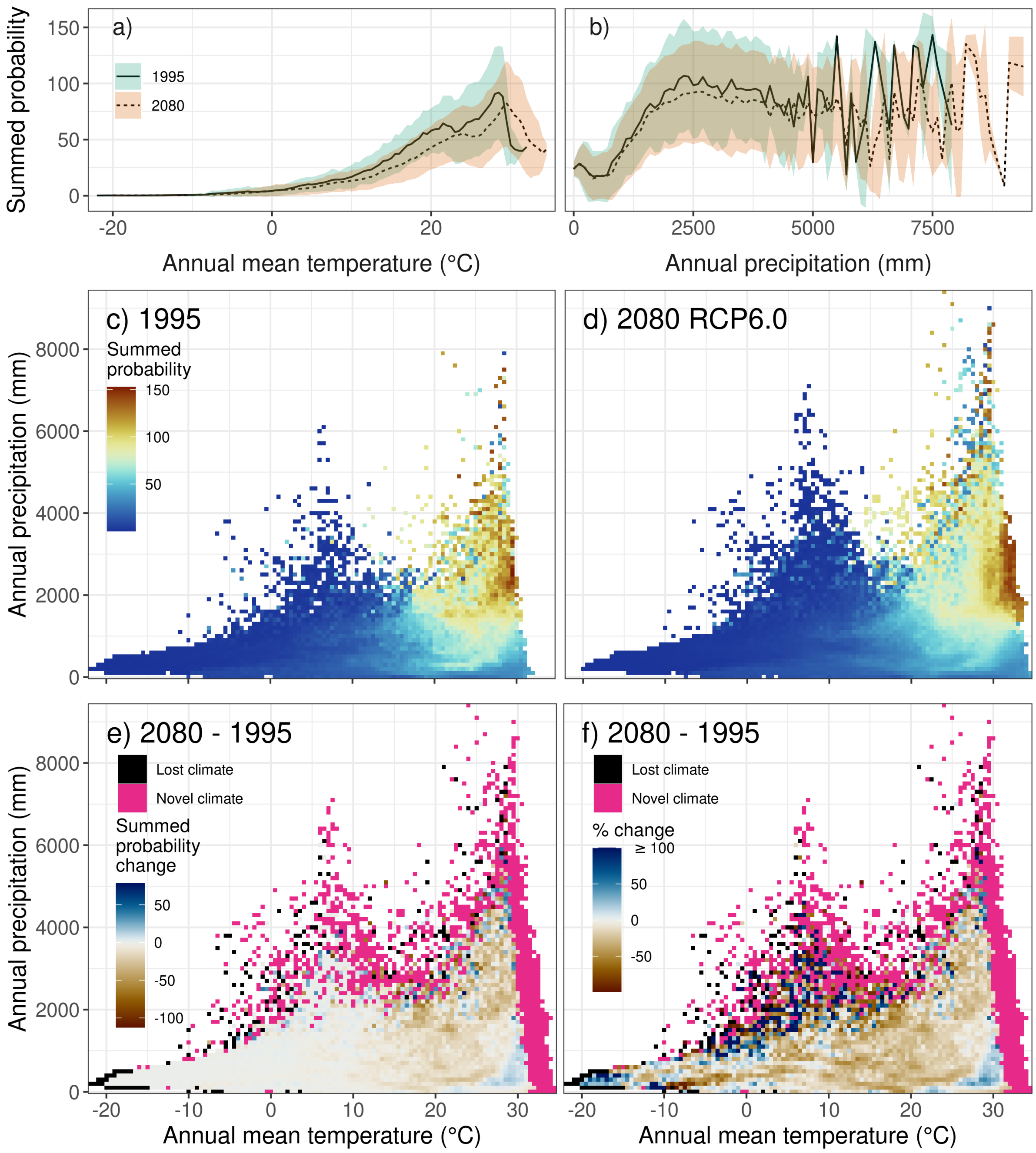


**Figure S4.20** Univariate relationship of current (1995) and future (2080 RCP6.0) reptile summed probability with a) temperature, b) precipitation and the bivariate relationship of temperature and precipitation with reptile summed probability for c) 1995 and d) 2080 RCP6.0 and the respective e) net change and f) relative change (%) under a medium dispersal scenario (d/8). Heat maps and lines show the mean, while ribbons indicate the standard deviation in variance across space, global circulation models (GCMs) and the two model algorithms.

#### Appendix S5 Variation across dispersal distances

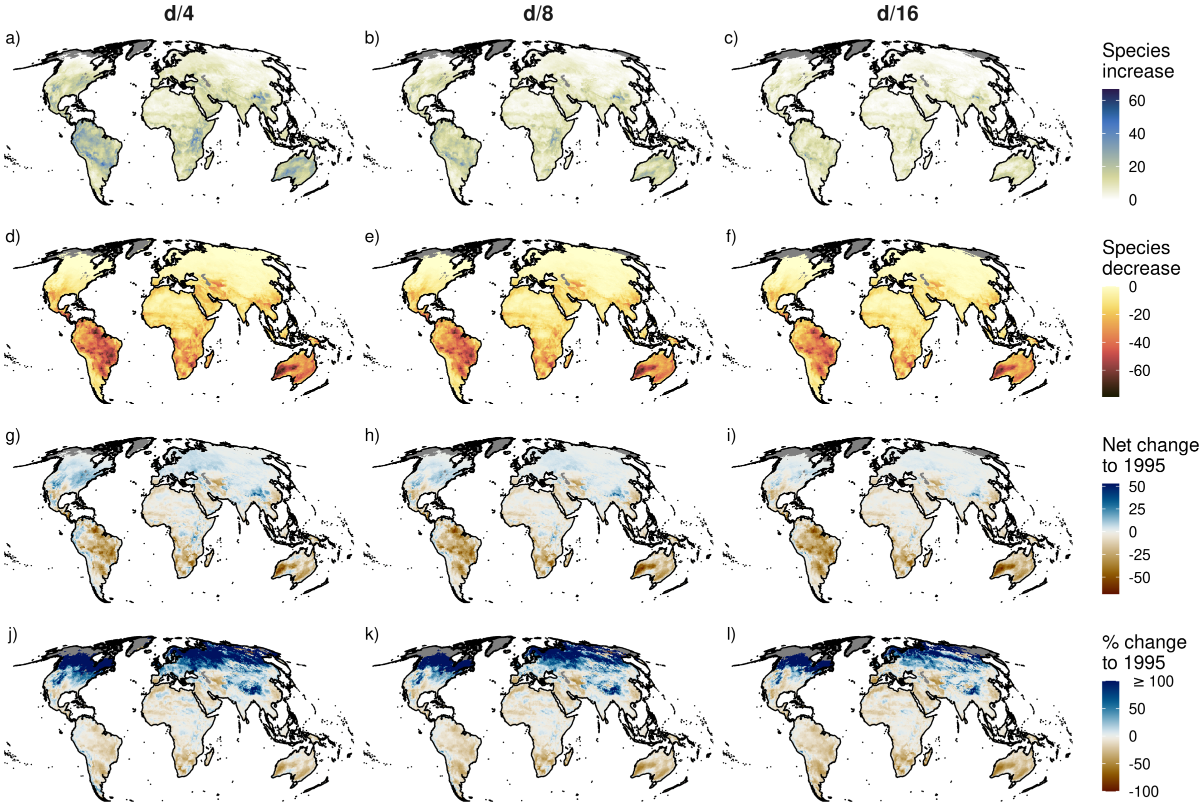


**Figure S5.21.** Influence of dispersal scenario (d/4, d/8, d/16) on potential change in species richness (increase, decrease, net change, relative change (%)) for all modelled reptile species (N = 6296). The results are presented as the ensemble mean, across the four global circulation models (GCMs) and two model algorithms (GAM & GBM) considered, for the year 2080 under a medium representative concentration pathway (RCP6.0). All maps are based on 0.5° x 0.5° grid cells, which have been projected to Mollweide equal-area projection (EPSG:54009). Grey areas are regions for which no projections are available.

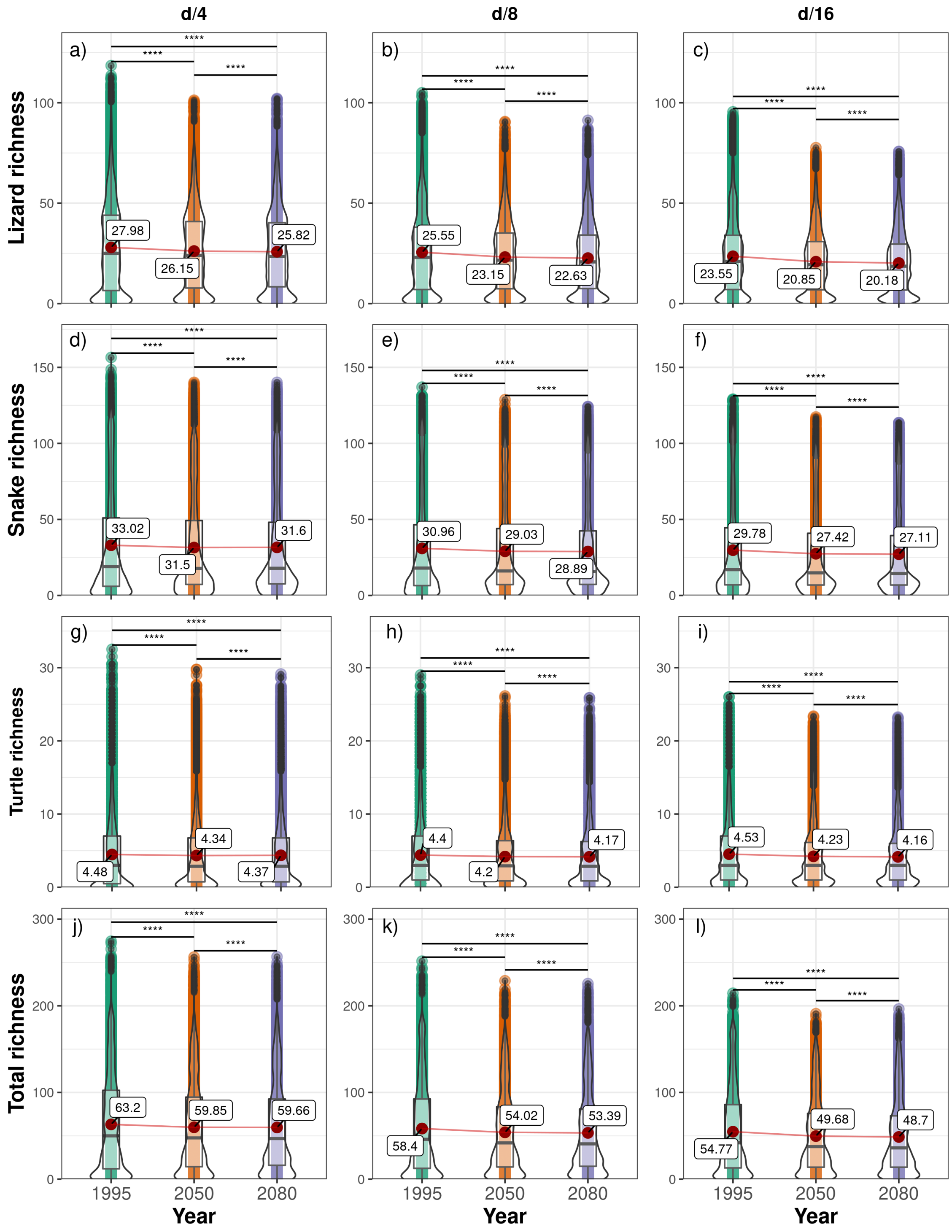


**Figure S5.22.** Variation in global terrestrial reptile richness across taxa (lizard, snake, turtle, total) and dispersal scenarios (d/4, d/8, d/16) over time (1995, 2050, 2080). The results are presented as the ensemble mean, across the four global circulation models (GCMs) and two model algorithms (GAM & GBM) considered, under a medium representative concentration pathway (RCP6.0). The statistical difference between years was tested using a paired Student's t-test with Holm correction ($p < 0.05$ = *, $p < 0.01$ = **, $p < 0.001$ = ***, $p < 0.0001$ = ****). Plots show mean (red point & label), median (black horizontal line), 25th to 75th percentiles (box), entire range of data (violin & data points) and density of values (width of violin).

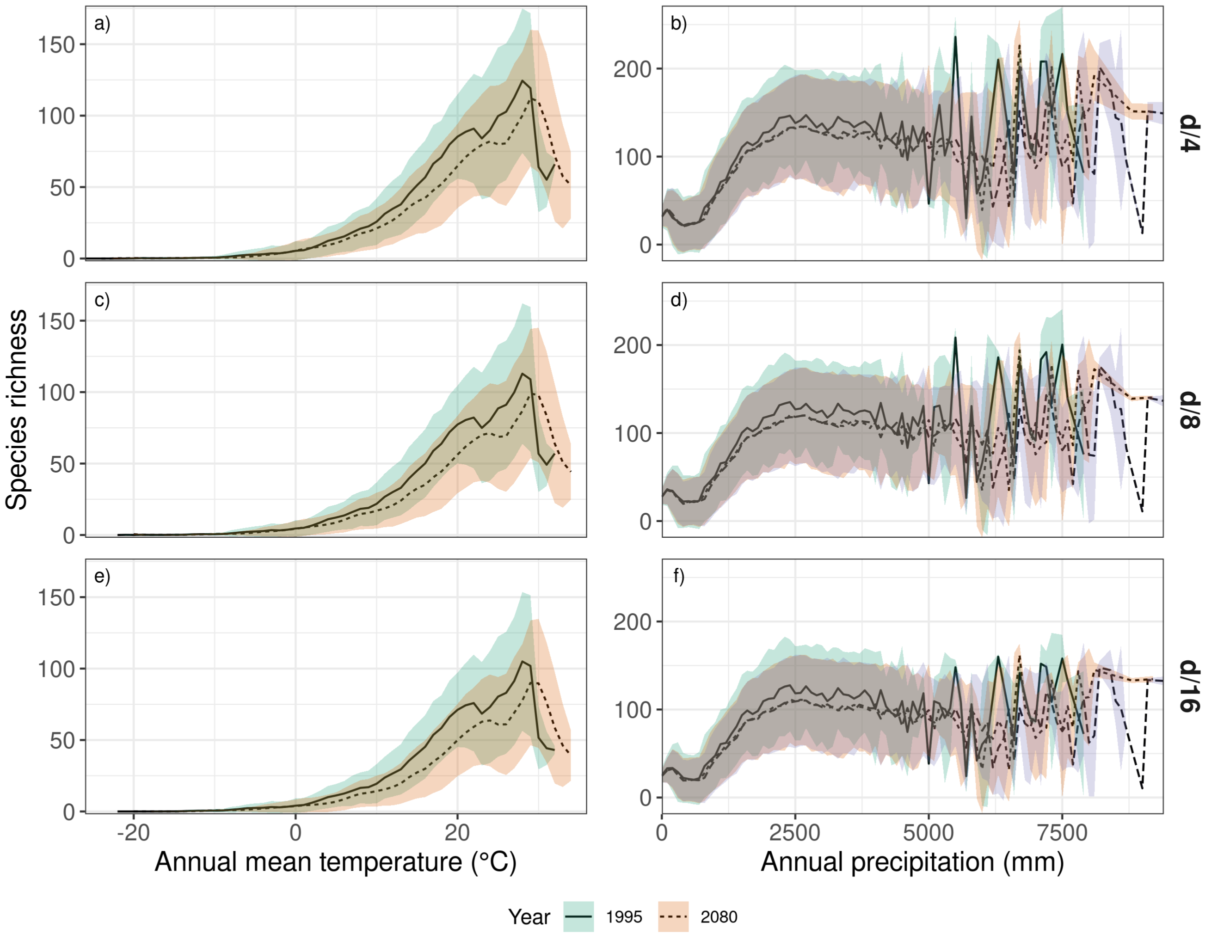


**Figure S5.23.** Univariate relationship of current (1995) and future (2080 RCP6.0) reptile species richness with a) temperature and b) precipitation under three dispersal scenarios (d/4, d/8, d/16) and a medium representative concentration pathway (RCP6.0). Lines display the mean and ribbons the standard deviation in variance across space, global circulation models (GCMs) and the two model algorithms (GAM& GBM).

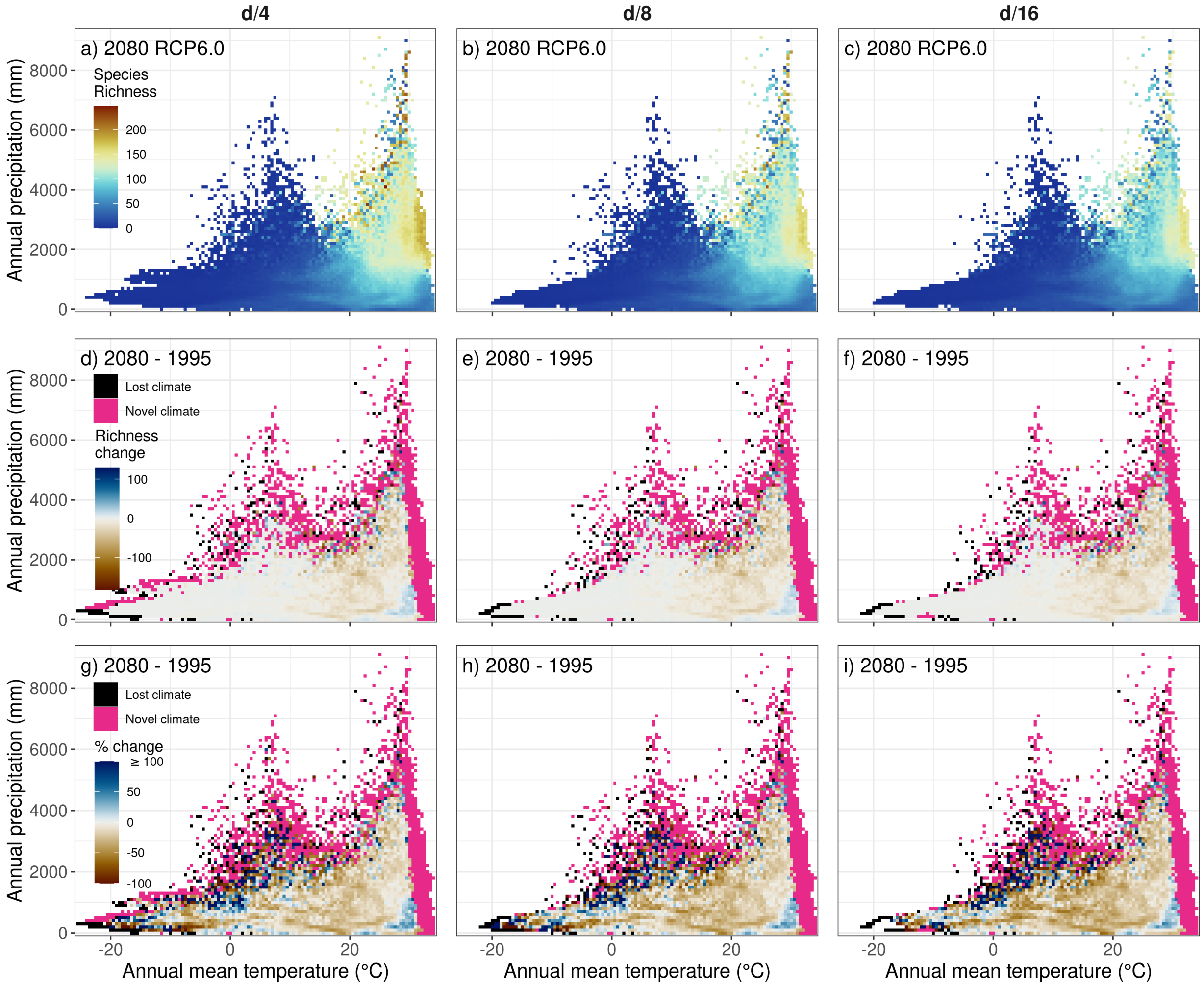


**Figure S5.24.** Bivariate relationship of temperature and precipitation with reptile species richness for different dispersal scenarios (d/4, d/8, d/16) for the year 2080 and a medium representative concentration pathway (RCP6.0). Heat maps show the mean across space, global circulation models (GCMs) and the two model algorithms (GAM & GBM).

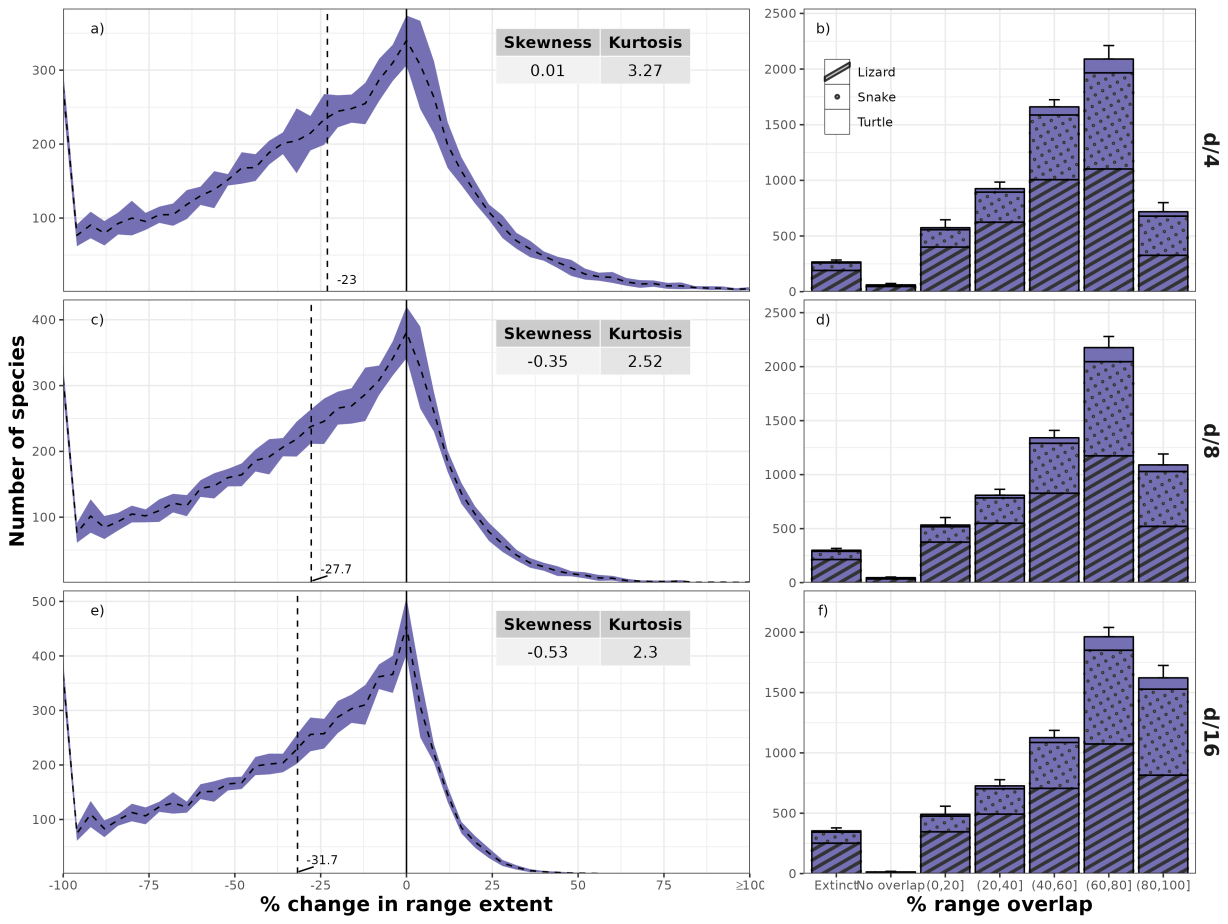


**Figure S5.25.** Frequency plots of the mean number of reptile species and their potential future change (%) in range extent and the mean number of reptile species per potential range overlap class (0-20, 20-40, 40-60, 60-80, 80-100) under different dispersal scenarios (d/4, d/8, d/16). Error margins/bars indicate the standard deviation across the four global circulation models (GCMs) and the two model algorithms (GAM & GBM) used. Both shown for 2080 under a medium representative concentration pathway (RCP6.0).

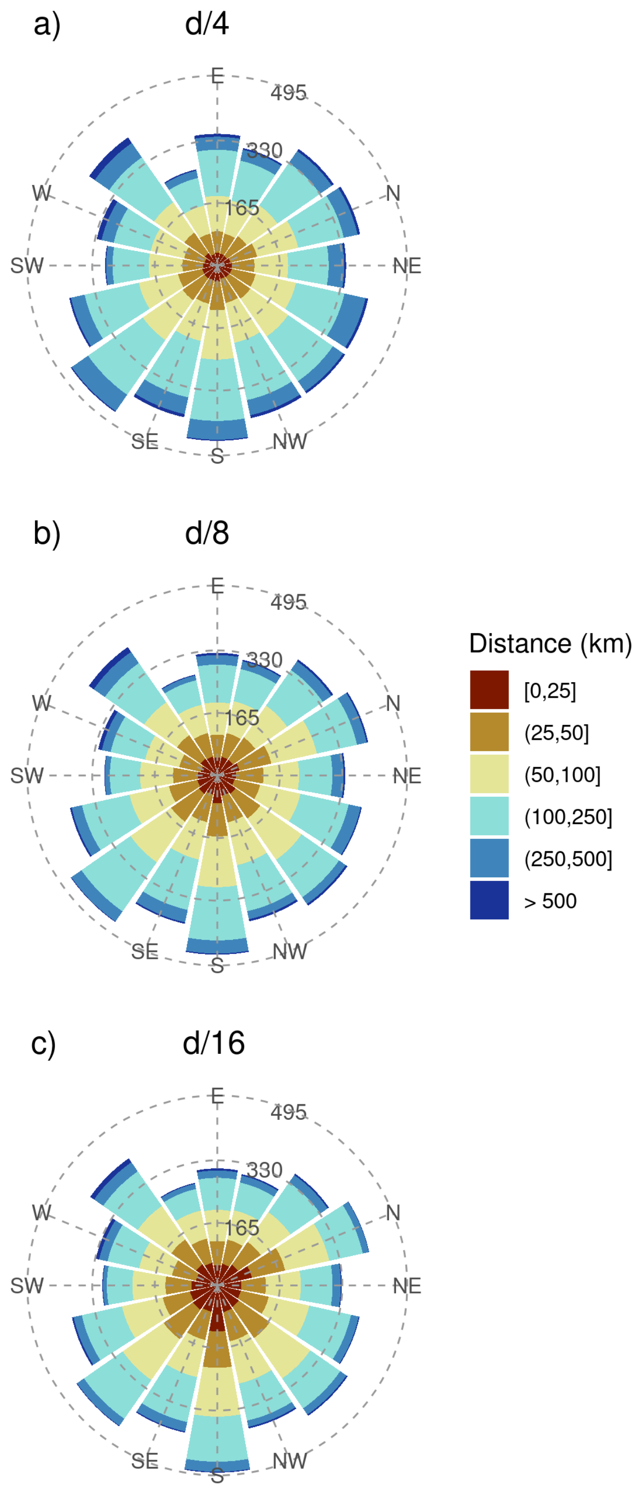


**Figure S5.26.** Cumulative direction and distance of potential range centroid changes under different dispersal scenarios (d/4, d/8, d/16). The results are presented as the ensemble mean, across the four global circulation models (GCMs) and two model algorithms (GAM & GBM) considered, for the year 2080 under a medium representative concentration pathway (RCP6.0).

#### Appendix S6 Variation across RCPs and years

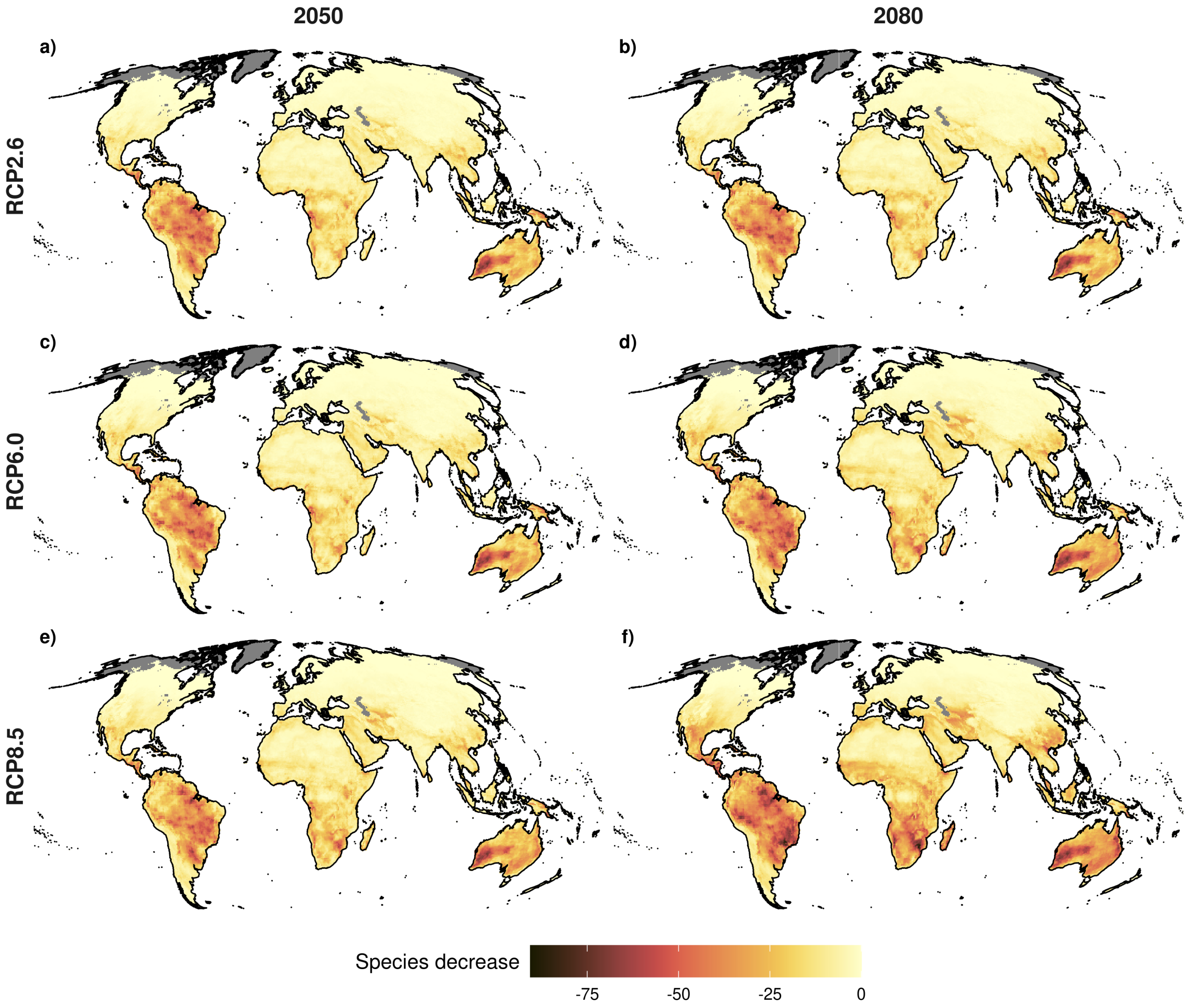


**Figure S6.27.** Influence of RCP (RCP2.6, RCP6.0, RCP8.5) and year (2050, 2080) on the potential decrease in species richness. The results are presented as the ensemble mean, across the four global circulation models (GCMs) and two model algorithms (GAM & GBM) considered, under a medium dispersal scenario (d/8). All maps are based on 0.5° x 0.5° grid cells, which have been projected to Mollweide equal-area projection (EPSG:54009). Grey areas are regions for which no projections are available.

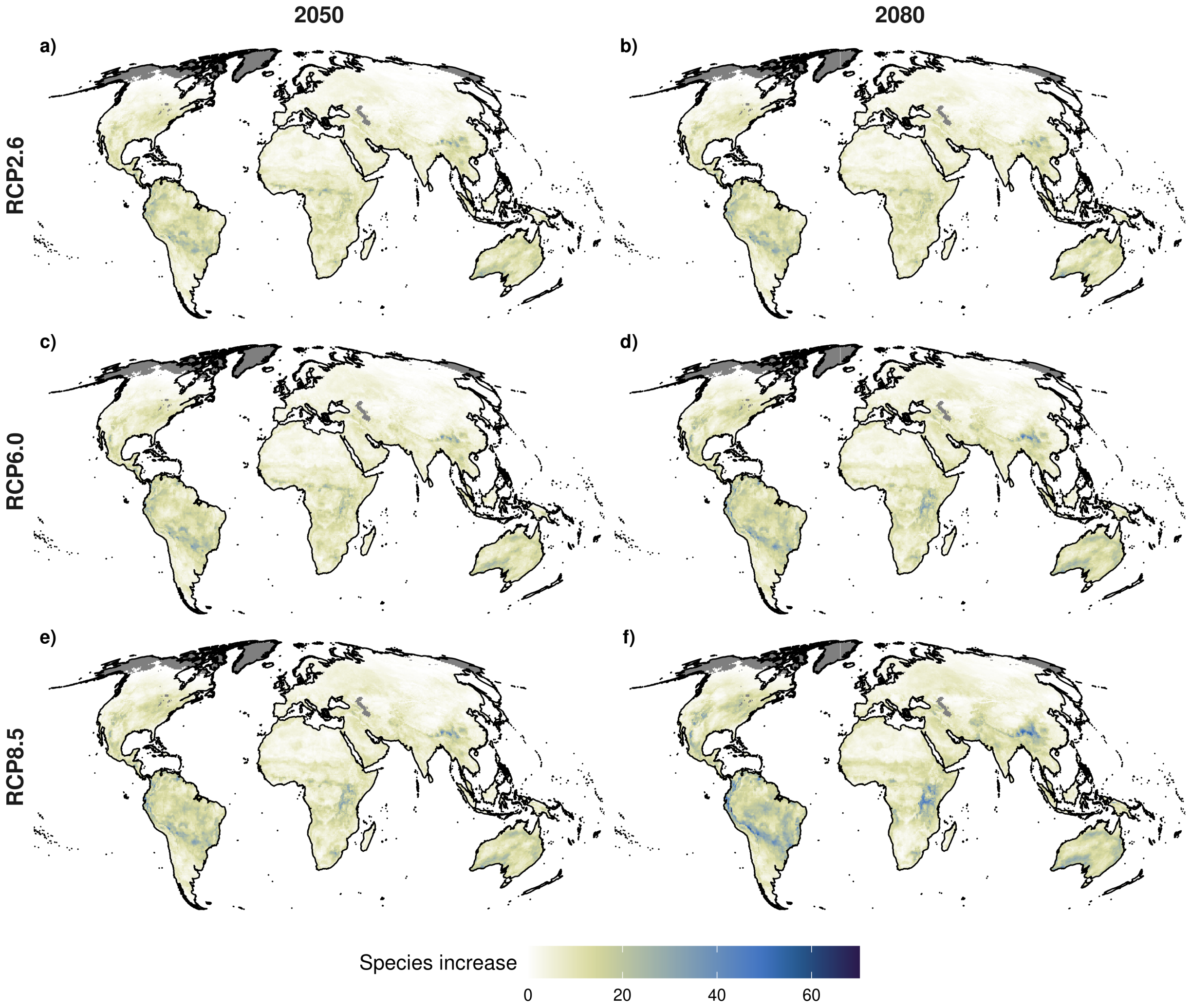


**Figure S6.28.** Influence of RCP (RCP2.6, RCP6.0, RCP8.5) and year (2050, 2080) on the potential increase in species richness. The results are presented as the ensemble mean, across the four global circulation models (GCMs) and two model algorithms (GAM & GBM) considered, under a medium dispersal scenario (d/8). All maps are based on 0.5° x 0.5° grid cells, which have been projected to Mollweide equal-area projection (EPSG:54009). Grey areas are regions for which no projections are available.

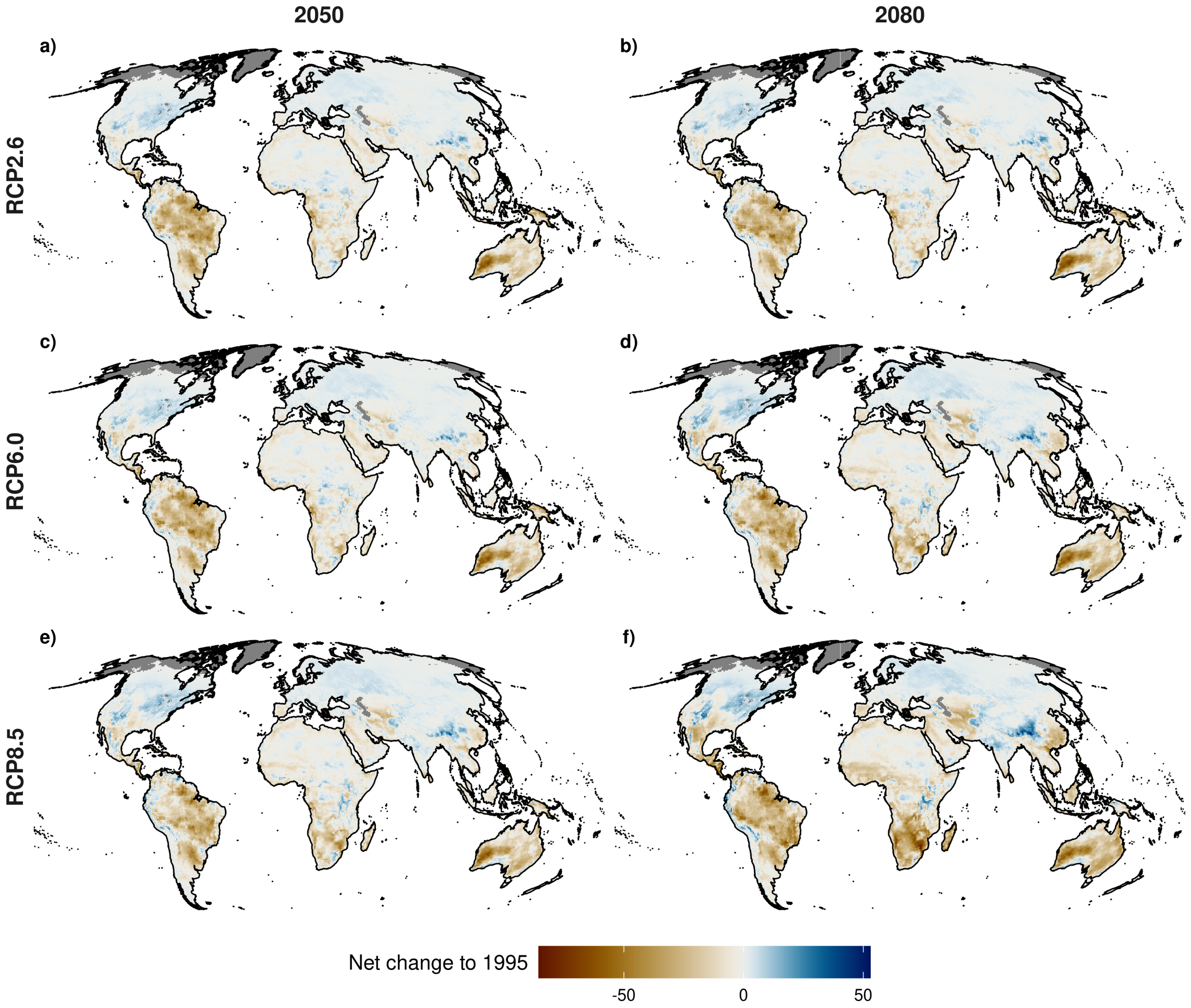


**Figure S6.29.** Influence of RCP (RCP2.6, RCP6.0, RCP8.5) and year (2050, 2080) on the potential net change in species richness. The results are presented as the ensemble mean, across the four global circulation models (GCMs) and two model algorithms (GAM & GBM) considered, under a medium dispersal scenario (d/8). All maps are based on 0.5° x 0.5° grid cells, which have been projected to Mollweide equal-area projection (EPSG:54009). Grey areas are regions for which no projections are available.

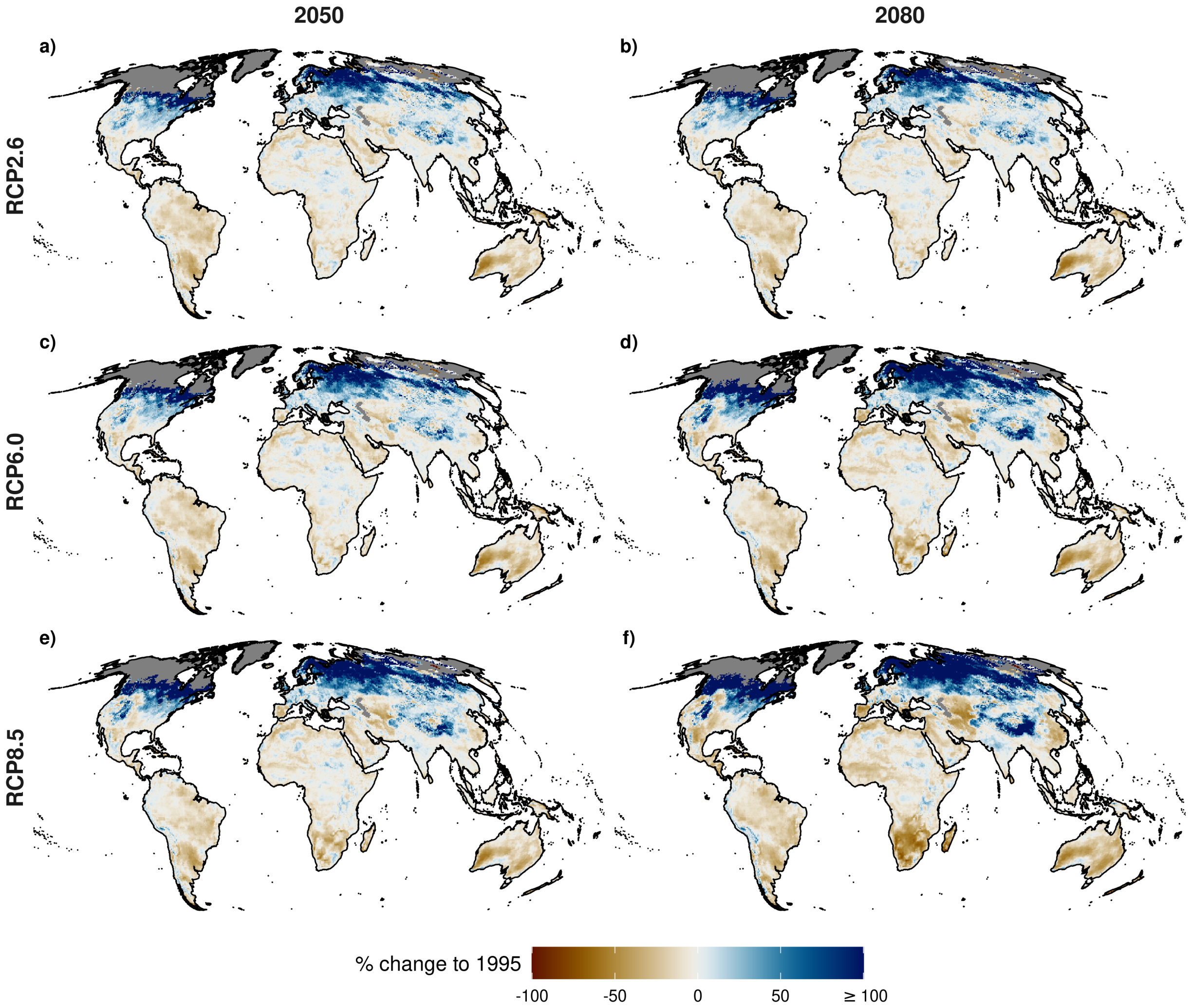


**Figure S6.30.** Influence of RCP (RCP2.6, RCP6.0, RCP8.5) and year (2050, 2080) on the potential relative change in species richness. The results are presented as the ensemble mean, across the four global circulation models (GCMs) and two model algorithms (GAM & GBM) considered, under a medium dispersal scenario (d/8). All maps are based on 0.5° x 0.5° grid cells, which have been projected to Mollweide equal-area projection (EPSG:54009). Grey areas are regions for which no projections are available.

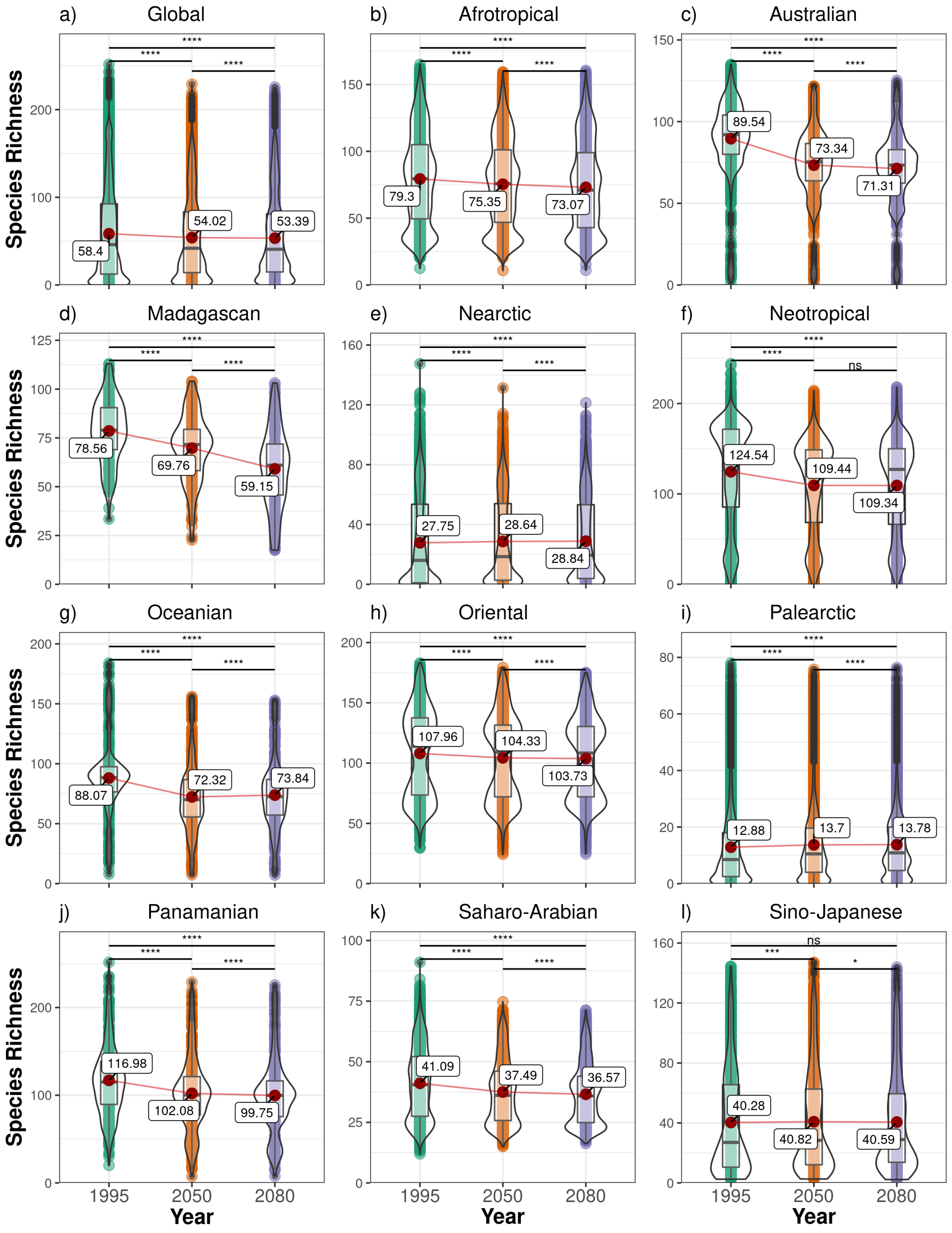


**Figure S6.31.** Terrestrial reptile species richness across the globe and for each zoogeographic realm (Afrotropical, Australian, Madagascan, Nearctic, Neotropical, Oceanian, Oriental, Palearctic, Panamanian, Saharo-Arabian & Sino-Japanese) over time (1995, 2050, 2080) based on all modelled reptile species (N = 6296). The results are presented as the ensemble mean, across the four global circulation models (GCMs) and two model algorithms (GAM & GBM) considered, under a medium representative concentration pathway (RCP6.0) and a medium dispersal scenario (d/8). The statistical difference between years was tested using a paired Student's t-test with Holm correction ($p < 0.05$ = *, $p < 0.01$ = **, $p < 0.001$ = ***, $p < 0.0001$ = ****). Plots show mean (red point & label), median (black horizontal line), 25th to 75th percentiles (box), entire range of data (violin & data points) and density of values (width of violin).

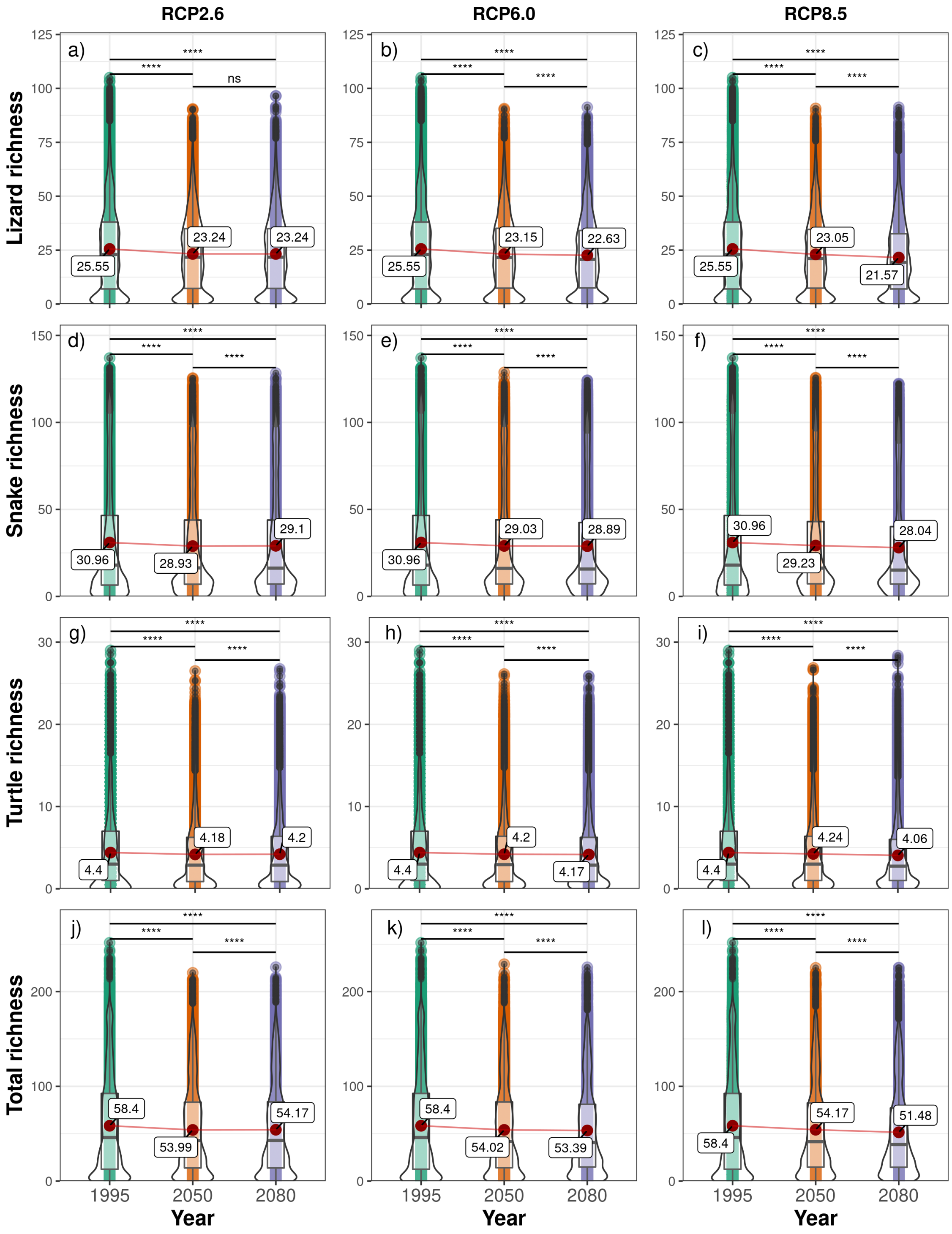


**Figure S6.32.** Variation in global terrestrial reptile richness across taxa and representative concentration pathways (RCP2.6, RCP6.0, RCP8.5) over time (1995, 2050, 2080). The results are presented as the ensemble mean, across the four global circulation models (GCMs) and two model algorithms (GAM & GBM) considered, under a medium dispersal scenario (d/8). The statistical difference between years was tested using a paired Student's t-test with Holm correction ($p < 0.05$ = *, $p < 0.01$ = **, $p < 0.001$ = ***, $p < 0.0001$ = ****). Plots show mean (red point & label), median (black horizontal line), 25th to 75th percentiles (box), entire range of data (violin & data points) and density of values (width of violin).

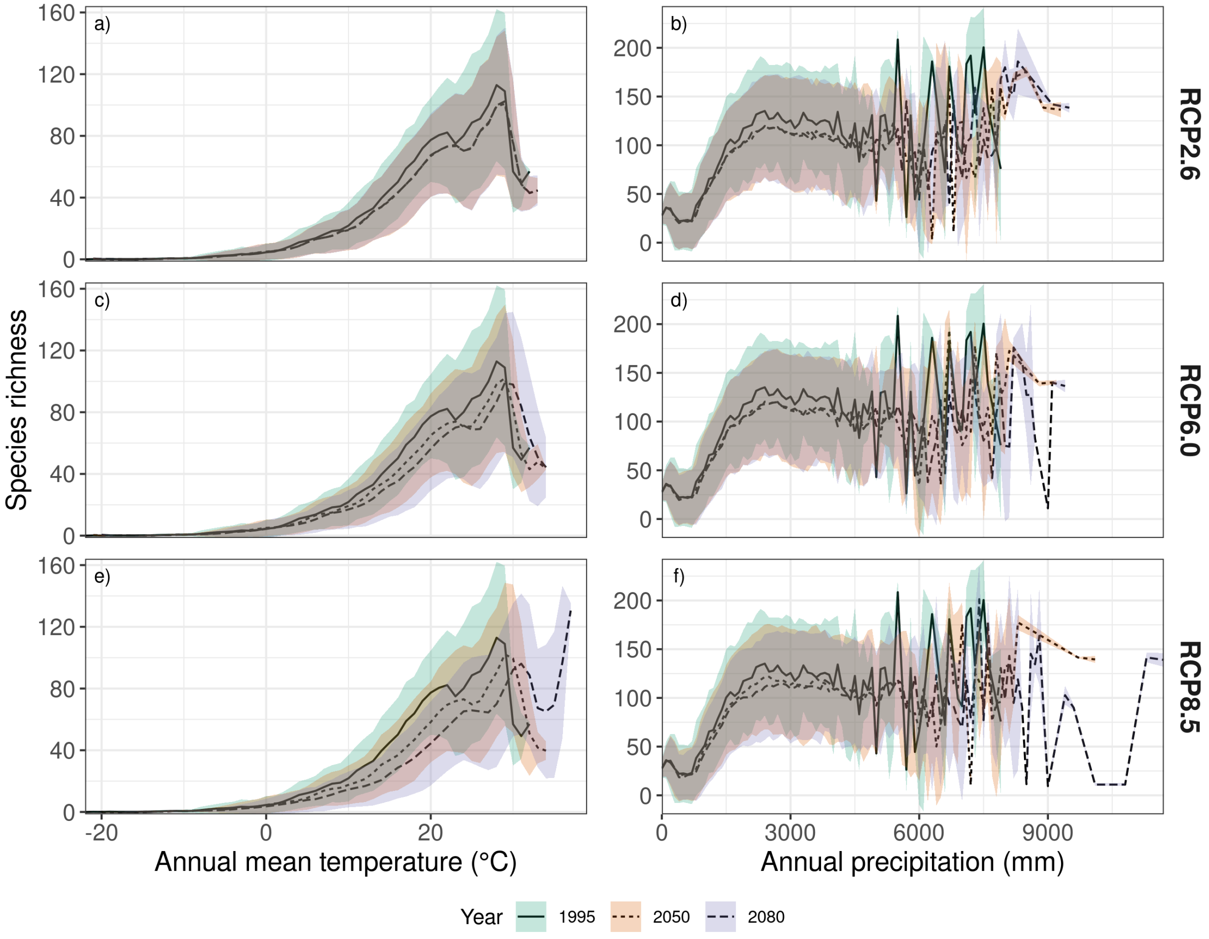


**Figure S6.33.** Univariate relationship of current (1995) and future reptile species richness (2050, 2080) with a) temperature, b) precipitation under different representative concentration pathways (RCP2.6, RCP6.0, RCP8.5) under a medium dispersal scenario (d/8). Lines display the mean and ribbons the standard deviation in variance across space, global circulation models (GCMs) and the two model algorithms (GAM & GBM).

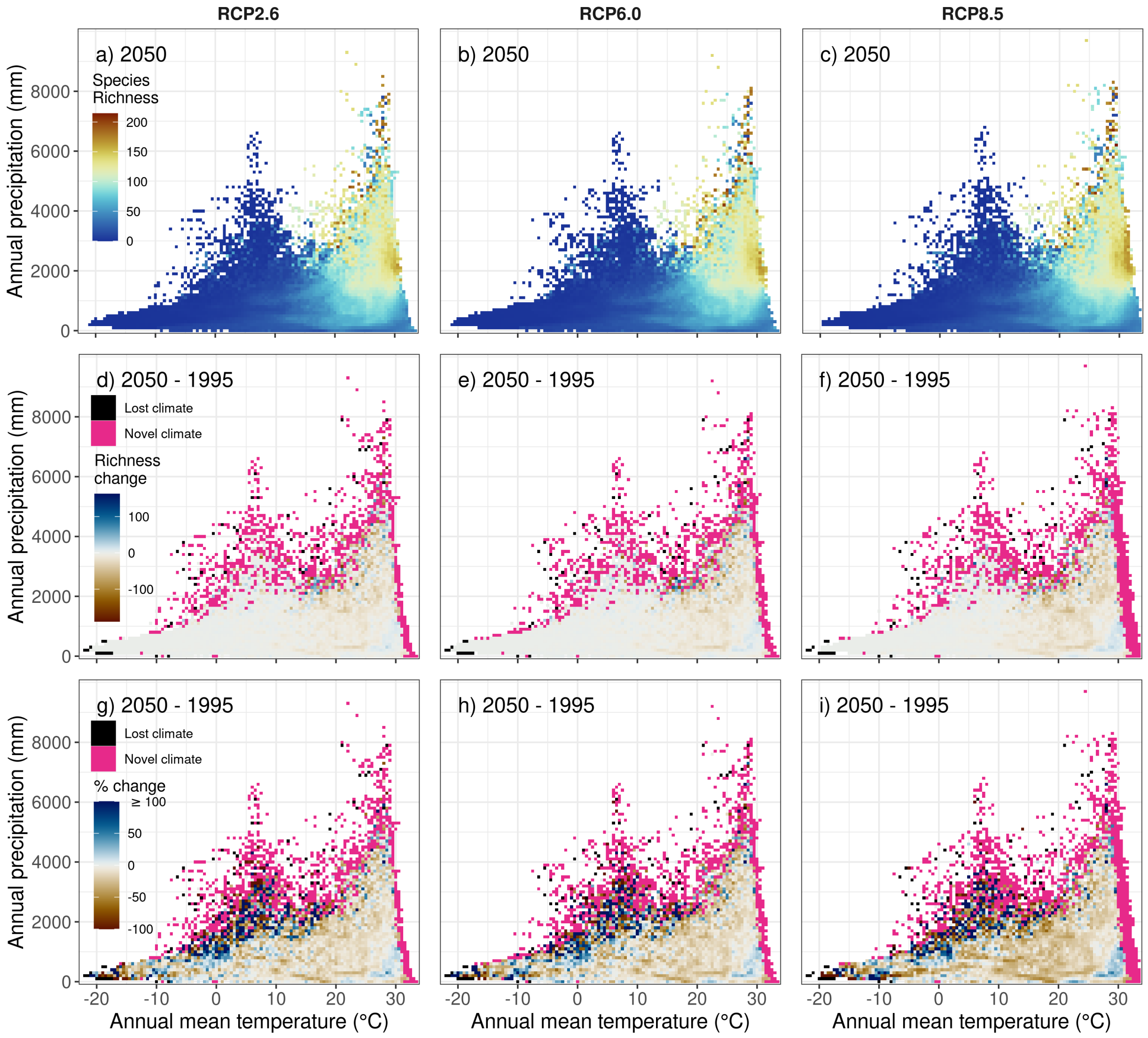


**Figure S6.34.** Bivariate relationship of temperature and precipitation with reptile species richness for the year 2050 and different representative concentration pathways (RCP2.6, RCP6.0, RCP8.5) under a medium dispersal scenario (d/8). Heat maps show the mean across space, global circulation models (GCMs) and the two model algorithms (GAM & GBM).

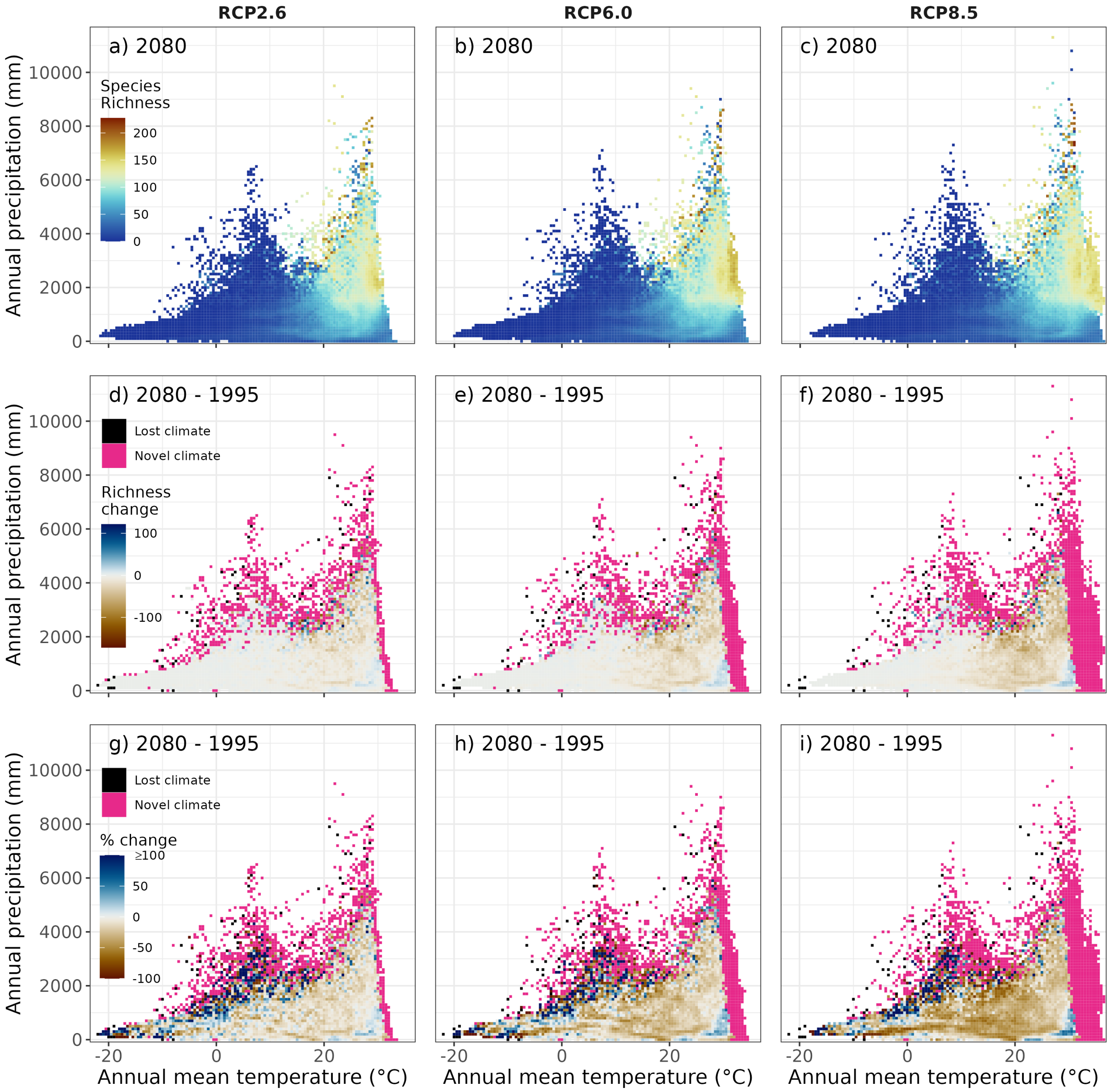


**Figure S6.35.** Bivariate relationship of temperature and precipitation with reptile species richness for the year 2080 and different representative concentration pathways (RCP2.6, RCP6.0, RCP8.5) under a medium dispersal scenario (d/8). Heat maps show the mean across space, global circulation models (GCMs) and the two model algorithms (GAM & GBM).

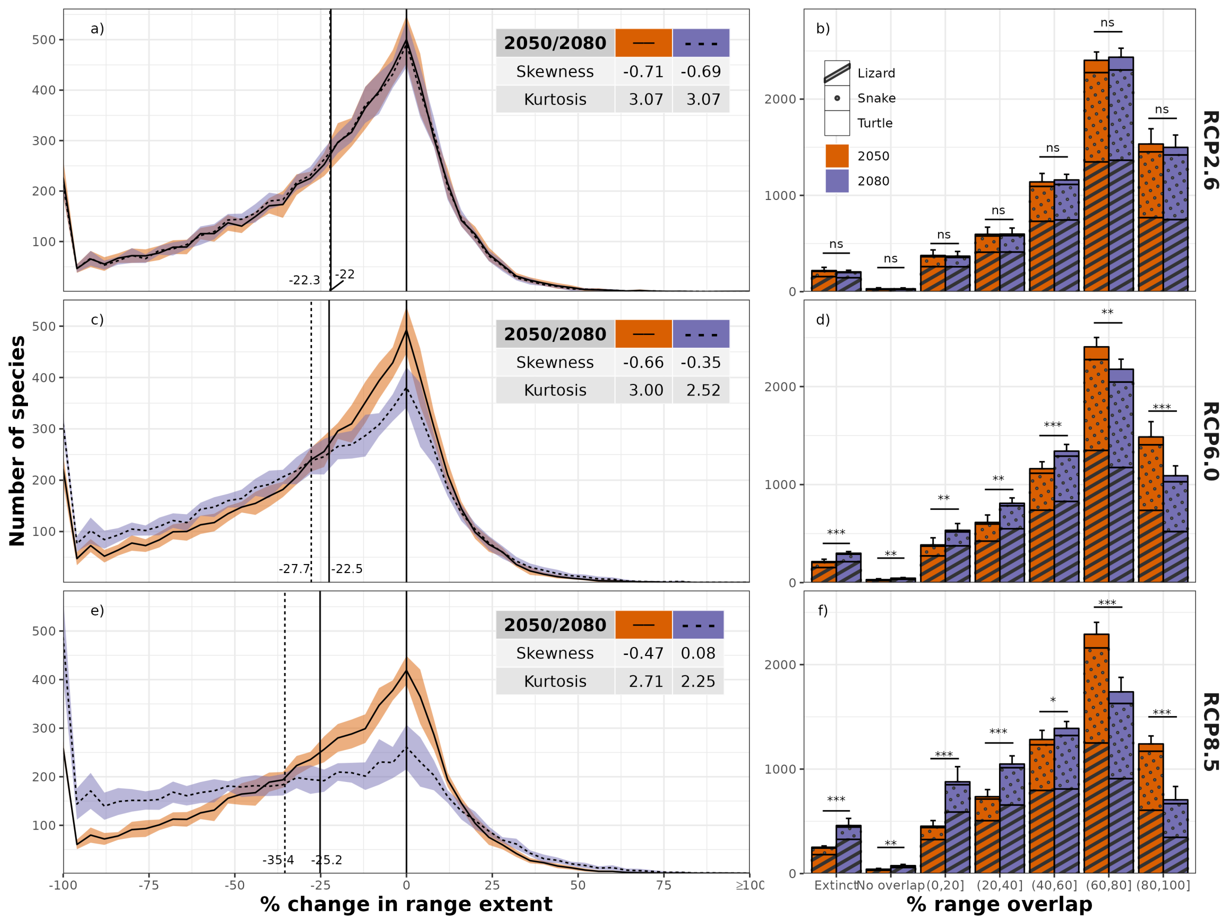


**Figure S6.36.** Frequency plots of the mean number of reptile species and their potential future change (%) in range extent and the mean number of reptile species per potential range overlap class (0-20, 20-40, 40-60, 60-80, 80-100) under different representative concentration pathways (RCP2.6, RCP6.0, RCP8.5). Error margins/bars indicate the standard deviation across the four global circulation models (GCMs) and the two model algorithms (GAM & GBM) used. Both shown for two time periods (2050, 2080) and a medium dispersal scenario (d/8).

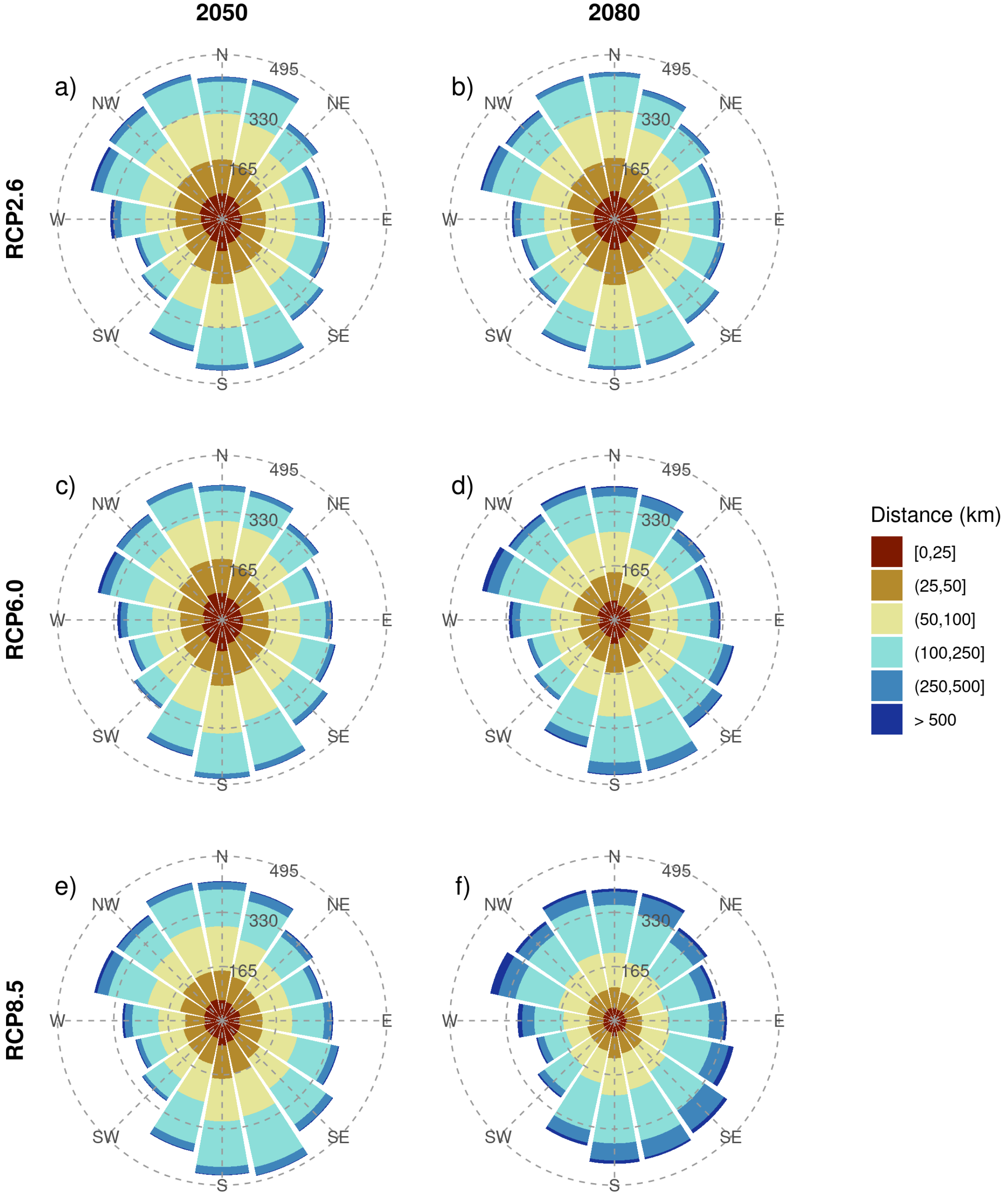


**Figure S6.37.** Cumulative direction and distance of potential range centroid changes under different representative concentration pathways (RCP2.6, RCP6.0, RCP8.5) and two time periods (2050, 2080). The results are presented as the ensemble mean, across the four global circulation models (GCMs) and the two model algorithms (GAM & GBM) considered, under a medium dispersal scenario (d/8).

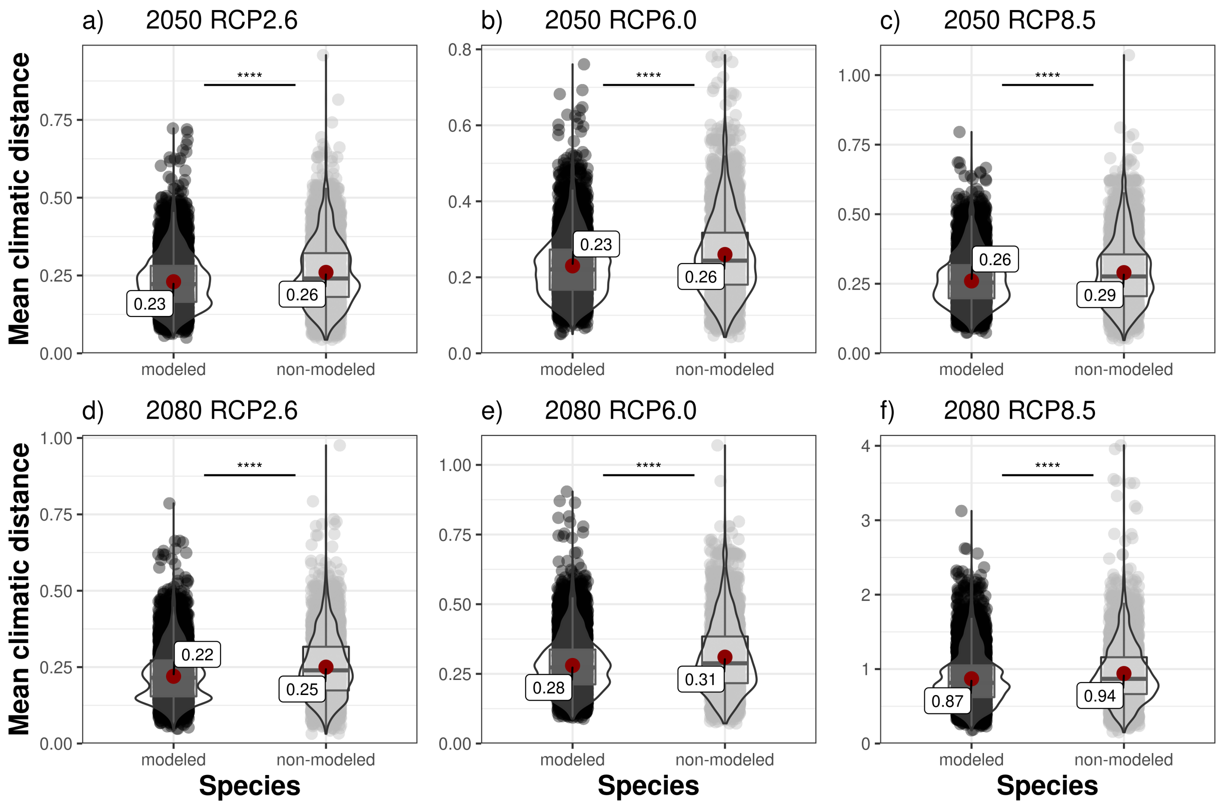


**Figure S6.38.** Mean climatic distance for modelled and non-modelled species, shown for different time periods (2050 & 2080) and representative concentration pathways (RCP2.6, RCP6.0, RCP8.5). The results are shown as the ensemble mean across the four global circulation models (GCMs). The statistical difference between modelled and non-modelled species was tested using a Student's t-test with Holm correction ($p < 0.05$ = *, $p < 0.01$ = **, $p < 0.001$ = ***, $p < 0.0001$ = ****). Plots show mean (red point & label), median (black horizontal line), 25th to 75th percentiles (box), entire range of data (violin & data points) and density of values (width of violin).
